# Cardio-respiratory interactions in interoceptive perception: The role of heartbeat-modulated cortical oscillations

**DOI:** 10.1101/2025.04.14.648690

**Authors:** Andrea Zaccaro, Francesca della Penna, Francesco Bubbico, Başak Bayram, Eleonora Parrotta, Mauro Gianni Perrucci, Marcello Costantini, Francesca Ferri

## Abstract

The cardiovascular and respiratory systems are anatomically and functionally integrated within the cardio-respiratory system. This close connection suggests that breathing continuously shapes cardiac interoceptive perception. Previously, we demonstrated cardio-respiratory interoceptive interactions in the heartbeat-evoked potential, a neural marker of cortical processing of cardiac signals. Specifically, we observed enhanced late heartbeat-evoked potential positivity and greater interoceptive accuracy during exhalation compared to inhalation in participants engaged in cardiac interoceptive tasks. Here, we extended these findings to the time-frequency domain by reanalysing our previous dataset. We investigated heartbeat-modulated cortical oscillations, examining power, inter-trial coherence, and functional connectivity across the respiratory cycle at rest, during a cardiac interoceptive task (heartbeat counting), and an exteroceptive control task (cardiac-tone counting). Results revealed that during the heartbeat counting task, late heartbeat-related power, inter-trial coherence, and functional connectivity increased during exhalation compared to inhalation, particularly in the alpha and theta frequency bands. These effects were primarily localized to right fronto-centro-parietal electrodes. Furthermore, we identified interactive relationships between heartbeat-evoked potential and heartbeat-modulated cortical oscillations in the alpha band that predicted interoceptive accuracy. These relationships were independent of cardiac physiology and were absent in the exteroceptive task. We proposed a model of cardio-respiratory interactions within the framework of interoceptive predictive coding, suggesting that these interactions occur at multiple levels of the interoceptive hierarchy: peripheral, brainstem, and cortical. Our interpretation highlights the role of heartbeat-related alpha-band modulations in enhancing the precision-weighting of cardiac prediction errors, thereby facilitating attentional allocation to interoceptive signals and the suppression of task-irrelevant distractors, particularly during exhalation.

## 1. Introduction

Beneath the threshold of awareness, the brain continuously processes signals from the body’s internal systems, inferring their states and modulating their activity to maintain vital functions essential for survival and psychophysical health, a process referred to as interoception (Berntson & Khalsa, 2021; Bonaz et al., 2021; Candia-Rivera et al., 2024; Chen et al., 2021; Craig, 2003; Khalsa et al., 2018; Nord & Garfinkel, 2022; Saltafossi, Heck, et al., 2025; Tallon-Baudry, 2023). Among these, the cardiovascular and respiratory systems, anatomically and functionally integrated within the cardio-respiratory system (Elstad et al., 2018; Parviainen et al., 2022), are regulated by the autonomic nervous system to optimize oxygen uptake and distribution to bodily tissues. This interplay is exemplified by respiratory sinus arrhythmia (RSA), a phenomenon in which heart rate (HR) fluctuates in synchrony with the phases of respiration (Berntson et al., 1993; Brecher and Hubay, 1955; Yasuma and Hayano, 2004). Furthermore, the two systems share bottom-up interoceptive pathways via the vagus nerve that converge in brain regions such as the insular, somatosensory, and cingulate cortices (Chan et al., 2024; Engelen et al., 2023; Lovelace et al., 2024; Park & Blanke, 2019; Weng et al., 2021). This close connection suggests that respiration continuously modulates the perception of cardiac signals, making it inherently challenging to dissociate the two, a perspective often overlooked in interoception research.

We have previously highlighted such cardio-respiratory interoceptive interactions as reflected in the heartbeat-evoked potential (HEP), an event-related potential time-locked to systole that indicates cortical processing of cardiac signals (Coll et al., 2021; Schandry and Weitkunat, 1986). In two independent samples of healthy individuals, we found increased late HEP positivity (emerging after 350 msec from the R-peak) during exhalation compared to inhalation while participants engaged in two cardiac interoceptive tasks: the heartbeat detection task (HBD; Zaccaro et al., 2022) and the heartbeat counting task (HCT; Zaccaro et al., 2024). In the HBD, greater HEP amplitudes during exhalation compared to inhalation were positively correlated with greater interoceptive accuracy (Zaccaro et al., 2022). In the HCT, late HEP enhancements relative to the C-TCT were predominantly driven by exhalation, with no significant contribution from inhalation (Zaccaro et al., 2024). Notably, these effects were absent while participants engaged in tasks involving attention to external tones (the exteroceptive condition of the HBD and the cardiac-tone counting task, C-TCT) or during respiratory interoception (the breath counting task). Furthermore, these findings could not be explained by differences in respiratory rate, heart rate, or heart rate variability across conditions.

Together, these results supported the hypothesis that exhalation may enhance cardiac interoception compared to inhalation (Molle & Coste, 2022). They were also consistent with evidence that breathing dynamically shapes neural processing in different domains. The respiratory cycle interacts with both resting (Herrero et al., 2018; Kluger et al., 2023; Kluger & Gross, 2021) and task-related brain activity, modulating perceptual (Flexman et al., 1974; Gallego et al., 1991; Goheen et al., 2024; Grund et al., 2022.; Harting et al., 2025; Johannknecht & Kayser, 2022; Kluger et al., 2021; Leupin & Britz, 2024; Mizuhara et al., 2024, 2025; Mizuhara & Nittono, 2022; Münch et al., 2019; Saltafossi, Zaccaro, et al., 2025; Stetza et al., 2024), cognitive-emotional (Arshamian et al., 2018; Belli & Fischer, 2024; Brændholt et al., 2024; Huijbers et al., 2014; Molle et al., 2023; Nakamura et al., 2018; Perl et al., 2019; Waselius et al., 2019, 2022; Zelano et al., 2016; Mizuhara and Nittono, 2023; Schaefer et al., 2024), and motor functions (Engelen et al., 2024; Kluger & Gross, 2020; Park et al., 2020, 2022). However, despite its well-established role in exteroceptive functions, the precise influence of respiration on cardiac interoception remains unclear, particularly due to the strong physiological coupling between heartbeat dynamics and respiratory cycles, which introduces potential confounds and complicates interpretation.

A growing body of evidence suggests that, beyond time-domain HEPs, heartbeat-modulated cortical oscillations in the time-frequency domain may be used as more precise electrophysiological markers of cardiac interoception. A landmark electrocorticography study (Kern et al., 2013) identified heartbeat-related power fluctuations over somatosensory areas at rest, predominantly occurring in the low-frequency band (<12 Hz). Similarly, an intracranial EEG (iEEG) study (Park et al., 2018) identified heartbeat-related increases in inter-trial coherence (ITC) in the theta band within the insula and frontotemporal regions. These increases were observed both at rest and during a task designed to modulate bodily self-awareness (a full-body illusion paradigm), while no heartbeat-related power changes were detected in the same areas. In contrast, an iEEG study by Treves (2020) employing ICA-based denoising reported heartbeat-related increases in alpha power but found no changes in ITC. Several EEG studies have investigated heartbeat-related modulations in power, ITC, and functional connectivity (FC) across a range of conditions, including sleep stages (Lechinger et al., 2015; Simor et al., 2024), states of consciousness (Liuzzi et al., 2024), resting-state (Kim & Jeong, 2019), emotions (Kato et al., 2020), interoceptive metacognitive awareness (Canales-Johnson et al., 2015), interoceptive-exteroceptive processing (Fló et al., 2024; García-Cordero et al., 2017), Alzheimer’s disease (Castellanos et al., 2022), and meditation expertise (Jiang et al., 2020). Recently, Lee et al. (2024) employed a classification approach to highlight the role of heartbeat-related spectral perturbations in the alpha and theta bands in distinguishing interoceptive (heartbeat-focused) from exteroceptive (sound-focused) attentional states. Their results suggest that time-frequency measures may be less susceptible to physiological artifacts, such as the cardiac field artifact (CFA).

Taken together, these findings indicate that studying heartbeat-modulated cortical oscillations in the time-frequency domain may help to overcome the limitations of time-domain HEP analysis and may provide deeper insights into the neural mechanisms of cardiac interoception. In this study, we re-analysed our previously collected dataset (Zaccaro et al., 2024) comprising 28 healthy participants. Our investigation focused on heartbeat-related activity during the HCT and its exteroceptive control task (C-TCT), analysing separately heartbeats during inhalation and exhalation. We hypothesised that heartbeat-related power, ITC and FC would be differentially modulated by respiratory phase and task demands. Building on the findings of Lee et al. (Lee et al., 2024), we expected modulations in the theta (4-8 Hz) and alpha (8-13 Hz) frequency bands. To further explore the relationship between HEPs and time-frequency measures, we adopted a methodology from a study that reported a negative association between auditory-evoked P300 responses and alpha activity (Studenova et al., 2023). Finally, we applied linear mixed-effects models (LMEMs) to assess trial-level relationships in the HCT between HEP, power, and ITC across respiratory phases, as well as their associations with interoceptive accuracy and cardiac physiology. Based on prior research identifying alpha power as a marker of interoceptive attention shown to correlate with emotional arousal-related HEP modulations (Luft & Bhattacharya, 2015), interoceptive accuracy (Rominger et al., 2025; Villena-González et al., 2017), and to increase during heartbeat-focused tasks (Crivelli & Balconi, 2025; Kritzman et al., 2022; Mccraty et al., 2009; Villena-González et al., 2017), we expected to find positive associations between HEPs, heartbeat-modulated cortical oscillations in the alpha band, and interoceptive accuracy. We interpreted our findings within the interoceptive predictive coding framework, examining how respiration shapes predictive processes underlying cardiac perception and how cardio-respiratory interoceptive interactions may modulate the precision-weighting of cardiac prediction errors.

## 2. Results

### 2.1. Resting-state and task-level comparisons

Following an eyes-open resting-state condition, participants engaged in tasks requiring either cardiac interoceptive (HCT) or exteroceptive (C-TCT) attention. To determine whether heartbeat-modulated cortical oscillations varied across respiratory phases during the resting-state, we conducted full-scalp cluster-based permutation tests comparing inhalation to exhalation. Across all frequency bands, we found no significant differences in heartbeat-related power (theta: t_27_ = 2.085, p_clust_ = .097, d = .394; alpha: t_27_ = .013, p_clust_ = .989, d = .003; beta: t_27_ = .089, p_clust_ = .929, d = .017; gamma: t_27_ = 2.12, p_clust_ = .15, d = .401; Fig. S1A) or ITC (theta: t_27_ = -.915, p_clust_ = .369, d = .173; alpha: t_27_ = -2.381, p_clust_ = .096, d = .45; beta: t_27_ = 1.161, p_clust_ = .178, d = .219; gamma: t_27_ = 1.46, p_clust_ = .156, d = .276; Fig. S1B). Similarly, when not accounting for respiratory phases (i.e., averaging heartbeats occurring during both inhalation and exhalation), we observed no significant differences between HCT and C-TCT in heartbeat-related power (theta: t_27_ = .162, p_clust_ = .872, d = .031; alpha: t_27_ = -.163, p_clust_ = .872, d = .031; beta: t_27_ = .718, p_clust_ = .108, d = .136; gamma: t_27_ = .129, p_clust_ = .899, d = .024; Fig. S2A) or ITC (theta: t_27_ = -1.4, p_clust_ = .174, d = .264; alpha: t_27_ = - 2.28, p_clust_ = .215, d = .431; beta: t_27_ = 1.12, p_clust_ = .075, d = .212; gamma: t_27_ = 1.7, p_clust_ = .359, d = .322; Fig. S2B).

### 2.2 Heartbeat-related power is modulated by task and the respiratory phase

We performed a full-scalp cluster-based permutation test using heartbeat-related power as dependent variable and task (HCT vs. C-TCT), respiratory phase (inhalation vs. exhalation), and their interaction as within-participant factors. The test revealed a significant task × phase interaction in the alpha band from 300 to 600 msec over electrodes C4, C6, CP4, and CP6 (t_27_ = 3.165, p_clust_ = .037, d = .598; Fig. 1A). No significant interaction was observed in other frequency bands (theta: t_27_ = 1.933, p_clust_ = .14, d = .365; beta: t_27_ = 1.714, p_clust_ = .122, d = .324; gamma: t_27_ = -.525, p_clust_ = .276, d = .099). This interaction was explained by planned t-tests (see section 4.6), which showed significantly higher heartbeat-related alpha power during HCT from 391 to 600 msec over FCz, FC2, FC4, Cz, C2, C4, C6, CP2, CP4, and CP6 (t_27_ = 3.049, p_clust_ = .039, d = .576; Fig. 1B) during exhalation compared to inhalation. An increase in heartbeat-related theta power was also found from 300 to 469 msec after the R-peak over F6, FC4, FC6, Cz, C2, C4, C6, CP3, CP1, CPz, CP2, and P1 (t_27_ = 3.896, p_clust_ = .039, d = .736), while no significant differences were found for beta (t_27_ = .793, p_clust_ = .343, d = .149) or gamma (t_27_ = -1.375, p_clust_ = .264, d = .259) bands. Conversely, during C-TCT, we found no change in heartbeat-related power between exhalation and inhalation in any of the investigated frequency bands (theta: t_27_ = .261, p_clust_ = 1, d = .049; alpha: t_27_ = -.448, p_clust_ = 1, d = .085; beta: t_27_ = -2.661, p_clust_ = .09, d = .503; gamma: t_27_ = 1.659, p_clust_ = .052, d = .314; Fig. 1C).

**Figure 1.**
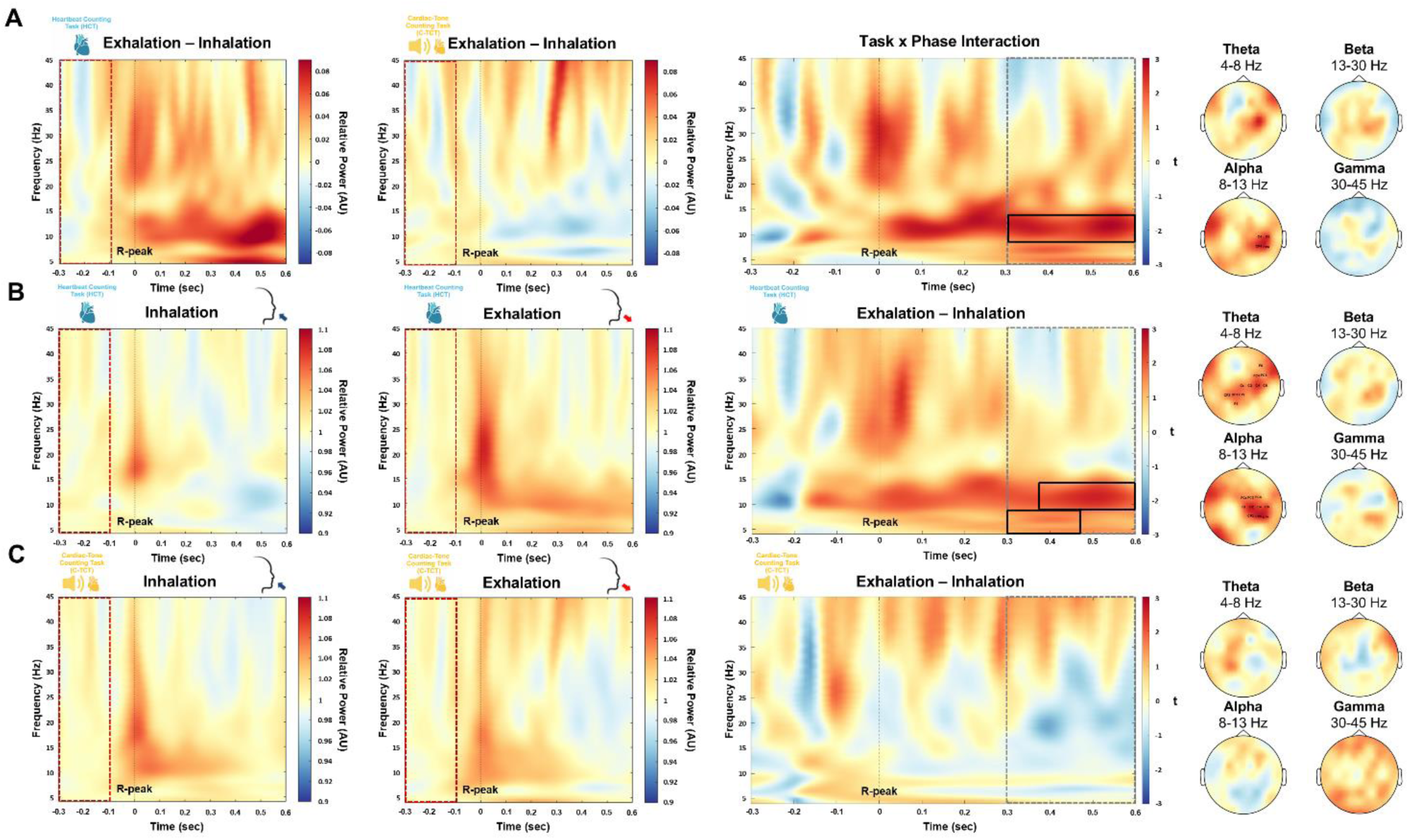
Heartbeat-related power changes across respiratory phases and tasks. (A) Grand-averaged heartbeat-related power changes between respiratory phases (exhalation vs. inhalation) during the heartbeat counting task (left), the cardiac-tone counting task (middle), and their task × phase interaction (right). The black rectangle highlights significant interactions (cluster-corrected). Topographical distributions depict the task × phase interaction, with labels indicating significant electrodes. (B) Grand-averaged heartbeat-related power in the heartbeat counting task during inhalation (left), exhalation (middle), and the exhalation-minus-inhalation difference (right). The black rectangle marks significant differences between exhalation and inhalation (cluster-corrected). Topographical distributions illustrate significant differences, with labels indicating significant electrodes. (C) Grand-averaged heartbeat-related power in the cardiac-tone counting task during inhalation (left), exhalation (middle), and the exhalation-minus-inhalation difference (right). The red dotted area represents the baseline window (-300 to -100 msec relative to R-peak onset). The grey dotted area marks the temporal window of interest for statistical analysis (300-600 msec after the R-peak). Abbreviations: AU = arbitrary unit.

To ensure that changes in alpha and theta band were not related to CFA, we tested the task × phase interaction from 300 to 600 msec over the time-frequency decomposed ECG signal, finding null results (theta: t_27_ = -1.174, p_FDR_ = .385, d = .222; alpha: t_27_ = -.428, p_FDR_ = .835, d = .081; Fig. S3).

### 2.3 Heartbeat-related inter-trial coherence is modulated by task and the respiratory phase

We performed a full-scalp cluster-based permutation test using heartbeat-related ITC as dependent variable and task (HCT vs. C-TCT), respiratory phase (inhalation vs. exhalation), and their interaction as within-participant factors. No significant task × phase interaction was observed for ITC in any of the frequency bands (theta: t_27_ = .129, p_clust_ = .132, d = .024; alpha: t_27_ = 2.104, p_clust_ = .072, d = .398; beta: t_27_ = .934, p_clust_ = .432, d = .177; gamma: t_27_ = 2.223, p_clust_ = .347, d = .421; Fig. 2A). When performing planned t-tests (see section 4.6) comparing heartbeat-related ITC between respiratory phases during HCT, we found a significant increase in alpha band during exhalation compared to inhalation from 422 to 559 msec after the R-peak over C2, C4, CP2, CP4, and CP6 (t_27_ = 3.238, p_clust_ = .023, d = .612). We also found a significant increase in gamma band ITC from 441 to 469 msec over C3, C5, CP3, CP5, and P5 (t_27_ = 3.825, p_clust_ = .038, d = .723; Fig. 2B). However, this effect was brief (28 msec) and localized over electrodes not typically associated with cardiac interoception (Coll et al., 2021). No significant ITC differences between respiratory phases were found for the other frequency bands (theta: t_27_ = .229, p_clust_ = 1, d = .043; beta: t_27_ = .629, p_clust_ = .345, d = .119). Similarly, no phase-related ITC changes were observed during the C-TCT across all frequency bands (theta: t_27_ = .013, p_clust_ = 1, d = .003; alpha: t_27_ = .256, p_clust_ = .186, d = .048; beta: t_27_ = -.669, p_clust_ = 1, d = .127; gamma: t_27_ = .324, p_clust_ = .289, d = .061; Fig. 2C).

**Figure 2.**
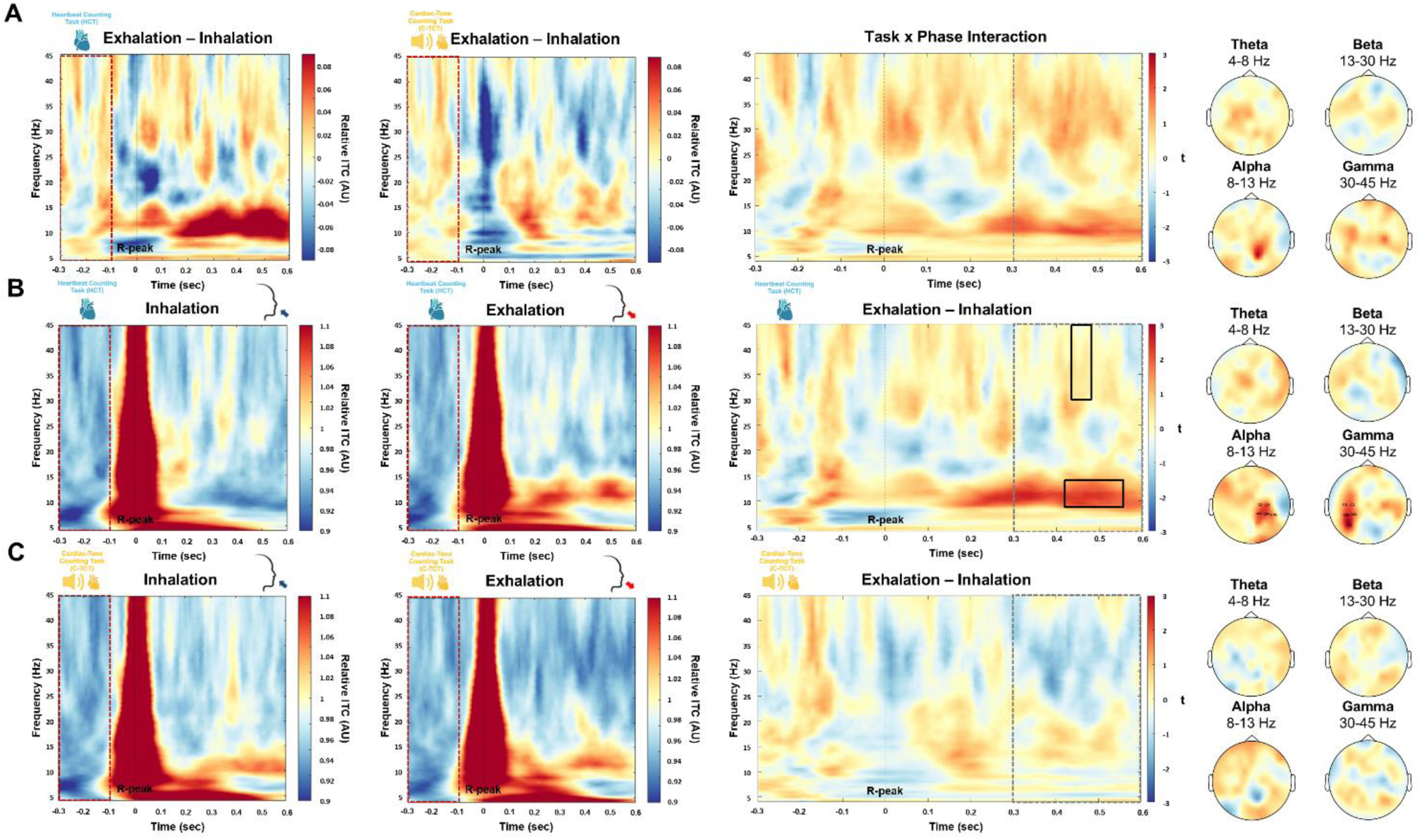
Heartbeat-related inter-trial coherence changes across respiratory phases and tasks. (A) Grand-averaged heartbeat-related inter-trial coherence changes between respiratory phases (exhalation vs. inhalation) during the heartbeat counting task (left), the cardiac-tone counting task (middle), and their task × phase interaction (right). (B) Grand-averaged heartbeat-related inter-trial coherence in the heartbeat counting task during inhalation (left), exhalation (middle), and the exhalation-minus-inhalation difference (right). The black rectangle marks significant differences between exhalation and inhalation (cluster-corrected). Topographical distributions illustrate significant differences, with labels indicating significant electrodes. (C) Grand-averaged heartbeat-related inter-trial coherence in the cardiac-tone counting task during inhalation (left), exhalation (middle), and the exhalation-minus-inhalation difference (right). The red dotted area represents the baseline window (-300 to -100 msec relative to R-peak onset). The grey dotted area marks the temporal window of interest for statistical analysis (300-600 msec after the R-peak). Abbreviations: AU = arbitrary unit.

### 2.4 Heartbeat-related functional connectivity is modulated by task and the respiratory phase

Heartbeat-related FC between all pairs of electrodes was estimated for each participant, task, and respiratory phase separately in the alpha and theta bands. We focused on these frequency bands because they showed significant modulation of heartbeat-related power and ITC between respiratory phases. We conducted permutation-based t-tests across the entire scalp, using the relative (i.e., active divided by baseline cardiac time-window, see section 4.5) imaginary part of coherency (iCOH) as the dependent variable, with task (HCT vs. C-TCT), respiratory phase (inhalation vs. exhalation) and their interaction as within-participant factors. The analysis revealed a significant task × phase interaction in the theta band over the parieto-occipital cortices, alongside a negative effect in frontal regions (Fig. 3A, Supplementary Table 1). Planned t-tests (see section 4.6) indicated greater theta-band FC during HCT in exhalation compared to inhalation, predominantly lateralized to the right hemisphere, spanning from prefrontal to parieto-occipital areas (Fig. 3B, Supplementary Table 2). During C-TCT, while some theta-band FC increases were observed during exhalation relative to inhalation, these were accompanied by a marked reduction in connectivity in the right hemisphere (Fig. 3C, Supplementary Table 3). In the alpha band, we observed a significant task × phase interaction, which was most pronounced over the right frontal cortices (Fig. 3D, Supplementary Table 4). This effect was driven by a significant increase in FC between right frontal and central regions during exhalation compared to inhalation during HCT (Fig. 3E, Supplementary Table 5). On the other hand, during C-TCT, FC was significantly reduced during exhalation relative to inhalation, particularly over the right frontal and prefrontal regions (Fig. 3F, Supplementary Table 6).

**Figure 3.**
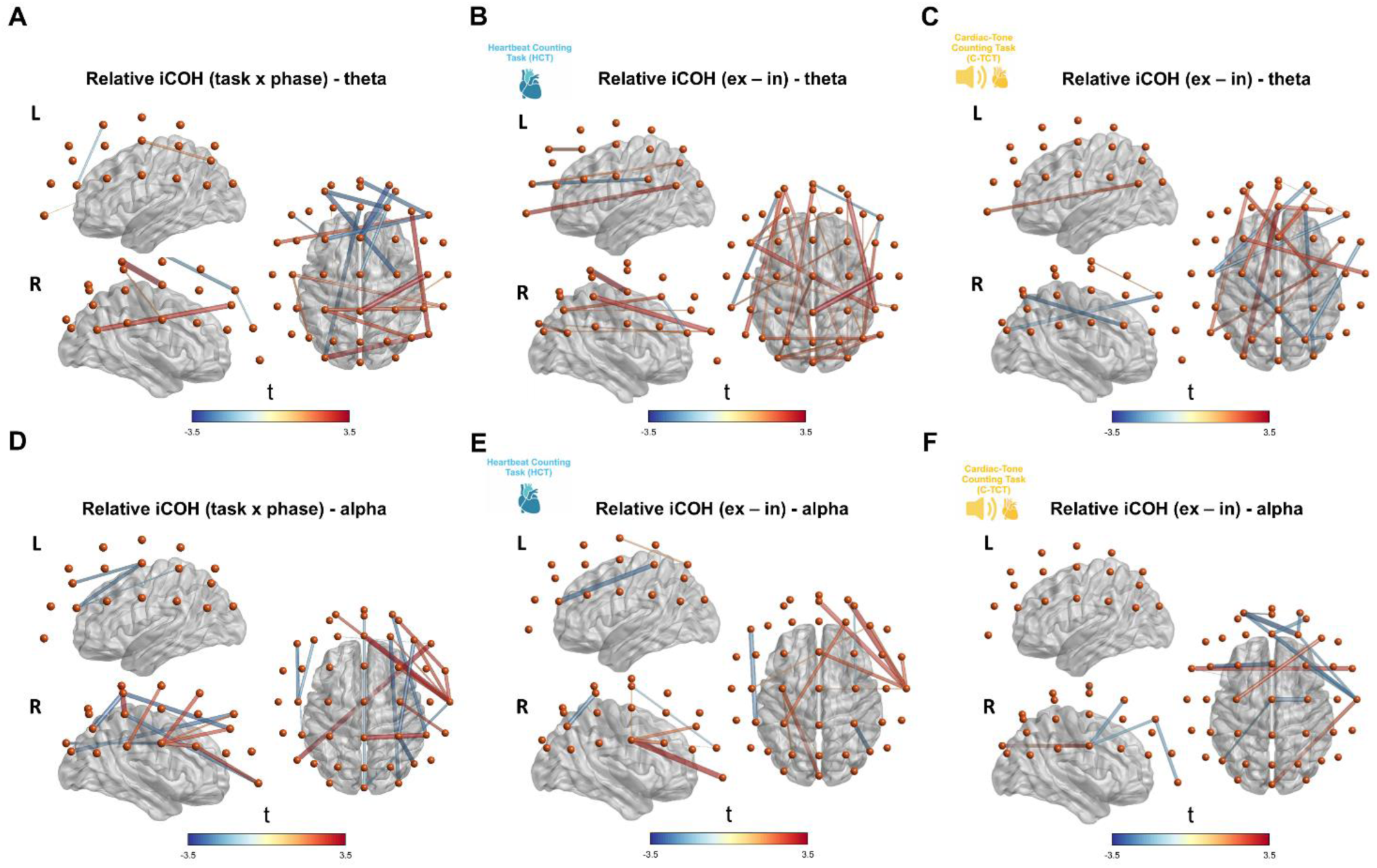
Heartbeat-related functional connectivity changes across respiratory phases and tasks. Visualization of functional connectivity (relative imaginary part of coherency) for: (A) Significant task × phase interaction in the theta band. (B) Significant differences between exhalation and inhalation in the theta band during the heartbeat counting task. (C) Significant differences between exhalation and inhalation in the theta band during the cardiac-tone counting task. (D) Significant task × phase interaction in the alpha band. (E) Significant differences between exhalation and inhalation in the alpha band during the heartbeat counting task. (F) Significant differences between exhalation and inhalation in the alpha band during the cardiac-tone counting task. Red lines indicate significant (p < .01) positive changes in the relative imaginary part of coherency, while blue lines indicate significant negative changes. Line thickness represents the strength of the difference (t-score). Abbreviations: iCOH = imaginary part of coherency, Ex = exhalation, In = inhalation, L = left, R = right.

To control for potential artificial increases in heartbeat-related FC induced by CFA, we examined whether relative alpha- and theta-band iCOH increased during exhalation compared to inhalation between the ECG and EEG electrodes that exhibited significant iCOH changes. Permutation-based dependent-samples t-tests comparing relative iCOH between exhalation and inhalation during HCT revealed a significant increase between the ECG and F3 electrode in the theta band (t_27_ = 1.07, p = .004) and between the ECG and Fpz electrode in the alpha band (t_27_ = 1.85, p = .006). No significant changes in ECG connectivity were found during C-TCT. These results show that the widespread functional connectivity changes observed between respiratory phases were not attributable to CFA-related connectivity effects.

### 2.5 Heartbeat-related graph metrics are modulated by task and the respiratory phase

A total of eight LMEMs were estimated including each participant, task, respiratory phase, cardiac time-window, and threshold. These models were computed for four graph metrics separately in the alpha and theta frequency bands: density, clustering coefficient, global efficiency, and local efficiency. The full models including the three-way interaction across all graph theory metrics were:

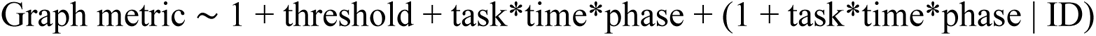

In the theta frequency band, beyond the expected main effect of the threshold factor, we observed a significant main effect of time (i.e., baseline vs. active cardiac time-windows) across all four graph theory metrics (density: b = .037, SE = .015, t = 2.402, p = .023; clustering coefficient: b = .039, SE = .014, t = 2.781, p = .009; local efficiency: b = .065, SE = .023, t = 2.834, p = .008; global efficiency: b = .048, SE = .019, t = 2.504, p = .019; Fig. 4A-D, Supplementary Tables 7-10). This suggests that the heartbeat generally reduces connectivity in the theta band. Despite the three-way interaction between task, time, and phase was not significant (b = .041, SE = .062, t = .664, p = .512), planned t-tests (see section 4.6) revealed a significant heartbeat-related reduction in network density during inhalation in HCT (t = -3.566, p = .001, p_FDR_ = .006), resulting in higher density during exhalation compared to inhalation in the active cardiac time-window (t = 2.672, p = .013, p_FDR_ = .025; Fig. 4A, Supplementary Table 11). A similar pattern was observed for the clustering coefficient in HCT, where t-tests indicated a heartbeat-related reduction during inhalation (t = -4.01, p < .001, p_FDR_ = .002), leading to higher theta clustering coefficients during exhalation than inhalation in the active cardiac time-window (t = 2.383, p = .024, p_FDR_ = .049; Fig. 4B, Supplementary Table 12). Furthermore, both heartbeat-related local and global efficiency significantly decreased during inhalation in HCT (local efficiency: t = -3.988, p < .001, p_FDR_ = .002, Fig. 4C; global efficiency: t = -3.793, p < .001, p_FDR_ = .003, Fig. 4D, Supplementary Table 13-14). However, no significant differences were observed between inhalation and exhalation in the active cardiac time-window (local efficiency: t = 1.967, p = .06; global efficiency: t = 1.444, p = .16).

**Figure 4.**
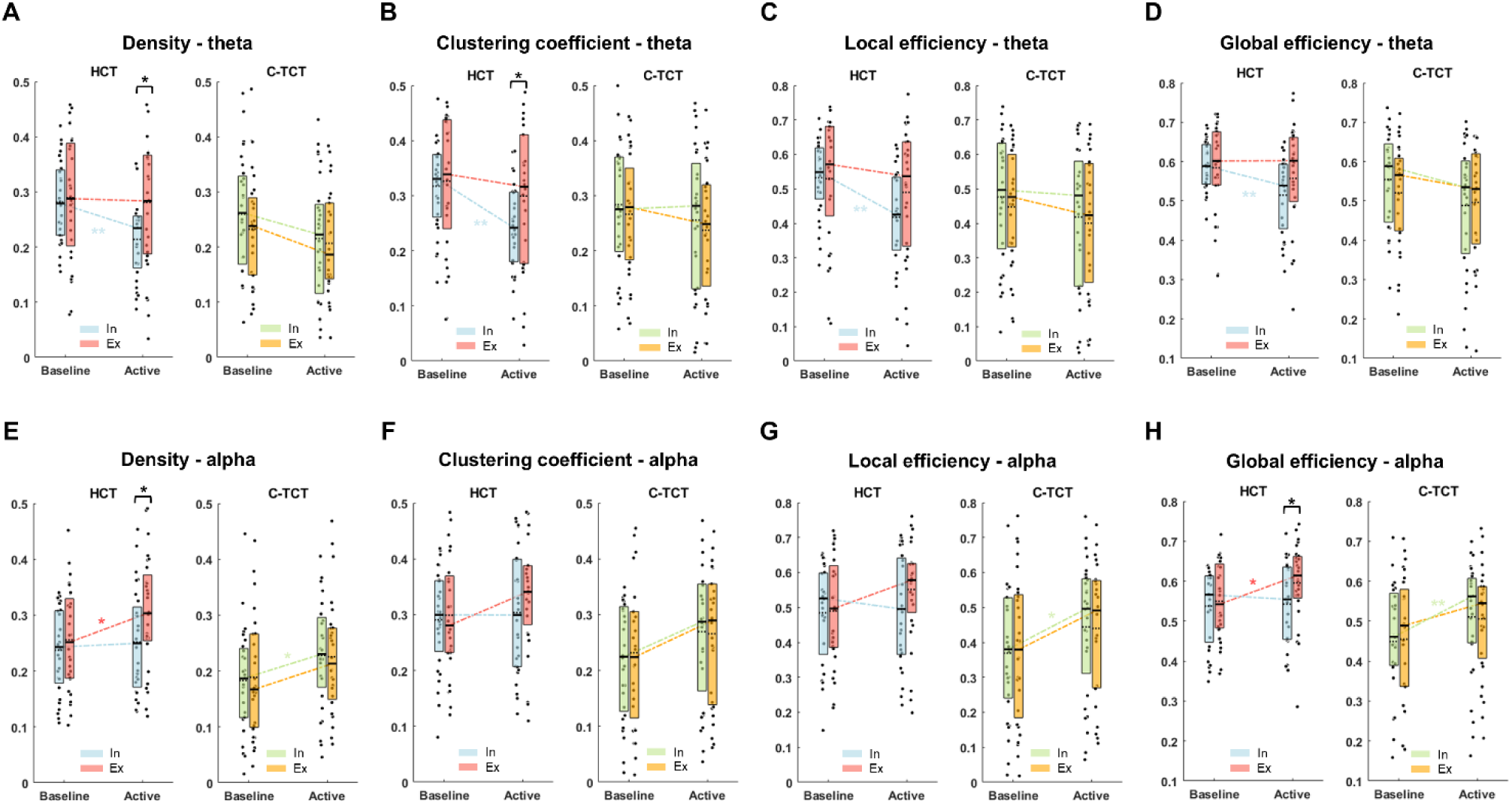
Graph theory metrics across respiratory phases and tasks. Box plots displaying descriptive statistics for graph-theoretic metrics before (baseline) and after (active) the heartbeat during inhalation and exhalation in the heartbeat counting task and the cardiac-tone counting task. (A) Network density in the theta band. (B) Network clustering coefficient in the theta band. (C) Network local efficiency in the theta band. (D) Network global efficiency in the theta band. (E) Network density in the alpha band. (F) Network clustering coefficient in the alpha band. (G) Network local efficiency in the alpha band. (H) Network global efficiency in the alpha band. Significant planned t-tests are indicated by asterisks (**p < .01, *p < .05). The y-axis units reflect normalized ratios (arbitrary units). Abbreviations: In = inhalation, Ex = exhalation, HCT = heartbeat counting task, C-TCT = cardiac-tone counting task.

In the alpha band, the heartbeat consistently induced a significant increase in alpha connectivity metrics (main effect of time for density: b = -.036, SE = .007, t = -4.72, p < .001; clustering coefficient: b = -.034, SE = .009, t = -3.73, p < .001; local efficiency: b = -.052, SE = .013, t = -4.11, p < .001; global efficiency: b = -.041, SE = .009, t = -4.34, p < .001, Supplementary Tables 15-18). Additionally, a significant main effect of task was observed, indicating higher alpha connectivity during HCT compared to C-TCT (density: b = .056, SE = .026, t = 2.13, p =.043; clustering coefficient: b = .061, SE = .029, t = 2.06, p = .049; local efficiency: b = .096, SE = .045, t = 2.18, p = .038; global efficiency: b = .076, SE = .032, t = 2.35, p = .026; as can be inferred from Fig. 4E–H, Supplementary Tables 15-18). Despite the three-way interaction between task, time, and phase was not significant (b = .04, SE = .039, t = 1.55, p = .3), planned t-tests (see section 4.6) for graph density revealed a significant heartbeat-evoked increase in density during exhalation in HCT (t = 2.82, p = .009, p_FDR_ = .036), leading to higher alpha density at exhalation compared to inhalation within the active cardiac time-window (t = 2.497, p = .019, p_FDR_ = .038; Fig. 4E, Supplementary Table 19). In contrast, during C-TCT, the heartbeat increased alpha density only during inhalation (t = 3.188, p = .004, p_FDR_ = .014), and resulted in no significant differences between respiratory phases (t = -.132, p = .896). For the clustering coefficient, no significant heartbeat-evoked differences between respiratory phases were significant (HCT: t = 1.615, p = .118; C-TCT: t = -.214, p = .833; Fig. 4F, Supplementary Table 20). For local efficiency, a significant heartbeat-induced increase in the alpha band was observed only during inhalation in C-TCT (t = 2.871, p = .008, p_FDR_ = .031, Fig. 4G, Supplementary Table 21). Finally, for global efficiency, the heartbeat evoked increased connectivity during exhalation (t = 2.62, p = .014, p_FDR_ = .038), leading to higher alpha global efficiency at exhalation compared to inhalation in the active cardiac time-window during HCT (t = 2.499, p = .018, p_FDR_ = .038; Fig. 4H, Supplementary Table 22). Conversely, in C-TCT, global efficiency was enhanced by the heartbeat only during inhalation (t = 3.751, p < .001, p_FDR_ = .003), but no significant differences were found between respiratory phases in the active phase of the cardiac cycle (t = -.292, p = .772).

### 2.6 Participant-level relationships between heartbeat-modulated cortical oscillations and heartbeat-evoked potential

We assessed the spatial overlap between the distributions of significant heartbeat-related alpha power and HEP amplitude changes across respiratory phases (Δalpha power and ΔHEP). As previously reported (Zaccaro et al., 2024), significant ΔHEP changes were localized to right centro-frontal electrodes (AFz, F1, Fz, F2, F4, FC1, FCz, FC2, FC4, C1, Cz, C2, and C4) from 360 to 528 msec after the R-peak (t_27_ = 3.29, p_clust_ = .016, d = .621; Fig. 5A). Although ΔHEP differences occurred in more anterior regions of the scalp compared to the more central Δalpha power differences (same cluster as in section 2.2; Fig. 5B), both distributions converged spatially in fronto-central areas, slightly lateralized to the right hemisphere. To statistically quantify this relationship, we computed Spearman’s correlation between participant-level mean heartbeat-related Δalpha power with the corresponding mean ΔHEP values within the late significant time window of heartbeat-related power effects (390–600 msec after the R-peak). A significant positive correlation was found over frontal electrodes (Fig. 5C), including AFz (rho = .519, p_FDR_ = .044), AF3 (rho = .521, p_FDR_ = .044), AF4 (rho = .522, p_FDR_ = .044), F4 (rho = .577, p_FDR_ = .038), and Fz (rho = .573, p_FDR_ = .038). A negative correlation was also observed over channel P6 (rho = -.512, p_FDR_ = .044). When examining correlations between ΔHEP amplitude and heartbeat-related Δpower in other frequency bands during both HCT and C-TCT, no significant results were found, confirming that the effects were specific to the alpha band during HCT. Additionally, no significant correlations were observed between ΔHEP amplitude and ΔITC, nor between Δpower and ΔITC across any frequency band in either task (during HCT or C-TCT). Further details on the Spearman’s rho values across tasks, frequencies, and electrodes are provided in the supplementary materials (Supplementary Table 23). Finally, no correlations were found between HEP, power, or ITC in any frequency band when analysing their absolute values recorded separately for inhalation and exhalation in either task (Supplementary Table 24).

**Figure 5.**
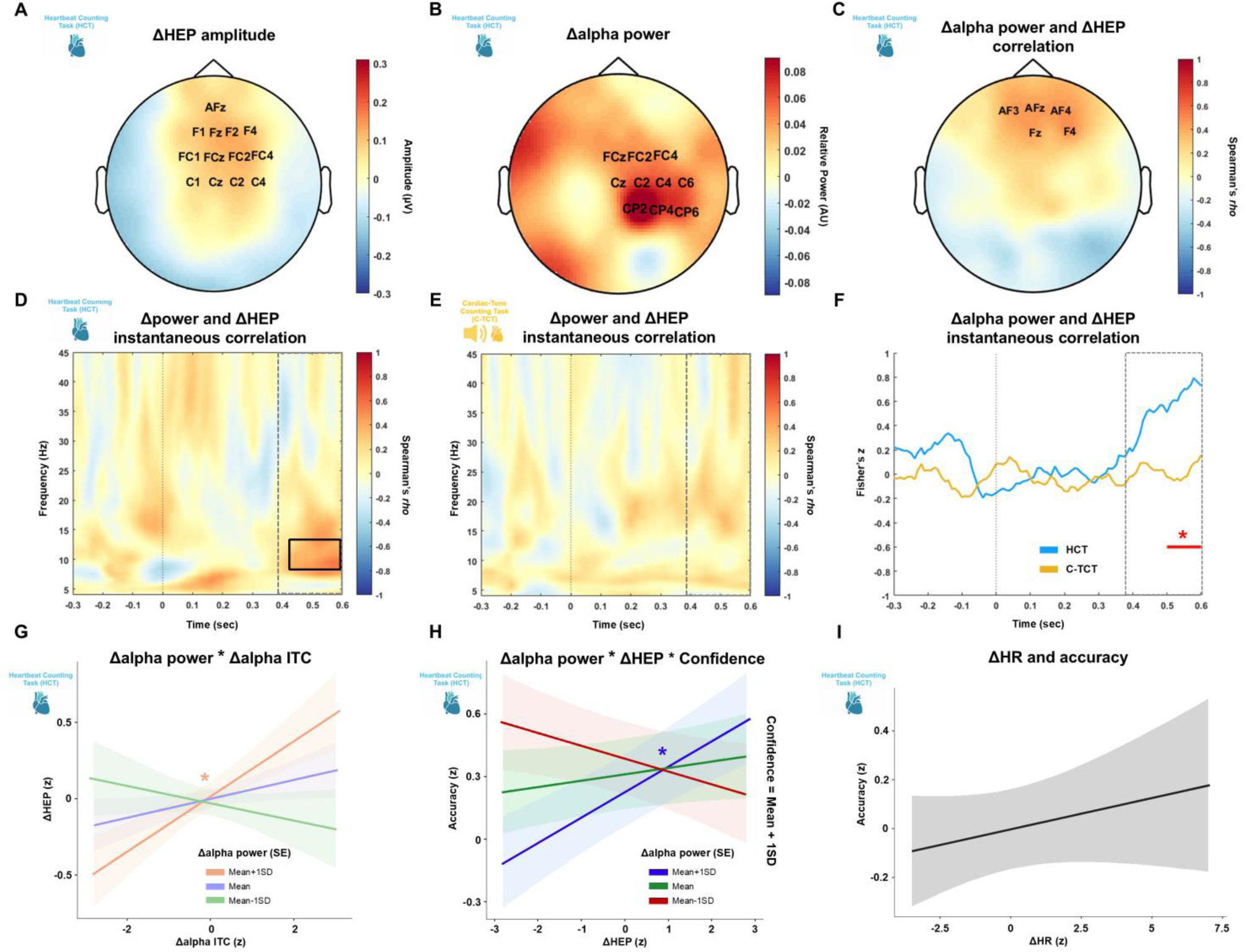
Relationships between heartbeat-modulated cortical oscillations, heartbeat-evoked potentials, and task accuracy. (A) Topographical distribution of heartbeat-evoked potential differences between exhalation and inhalation during the heartbeat counting task. Labels indicate significant electrodes. (B) Topographical distribution of heartbeat-related alpha power differences between exhalation and inhalation during the heartbeat counting task. Labels indicate significant electrodes. (C) Topographical distribution of Spearman’s correlation between heartbeat-related alpha power and heartbeat-evoked potential differences between exhalation and inhalation during the heartbeat counting task. Labels indicate significant electrodes. (D) Time-frequency representation of the instantaneous Spearman’s correlation between heartbeat-related power and heartbeat-evoked potential differences between exhalation and inhalation during the heartbeat counting task. Data are averaged across significant electrodes (AFz, AF3, AF4, F4, Fz). The black rectangle highlights significant correlations (FDR-corrected). The grey dotted area marks the temporal window of interest for statistical analysis (390-600 msec after the R-peak). (E) Time-frequency representation of the instantaneous Spearman’s correlation between heartbeat-related power and heartbeat-evoked potential differences between exhalation and inhalation during the cardiac-tone counting task. The grey dotted area marks the temporal window of interest for statistical analysis (390-600 msec after the R-peak). (F) Fisher’s z-transformed instantaneous Spearman’s correlation values comparing the heartbeat counting task and the cardiac-tone counting task. Data are averaged in the alpha band (8-13 Hz) across significant electrodes (AFz, AF3, AF4, F4, Fz). The grey dotted area marks the temporal window of interest for statistical analysis (390-600 msec after the R-peak). Significant differences are indicated by asterisks (*p < .05, FDR-corrected). (G) Linear mixed-effects model showing the significant interaction between Δalpha power and Δalpha inter-trial coherence during the heartbeat counting task. Simple effects analysis reveals a significant relationship between Δalpha inter-trial coherence and Δheartbeat-evoked potential only when Δalpha power is high (mean + 1SD). Significant relationships are indicated by asterisks (*p < .05). (H) Linear mixed-effects model showing the significant interaction between Δheartbeat-related alpha power, Δ heartbeat-evoked potential, and confidence during the heartbeat counting task. Simple effects analysis reveals a significant relationship between Δ heartbeat-evoked potential and accuracy only when both Δheartbeat-related alpha power and confidence are high (mean + 1SD). Significant relationships are indicated by asterisks (*p < .05). (I) Linear mixed-effects model during the heartbeat counting task showing no relationship between ΔHR and accuracy. Abbreviations: HEP = heartbeat-evoked potential, ITC = inter-trial coherence, HCT = heartbeat counting task, C-TCT = cardiac-tone counting task, ECG = electrocardiogram, SD = standard deviation, SE = standard error, AU = arbitrary unit.

We computed the instantaneous group-level Spearman’s correlation between heartbeat-related Δpower and the corresponding ΔHEP values for each time point and frequency band, averaged across the previously identified channels (AFz, AF3, AF4, F4, and Fz), during both the HCT and C-TCT tasks. A significant correlation emerged in the alpha band exclusively during HCT, within a late time window from 422 to 600 msec post-R-peak (Fig. 5E, Supplementary Tables 25-26). We compared instantaneous correlations between HCT and C-TCT by applying Fisher’s z-transformation to Spearman’s rho values averaged across the previously selected channels in the alpha band. This analysis revealed significantly stronger correlations for HCT relative to C-TCT between 512 and 600 msec post-R-peak (Fig. 5F, Supplementary Table 27). These findings indicate that the coupling between ΔHEP amplitude and heartbeat-related Δalpha power is selectively enhanced during cardiac interoceptive attention.

### 2.7 Trial-level relationships between heartbeat-modulated cortical oscillations and heartbeat-evoked potential

To examine the relationships between HCT trial-level changes in HEP, heartbeat-related power, and ITC across respiratory phases, as well as their associations with accuracy and cardiac physiology, we applied multiple LMEMs. Following previous studies (Fló et al., 2024; Park et al., 2018), we first tested the relationship between changes in heartbeat-related alpha power and ITC across respiratory phases and confirmed that no significant relationship was found between the two measures (b = .022, SE = .056, t = .399, p = .69). Using ΔHEP as the dependent variable, the best-fitting model was:

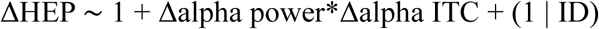

The model revealed a significant interaction between Δalpha power and Δalpha ITC (b = .118, SE = .056, t = 2.096, p = .037; Fig. 5G), suggesting that Δalpha power moderates the effects of Δalpha ITC on ΔHEP. A simple effects analysis of the interaction indicated a significant relationship between Δalpha ITC and ΔHEP only when Δalpha power was high (mean + 1SD: b = .18, SE = .078, t = 2.306, p = .022; mean: b = .062, SE = .057, t = 1.089, p = .277; mean – 1SD: b = -.056, SE = .082, t = .686, p = .493). No main effects were observed for Δalpha power (b = .027, SE = .057, t = .479, p = .632), Δalpha ITC (b = .062, SE = .057, t = 1.089, p = .277), or the intercept (b = .003, SE = .061, t = .051, p = .959). Notably, no main effects or interactions involving ΔHR (b = .015, SE = .059, t = .258, p = .797) or ΔECG amplitude (b = .011, SE = .059, t = .188, p = .851) were found, therefore these variables were not included in the final model (Supplementary Tables 28-30).

For HCT trial-level accuracy as the dependent variable, the best-fitting model was:

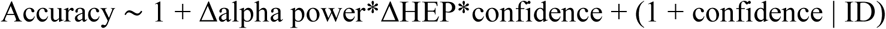

The model revealed a significant three-way interaction between ΔHEP, Δalpha power, and confidence (b = .068, SE = .03, t = 2.264, p = .025; Fig. 5H), indicating that changes in alpha power and confidence together moderated the effect of ΔHEP on task accuracy. Simple effects analysis showed a significant relationship between ΔHEP and accuracy only when both Δalpha power and confidence were high (mean + 1SD: b = .127, SE = .048, t = 2.628, p = .009). Additionally, significant main effects were observed for confidence (b = .332, SE = .082, t = 4.031, p < .001) and Δalpha power (b = -.062, SE = .03, t = -2.059, p = .04) on accuracy. No main effects were found for the intercept (b = -.011, SE = .152, t = -.072, p = .943), or ΔHEP (b = .033, SE = .028, t = 1.184, p = .238). There were no significant interactions between Δalpha power and ΔHEP (b = .027, SE = .025, t = 1.112, p = .267), Δalpha power and confidence (b = -.022, SE = .033, t = -.042, p = .493), or ΔHEP and confidence (b = -.001, SE = .03, t = -.042, p = .967). Notably, no main effects or interactions involving Δalpha ITC (b = .028, SE = .032, t = .858, p = .392), ΔHR (b = .029, SE = .043, t = .686, p = .493), or ΔECG amplitude (b = -.022, SE = .041, t = -.549, p = .584) were found, therefore these variables were not included in the final model (Supplementary Tables 31-33). Fig. 5I shows the non-significant relationship between ΔHR and HCT accuracy.

Performing an LMEM for C-TCT with ΔHEP as the dependent variable was not feasible due to insufficient random coefficient variance. A general linear model was implemented, which was:

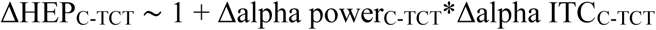

This model revealed no significant interaction between Δalpha power and Δalpha ITC (b = - .112, SE = .062, t = -1.815, p = .071), nor any main effects for the intercept (b = .003, SE = .057, t = .055, p = .956), Δalpha power (b = .003, SE = .058, t = .046, p = .964), or Δalpha ITC (b = .004, SE = .058, t = .066, p = .947).

Using the trial-level C-TCT accuracy as the dependent variable, the best-fitting model was:

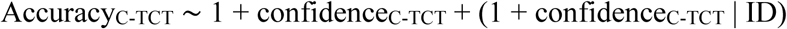

This model indicated that, during the C-TCT, only the main effects of confidence (b = .378, SE = .11, t = 3.433, p = .002) predicted task accuracy (Supplementary Table 34-35).

## 3. Discussion

The multilevel connections between the cardiac and respiratory systems suggest that breathing continuously shapes cardiac interoception at both behavioural and neural levels, an aspect that has been often overlooked in interoception research. In previous studies, we found greater late HEP positivity during exhalation compared to inhalation in healthy participants performing two cardiac interoceptive tasks (HBD and HCT, Zaccaro et al., 2022, 2024). In the present study, we aimed to extend previous findings to the time-frequency domain by re-analysing the same dataset and focusing on the interoceptive HCT and the exteroceptive C-TCT (Zaccaro et al., 2024). As hypothesized, and consistent with previous time-domain results, we observed increased late heartbeat-related power and ITC in exhalation compared to inhalation during the HCT. These effects were observed in the alpha and theta frequency bands over right fronto-centro-parietal electrodes. In contrast, no differences were found between exhalation and inhalation during C-TCT. During HCT, we also observed a widespread increase in heartbeat-related FC in the alpha and theta bands during exhalation compared to inhalation, mostly lateralized to the right fronto-central regions. Conversely, during C-TCT, alpha and theta FC in the right hemisphere decreased during exhalation compared to inhalation. Graph theory analysis further clarified these findings. During HCT, the heartbeat induced a reduction in theta-band graph connectivity during inhalation and increased alpha-band graph connectivity during exhalation. Consequently, both theta-band (density and clustering coefficient) and alpha-band (density and global efficiency) heartbeat-modulated graph connectivity metrics were higher during exhalation than inhalation. No changes were observed in the theta-band graph connectivity during C-TCT, whereas in the alpha band the heartbeat caused some inhalation-related increases that did not result in significant differences between respiratory phases. At the participant level, heartbeat-related alpha power and HEP changes across respiratory phases (Δalpha power and ΔHEP) were positively correlated over frontal electrodes. This relationship was specific to HCT and was not observed for theta or other frequency bands nor ITC changes across respiratory phases (ΔITC). At the HCT-trial level, Δalpha ITC predicted ΔHEP only when Δalpha power was high, whereas ΔHEP predicted interoceptive accuracy only when both Δalpha power and confidence were high. Crucially, these relationships were independent of cardiac physiology and were absent during C-TCT. Overall, these results indicate that when participants focus on their cardiac sensations, the heartbeat elicits higher power, ITC, and FC during exhalation compared to inhalation primarily in the alpha band, which in turn predicts cardiac interoceptive accuracy.

To interpret these findings, we outline a model of cardio-respiratory interactions within the framework of interoceptive predictive coding. The predictive coding framework posits that the brain continuously interprets sensory information in light of expectations based on past experience (Clark, 2013; Friston, 2010). When sensory data deviates from predicted outcomes, the brain generates prediction errors. Prediction errors are resolved through either “perceptual inference” (where prediction errors update prior models) or “active inference” (where actions modify sensory inputs to match predictions) (Friston et al., 2017; Hohwy, 2012; Palmer et al., 2019; Parr et al., 2018). The selection of either strategy depends on the relative reliability of sensory data versus prior beliefs. This reliability, known as “precision”, determines the weighting assigned to sensory inputs or priors, and its modulation via top-down attention is referred to as “precision-weighting” (Brown & Friston, 2012; Feldman & Friston, 2010; Kanai et al., 2015). The interplay between top-down predictions and bottom-up prediction errors also occurs in the interoceptive domain, where information about bodily states is processed within a hierarchical interoceptive network (Barrett & Simmons, 2015; Ondobaka et al., 2017; Owens et al., 2018; Petzschner et al., 2021; Pezzulo, 2014; Seth, 2013; Seth & Friston, 2016; Smith et al., 2017). According to this framework, cardiac signals during systole are usually predicted and attenuated by the brain and do not generate conscious perception (Ainley et al., 2016; Allen, Levy, et al., 2022). Furthermore, cardiac signals are generally inhibitory (Lacey, 1978; Paci et al., 2024; Rau & Elbert, 2001; Saltafossi et al., 2023), so when perceptual sensitivity is required, the brain actively reduces the impact of systole through a parasympathetically driven decrease in HR (Skora et al., 2022). On the contrary, when cardiac interoceptive accuracy is required, the brain increases the precision of cardiac signals through attentional focus, leading to higher cardiac prediction errors, prior updating, and conscious heartbeat perception (Ainley et al., 2016; Petzschner et al., 2019). In this scenario, cardiac interoceptive accuracy is determined by an increase in the precision (saliency) of each heartbeat, rather than by an increase in HR (Leganes-Fonteneau et al., 2021; MacKinnon et al., 2013; Larsson et al., 2021).

The present findings, together with previous studies (Zaccaro et al., 2022, 2024), highlight the role of cardio-respiratory interactions in interoceptive perception, which occur across at least three levels of the interoceptive hierarchy: the peripheral, brainstem, and cortical levels. At the peripheral level, cardiovascular and respiratory processes are coupled on a cycle-by-cycle basis by the baroreflex (Draghici & Taylor, 2016; Magder, 2018; Shaffer et al., 2014; Shaffer & Venner, 2013). During inhalation, the decrease in thoracic pressure reduces left ventricular filling and stroke volume, leading to a decrease in blood pressure and baroreceptor activity. This initiates the baroreflex, mediated by the nucleus of the solitary tract (NTS) and the nucleus ambiguous, which stabilises blood pressure and cardiac output by increasing the HR when heartbeats are weak. In contrast, higher baroreceptor activity during exhalation leads to the reduction of the sympathetic tone and HR. This interplay creates oscillations in HR between respiratory phases (RSA; Ben-Tal et al., 2012; Bernardi et al., 2001; Giardino et al., 2003; Hayano & Yasuma, 2003) and leads to slower but stronger (more salient) cardiac signals during exhalation than inhalation (Larsson et al., 2021). At the brainstem level, cardiac and respiratory afferents are then integrated into the caudal NTS, which receives both aortic and carotid baroreceptors and rapidly adapting pulmonary receptors (RARs) terminations from the vagus and glossopharyngeal nerves (Neuhuber & Berthoud, 2022; Smith et al., 2017; Zoccal et al., 2014). As RARs are mainly activated by lung stretch during inhalation and are relatively silent during exhalation (Del Negro et al., 2018; Krohn et al., 2023; Kubin et al., 2006; Noble & Hochman, 2019), weak cardiac signals are integrated with strong respiratory stimuli during inhalation and, on the other hand, strong cardiac signals are processed with minimal respiratory interference during exhalation. In this context, recent predictive coding models of respiratory interoception (Allen, Varga, et al., 2022; Boyadzhieva & Kayhan, 2021; Braendholt et al., 2023; Corcoran et al., 2018) propose that similar to the heartbeat, the brain generates predictions about the interoceptive consequences of each inhalation, attenuating or “predicting them away” to adaptively optimize perception and action. We propose that in the NTS, as weak cardiac inputs are processed together with strong respiratory inputs, the brain is more likely to attenuate cardiac signals together with respiratory signals during inhalation (see Al et al., 2020, 2021, for a similar interpretation in the somatosensory domain), leading to reduced cardiac interoception during inhalation compared to exhalation (Zaccaro et al., 2022, 2024). Finally, at the highest cortical level, a similar predictive process may also occur, given the topological overlap between cardiac and respiratory signal processing in the cingulate, orbitofrontal/medial prefrontal and somatosensory cortices, amygdala, insula, and precuneus (Engelen et al., 2023; Goheen et al., 2022; Heck et al., 2022; Kluger & Gross, 2021; Park & Blanke, 2019).

We further propose that, at this highest level, reduced cortical excitability during exhalation, particularly in the insula and somatosensory regions, may enhance cardiac interoception, as reflected in increased heartbeat-modulated cortical oscillations in the alpha range. Studies using both periodic and aperiodic MEG/EEG signal analysis have found higher cortical inhibition in the late phase of exhalation and in the transition from exhalation to inhalation (Kluger et al., 2023; Nakamura et al., 2024). Furthermore, neuroimaging studies have found that interoceptive accuracy is mediated by insula inhibition, suggesting an “addition by subtraction” mechanism for interoceptive attention (Farb et al., 2023; Kuehn et al., 2016; Wiebking et al., 2015). Complementary EEG evidence has shown enhanced alpha power during interoceptive tasks (Crivelli & Balconi, 2025; Kritzman et al., 2022; Mccraty et al., 2009), which correlates with task accuracy (Rominger et al., 2025; Villena-González et al., 2017). This has been interpreted as an active inhibitory mechanism that suppresses the processing of distracting signals (Cooper et al., 2003; Foxe & Snyder, 2011; Jensen & Mazaheri, 2010; Klimesch et al., 2007; Van Diepen et al., 2019). Additionally, in the context of predictive coding, alpha activity has been linked to the precision-weighting of prediction errors and prior updating (Bastos et al., 2012; Braendholt et al., 2023; Friston, 2019; Friston et al., 2015; Sherman et al., 2016). Therefore, synthesizing this body of literature, our findings of increased heartbeat-related alpha power, ITC, and FC during exhalation can be interpreted as neurophysiological signatures of enhanced cardiac interoception. This enhancement likely reflects a state of increased top-down attentional allocation toward interoceptive signals (Molle & Coste, 2022), supported by active inhibition of irrelevant sensory input facilitated by heightened cortical inhibition (particularly in insular and somatosensory regions) during the exhalation phase. Such a configuration may optimize the precision-weighting of cardiac prediction errors, ultimately contributing to improved heartbeat perception and confidence.

As we observed increases in both heartbeat-related alpha power and ITC during exhalation, we cannot conclusively determine whether the observed HEP modulations between respiratory phases result from an enhanced phase-reset mechanism. Such modulations would be supported by ITC increases without concurrent power increases (Lopour et al., 2013; Park et al., 2018). We propose that respiration modulates HEP amplitude through increases in both evoked amplitude and phase-reset mechanisms in the alpha band. This interpretation is supported by the moderating effect of heartbeat-related power changes on the ITC-HEP relationship: while power and ITC changes across respiratory phases did not correlate (as in Park et al., 2018), ITC increases were positively associated with HEP enhancements only when power increases were also high. This suggests that the two processes (alpha power and ITC increases) stem from distinct yet interactive mechanisms, both essential for cardiac perception. We speculate that positive Δalpha power may serve as an index of interoceptive sensory gating mediated by top-down attention (Crivelli & Balconi, 2025; Kritzman et al., 2022; Molle & Coste, 2022; Rominger et al., 2025; Villena-González et al., 2017), whereas increased Δalpha ITC may reflect increased precision-weighting of cardiac prediction errors (Park et al., 2018; Petzschner et al., 2019).

Several additional questions remain to be answered. First, another key mechanism in cardio-respiratory interoceptive interactions may involve somatosensory and motor pathways (Cameron & Minoshima, 2002; Kern et al., 2013; Khalsa et al., 2009). Since inhalation is an active process driven by motor commands, the brain generates predictions about the sensory consequences of chest expansion to attenuate somatosensory feedback. Because somatosensory and tactile inputs from inhalation overlap with those generated by heartbeats at the level of the somatosensory cortices, the brain may primarily interpret cardiac tactile signals as a byproduct of chest expansion (Al et al., 2020, 2021). In contrast, exhalation is largely passive, producing weaker and less precise expected sensory consequences, which may lead to reduced predictive suppression of cardiac-related tactile inputs. Second, the breathing pathway (nasal vs. mouth) may also play a role in cardio-respiratory interoceptive interactions (Fontanini & Bower, 2006; Tort et al., 2018). Further investigation is needed to determine whether inhalation disrupts cardiac interoception specifically during nasal breathing or whether this effect persists during mouth breathing. Third, directing attention to the heart reduces respiratory rate, likely as a form of respiratory active inference to increase cardiac salience (Candia-Rivera et al., 2022; Zaccaro et al., 2022, 2024). Future studies should investigate the role of voluntary respiratory modulation (in depth, phase, or frequency) in shaping cardiac interoceptive accuracy, emphasizing the contribution of respiratory active inference to cardiac interoceptive attention.

In conclusion, our findings indicate that cardio-respiratory interactions occur at high levels of the interoceptive hierarchy (Smith et al., 2017; Suksasilp & Garfinkel, 2022). We propose that these interactions are fundamental to understanding cardiac interoception, as they provide additional explanatory power for how the central nervous system optimizes the interplay between interoceptive processes. Furthermore, we suggest that such interactions extend beyond the cardiac and respiratory systems and that the predictive coding framework offers valuable insights for their investigation. Finally, we hypothesize that the dynamic interplay between the cardio-respiratory system and the brain has functional significance and may be altered in clinical conditions. This is supported by evidence showing that a respiratory perturbation (breath-holding) enhanced the interoceptive precision assigned to cardiac signals only in healthy individuals, whereas no such effect was observed in individuals with various psychiatric disorders, including depression, anxiety, substance use, and eating disorders (Lavalley et al., 2024; Smith et al., 2020, 2021). Thus, difficulties in adjusting the precision of afferent cardiac signals across respiratory phases during spontaneous breathing may represent a transdiagnostic marker of vulnerability to somatic and mental health disorders.

## 4. Materials and methods

### 4.1 Ethics statement

The study from which the original dataset was derived (Zaccaro et al., 2024) was approved by the Institutional Review Board of the Department of Psychological Sciences, Health and Territory at the “G. d’Annunzio” University of Chieti-Pescara, Italy (Protocol Number 44_26_07_2021_21016). The study adhered to the ethical guidelines of the Italian Association of Psychology and the Declaration of Helsinki, including its amendments. All participants provided their written informed consent and were unaware of the experimental aims prior to data collection.

### 4.2 Participants

Twenty-eight healthy volunteers (13 female; 1 left-handed; mean age: 27.88 ± 4.46 years) were analysed. Eligibility was determined based on self-reported criteria: (1) no personal or family history of psychiatric, neurological or somatic disorders; (2) no chronic or acute respiratory diseases; (3) no use of central nervous system-affecting substances in the previous week; and (4) no previous experience with mindfulness-based meditation or breath control practices. All participants were of normal weight and had normal or corrected-to-normal vision.

### 4.3 Experimental procedure

As described in Zaccaro et al. (2024), the tasks were administered using the E-Prime 3.0 software (Psychology Software Tools, Pittsburgh, PA, USA). Participants seated comfortably in front of a computer screen positioned approximately 60 cm away for optimal keyboard use and screen visibility. Room lighting and noise levels were minimised, and temperature was maintained at a comfortable level. Written instructions were presented before each task. The experimenter then ensured that participants understood the instructions well and invited them to ask questions if necessary. EEG, ECG and respiratory signals were continuously recorded throughout the experiment.

The resting-state condition was recorded first. Participants sat for 10 minutes resting with their eyes open, looking at a fixation cross in the centre of the screen and allowing their thoughts to wander without focusing on anything in particular. They were also instructed to breathe spontaneously.

The cardiac interoceptive task used was the HCT (Pollatos et al., 2005; Schandry & Weitkunat, 1986). Participants were given the following instructions, adapted from Desmedt et al. (2018): “*In this task, direct your attention to your heart and the associated physical sensations. You are required to sustain your focus on your heart for various durations and count the number of heartbeats you perceive. Begin silently counting when the heart symbol appears on the screen. When the heart symbol disappears, report the number of heartbeats you are sure you felt. Only report the number of heartbeats you are sure about, without trying to estimate your heart rate*. *During each trial, keep your eyes open, gaze at the screen and avoid moving. Refrain from guiding your responses by checking your pulse in your wrists or neck. Breathe spontaneously and avoid changing your breathing frequency or holding your breath*”. The HCT consisted of three blocks of four randomised trials lasting 25, 35, 45 and 100 seconds (Ardizzi & Ferri, 2018; Di Lernia et al., 2018; Ueno et al., 2020), for a total of 12 trials. After each trial, participants reported the number of perceived heartbeats and rated their confidence (HCT confidence) on a 10-point Likert scale, ranging from 0 (very poor) to 9 (excellent). They did not receive any feedback on their performance.

The exteroceptive task used was the C-TCT. Participants were presented with digitally generated heartbeat sounds and instructed to count and report the number of heartbeats they heard (Wiebking et al., 2015). The C-TCT consisted of three blocks of four randomised trials lasting 25, 35, 45 and 100 seconds. The heartbeat sounds were presented at irregular intervals, with an average frequency of 60 bpm. After each trial, participants reported the number of heartbeats they heard and rated their confidence (C-TCT confidence). Before starting the experiment, participants completed a short training session and verbally confirmed that they could not perceive their heartbeat through the breathing belt to ensure that it was not too tight.

### 4.4 Electrophysiological recording and pre-processing

Respiratory activity was recorded using a respiratory belt placed around the chest (Respiratory Transducer TSD201, BIOPAC Systems Inc, Goleta, CA, USA). EEG signals were recorded using a 64-channel BrainAmp MR EEG system (BrainCap, BrainVision LLC, Garner, NC, USA) with electrodes positioned according to the International 10-20 system. Electrode impedance was kept below 10 kΩ for all channels. The ECG was recorded using three electrodes: two placed over the left costal margin and the right clavicle, with the ground electrode placed over the right costal margin (MP160, BIOPAC Systems Inc, Goleta, CA, USA). Three additional backup electrodes were integrated into the EEG recording system via an external electrode input box (BrainProducts GmbH, BrainVision LLC, Gilching, Germany) and placed over the left and right clavicles, with the ground at the left costal margin. The BrainAmp MR EEG, MP160 and TSD201 systems were synchronised with the E-Prime 3.0 software using a TriggerStation™ (BRAINTRENDS LTD 2010, Italy). All signals were sampled at 2.5 kHz with a .016 to 250 Hz bandpass filter and a 50 Hz notch filter. Signals were then filtered and down-sampled to 256 Hz.

Respiratory data were processed using custom MATLAB scripts (R2021a, MathWorks Inc, Natick, MA, USA). The respiratory signal was high-pass filtered at .1 Hz to remove baseline drift. Respiratory phases were detected using a validated method (Grund et al., 2022; Power et al., 2020). Outliers exceeding three scaled median absolute deviations from the local median within a 1-s moving window were linearly interpolated with the adjacent values. Data were then smoothed with a 1-s window filter (Savitzky & Golay, 1964) and z-scored. Inhalation and exhalation onsets were identified as local minima (troughs) and maxima (peaks), respectively. Peaks and troughs had to be at least 2 seconds apart, and the minimum prominence of the interquartile range of the z-scored data was multiplied by a factor ranging from .3 to .8, depending on the participant’s respiratory trace (Grund et al., 2022). A breath cycle was defined as the interval between successive inhalations (i.e. from one trough to the next).

EEG data were pre-processed using the EEGLAB toolbox (v2022.1; Delorme & Makeig, 2004). Data were filtered using a Hamming windowed FIR filter between .5 and 45 Hz (Al et al., 2020, 2021). EEG segments with poor signal quality were manually rejected. ICA was performed to remove artefacts related to heartbeat, eye and muscle activity (FastICA; Hyvärinen, 1999). The ECG, respiratory channels and excessively noisy channels were excluded from the ICA decomposition. A few noisy channels (less than 5), typically peripheral and not known to contribute to HEP generation (Coll et al., 2021), were identified. Independent components (ICs) were rejected based on IC time series, scalp maps, and power spectra (Delorme & Makeig, 2004). Special attention was given to the CFA and the pulse-related artifact (PA) as they significantly affect HEP activity (Coll et al., 2021). ICs contaminated by CFA and PA were identified by plotting R-peaks over IC time series, and ICs with activations time-locked to the ECG R-peak were removed (Zaccaro et al., 2024). The number of ICs removed ranged from 1 to 6 (3 ± 1 [mean ± SD]), with CFA-related ICs ranging from 0 to 3 (1 ± 1 [mean ± SD]). After ICA, previously identified noisy channels were interpolated using adjacent electrodes (Junghöfer et al., 2000). The cleaned EEG signals were then referenced to the mean reference (Antonacci et al., 2023; Barà et al., 2023; Zaccaro et al., 2024) as recommended (Candia-Rivera et al., 2021). ECG R-peaks were identified using the HEPLAB toolbox (Perakakis, 2019), with manual correction of mis-detected peaks (Al et al., 2021; Zaccaro et al., 2022). EEG data were epoched from -2 to 2 seconds around each R-peak (Kern et al., 2013). Epochs were averaged within participants for each task and were further classified based on the respiratory phase in which the corresponding R-peak occurred. This resulted in four epoch bins: HCT inhalation and exhalation, and C-TCT inhalation and exhalation.

### 4.5 Heartbeat-modulated cortical oscillations

Heartbeat-modulated cortical oscillations were calculated using the FieldTrip toolbox (version 20231025; Oostenveld et al., 2011) according to established procedures (Park et al., 2018). Time-frequency analysis was performed on the time-series data for each experimental condition. For each trial, we convolved the signal with a complex wavelet constructed by pointwise multiplying the real cosine and imaginary sine components at each frequency with a Hann taper. Both the data and the tapered wavelet were then Fourier transformed, and their element-wise multiplication in the frequency domain, followed by an inverse Fourier transform, yielded the convolution in the time domain. A sliding window with a Hann taper was used, maintaining a constant number of cycles per time window for all frequencies (4 cycles per time window). Zero padding was applied to round the maximum trial length to the nearest power of 2. Power was then extracted from the resulting complex representation by computing the squared magnitude of the signal. Time-frequency analysis was performed on each EEG channel from 1 Hz to 45 Hz, with a step size of .5 Hz, and from -1.5 s to 2 s after the R-peak, with steps of .01 sec. The mean power value of the pre-R-peak time window (-300 msec to -100 msec) was designated as the baseline and used to normalise the post-R-peak power values in each frequency bin (relative heartbeat-related power; Park et al., 2018). This baseline normalisation was intended to prevent smearing of post-R-peak activity into baseline and to mitigate the potential influence of CFA around the ECG P-wave (Park et al., 2018).

To compute heartbeat-related ITC, we first normalised the complex Fourier coefficients by their magnitude to focus on the phase information. To derive the average ITC, the resulting phase information was summed and normalised by the number of trials. ITC reflects the consistency of phase across multiple trials at different times and frequencies, with values ranging from 0 to 1, with higher values indicating more consistent phase alignment across trials. Similar to power values, we used the baseline window to normalise ITC values (relative heartbeat-related ITC; Park et al., 2018). To remove components reflecting PA and its harmonics, we excluded frequency bands below 4 Hz (Kern et al., 2013; Park et al., 2018).

To extract the heartbeat-related FC, we computed the complex Fourier spectra from 1 Hz to 45 Hz (all settings were the same as for power calculation). Fourier spectra were calculated between -300 and -100 msec before the R-peak (“baseline”) and between 300 and 600 msec after the R-peak (“active” period of heartbeat effects; (Liuzzi et al., 2024; Zaccaro et al., 2024). The iCOH was used as a measure of undirected FC between electrodes in the sensor space, resulting in a connectivity matrix between all pairs of electrodes. The iCOH suppresses spurious coherence due to electromagnetic field propagation, which is indicated by zero-phase synchronisation between sensors (Nolte et al., 2004). We hypothesised that FC would change in the theta (4-8 Hz) and alpha (8-13 Hz) bands, according to previous studies (Fló et al., 2024; Kern et al., 2013; Kim & Jeong, 2019). Correlation matrices were calculated across trials, resulting in a correlation matrix for each participant, task (HCT and C-TCT), cardiac time window (baseline and active), respiratory phase (inhalation and exhalation), and frequency band (alpha and theta). To assess heartbeat-related FC, we calculated the ratio of connectivity during the active cardiac time window to that of the baseline cardiac time window (relative heartbeat-related FC).

To extract undirected graph connectivity metrics, we thresholded the iCOH matrices using a proportional threshold that preserves a percentage of the strongest matrix weights (Perl et al., 2019). In graph theory, a network is represented by nodes (each EEG channel) and edges indicating the strength of the connection (the iCOH values for each pair of EEG channels). We used six incremental thresholds ranging from 5% to 30% as cutoffs (e.g., a threshold of 5% means that out of 100 nodes, each node is connected to five other nodes on average) (Perl et al., 2019). All edges that did not reach the threshold were removed. We extracted the following network connectivity measures for each participant, threshold, task, cardiac time window, respiratory phase, and frequency band using the brain connectivity toolbox (Rubinov & Sporns, 2010): i) density: the ratio of current connections to all possible connections, reflecting the overall interconnectivity of the network; ii) clustering coefficient: the ratio of triangles formed around each node, illustrating the degree of dense interconnectivity among neighbouring nodes; iii) global efficiency: a measure of network integration that reflects the network’s ability to facilitate long-range information transfer; and iv) local efficiency: a measure of the network’s capacity to support short-range information transfer and its resilience to incremental disconnections.

### 4.6 Statistical analyses for heartbeat-modulated cortical oscillations

Changes in heartbeat-modulated cortical oscillations were assessed using the FieldTrip toolbox (v.20231025; Oostenveld et al., 2011). The analysis focused on the cluster mass rather than individual electrodes or time points. Initially, spatial (neighbouring channels), frequency (adjacent frequency bins), and temporal (adjacent time points) points with a p-value below .05 were included in a cluster. A cluster statistic was calculated by summing the t-values within each cluster. The null distribution was estimated by Monte Carlo permutations, with condition labels randomly shuffled 5,000 times. The largest cluster-level statistic from each randomisation was included in the null distribution. Statistical significance was determined by comparing the experimentally observed cluster-level statistics with the randomly generated null distribution. Clusters exceeding the 1 – α percentile of this distribution (two-tailed) were considered significant. The minimum number of adjacent channels required to form a cluster was set to 2. To test for interaction effects, we used the permutation of residuals method (Anderson & Ter Braak, 2003; Still & White, 1981; Winkler et al., 2014). For instance, to test the 2×2 interaction (task × respiratory phase), we calculated the difference between the two levels of phase for each level of task and conducted paired cluster-based permutation t-tests comparing difference values (HCT-exhalation minus HCT-inhalation vs. C-TCT-exhalation minus C-TCT-inhalation). Based on Zaccaro et al., (2024), tests were performed at each electrode and time point for the whole late time window (300-600 msec) following the R-peak. We tested each frequency band of interest separately, averaging over their respective frequency bins: theta (4-8 Hz), alpha (8-13 Hz), beta (13-30 Hz), and low-gamma (30-45 Hz). We excluded external electrodes (F7, F8, FT7, FT8, T7, T8, TP7, TP8, P7, P8, PO7, PO8, O1, O2, Oz, TP9, and TP10), which are prone to noise contamination, in line with a recent meta-analysis (Coll et al., 2021). Cluster-based permutation t-tests were conducted on the time-frequency data, with heartbeat-related power and ITC values as dependent variables. Planned comparisons included HCT exhalation vs. HCT inhalation, and C-TCT exhalation vs. C-TCT inhalation (Zaccaro et al., 2024).

To examine the interaction effects between tasks and respiratory phases on heartbeat-related FC, we applied the same permutation-of-residuals method used for power and ITC. We excluded from the FC analysis the same external electrodes omitted in the power and ITC analyses, as they are particularly susceptible to noise contamination. Relative iCOH values between electrode pairs for each frequency band (theta and alpha) were used as the dependent variable. Planned permutation-based t-tests were performed for both the theta and alpha bands, comparing HCT exhalation vs. HCT inhalation, and C-TCT exhalation vs. C-TCT inhalation. Significance was assessed using a single-threshold permutation test for the maximum t-statistics (5,000 permutations). Connectivity values showing a significant effect with a p-value below .01 (uncorrected) were used to construct symmetric connectivity matrices for theta and alpha using the BrainNet Viewer toolbox (Xia et al., 2013).

Undirected graph connectivity metrics were analysed using a series of LMEMs. The models were implemented with the general analyses for the linear model (GAMLj) module in jamovi (v2.3.21; the jamovi project, 2022; Gallucci, 2019), using the restricted maximum likelihood (REML) estimation method. Fixed factors included task (HCT and C-TCT), respiratory phase (inhalation and exhalation), cardiac time window (baseline and active), and threshold (5%, 10%, 15%, 20%, 25%, and 30%). Degrees of freedom were estimated using the Satterthwaite method. The full model incorporated a three-way interaction (task × time × phase) across all graph theory metrics, along with a random intercept and random slopes for the interaction at the participant level. For each model parameter, we reported the estimated coefficient (b), standard error (SE), t-statistic, and p-value. While model selection is often guided by statistical fit indices, the inclusion of the three-way interaction was theoretically justified to enable planned contrasts, revealing condition-specific effects that may not be apparent in the omnibus interaction term. Planned t-tests were conducted separately for each task and respiratory phase and corrected for multiple comparisons using the Benjamini-Yekutieli false discovery rate (FDR) method. The comparisons were as follows: HCT baseline inhalation vs. HCT baseline exhalation; HCT active inhalation vs. HCT active exhalation; HCT baseline inhalation vs. HCT active inhalation; HCT baseline exhalation vs. HCT active exhalation; C-TCT baseline inhalation vs. C-TCT baseline exhalation; C-TCT active inhalation vs. C-TCT active exhalation; C-TCT baseline inhalation vs. C-TCT active inhalation; C-TCT baseline exhalation vs. C-TCT active exhalation (number of planned t-tests: 8).

We verified that changes in alpha and theta heartbeat-related power did not reflect an artificial increase induced by CFA changes by testing the interaction between tasks and respiratory phases on the time-frequency decomposed ECG signal, time-locked to the R-peak (Daoui et al., 2022; Tsutsumi et al., 2014). This was done using a repeated-measures, FDR-corrected two-tailed t-test within the time window and frequency bands where significant power differences were observed. We also ensured that any significant changes in iCOH did not represent an artificial increase in phase synchronisation caused by CFA. Following the procedure outlined in Kim et al. (2019), we tested whether the relative alpha and theta iCOH increased during exhalation compared to inhalation between the ECG and EEG electrodes that showed significant iCOH changes, using permutation-based dependent samples t-tests.

### 4.7 Relationships between heartbeat-modulated cortical oscillations and heartbeat-evoked potential

Based on prior research (Crivelli & Balconi, 2025; Kritzman et al., 2022; Luft & Bhattacharya, 2015; Mccraty et al., 2009; Rominger et al., 2025; Villena-González et al., 2017), we assessed relationships between changes in HEP amplitude across respiratory phases and corresponding differences in heartbeat-related alpha power during HCT. Single-trial epochs around each R peak (from -300 to 600 msec) were filtered between .5 and 3 Hz (Studenova et al., 2023). To assess the spatial overlap between the distributions of significant heartbeat-related alpha power and HEP amplitude changes across respiratory phases, a full-scalp cluster-based permutation t-test was performed, using HEP amplitude as the dependent variable, comparing inhalation and exhalation in the late time window (300-600 msec after the R-peak; Zaccaro et al., 2024). HEP amplitude and heartbeat-related alpha power values were averaged for each participant, and Δ values between respiratory phases were extracted. Mean values were obtained by averaging across the time windows that previously showed significant power differences between respiratory phases. We computed Spearman’s rho and p-values for each channel (α = .05, two-tailed) by correlating participant-level mean heartbeat-related Δalpha power with the corresponding mean ΔHEP values. We applied the FDR correction procedure to control for multiple comparisons across channels. The analysis was repeated for other frequency bands during both HCT and C-TCT, and we also assessed correlations between ΔHEP amplitude and ΔITC, as well as between Δpower and ΔITC across all frequency bands. Finally, we examined correlations between HEP, power, and ITC across all frequency bands during HCT and C-TCT by analysing their absolute values, rather than Δ values. We computed the instantaneous group-level correlation between Δalpha power and the corresponding ΔHEP values for each time point and frequency bin, yielding a time-frequency representation of Spearman’s rho and p-values (α = .05, two-tailed). The analysis focused on channels that had previously shown a significant participant-level correlation between ΔHEP amplitude and Δalpha power. We compared instantaneous correlations between HCT and C-TCT by applying Fisher’s z-normalization to Spearman’s rho values averaged across the previously selected channels. Statistical significance at each time point was determined using the cumulative distribution function of the standard normal distribution to evaluate the probability of observing extreme z-differences under the null hypothesis. We applied the FDR correction procedure to control for multiple comparisons across time points.

To examine the relationships between HCT/C-TCT trial-level changes in HEP, heartbeat-related power and ITC across respiratory phases, as well as their associations with accuracy and cardiac physiology, we applied multiple LMEMs. For each HCT and C-TCT trial (12 trials per participant), we computed the changes across respiratory phases (Δ) in HEP, heartbeat-related power, and ITC within the alpha band, along with accuracy and confidence. HCT/C-TCT trial-level Δ values were derived by first extracting the mean values from significant channels and time windows during exhalation and inhalation and then calculating the Δ (exhalation minus inhalation). To account for the influence of cardiac physiology, we also computed ΔHR and ΔECG. ΔHR and ΔECG were derived by subtracting the HCT/C-TCT trial-level mean HR and mean ECG amplitude during inhalation from that during exhalation, respectively. LMEMs were implemented with the general analyses for the linear model (GAMLj) module in jamovi (v2.3.21; the jamovi project, 2022; Gallucci, 2019), using the restricted maximum likelihood (REML) method. We computed multiple models, starting from the simplest model and then adding factors stepwise. The Akaike information criterion (AIC) was used to determine the best-fitting model. Degrees of freedom were estimated using the Satterthwaite method. For each parameter of the best-fitting model, we reported the estimated coefficient (b), standard error (SE), t-statistic and p-value. We fitted four separate LMEMs, with ΔHEP and accuracy as dependent variables, respectively for HCT and C-TCT.

## Acknowledgments

We sincerely thank Isabella Pohle-Kaupp for her valuable support in the implementation and interpretation of linear mixed-effects models.

## Author contributions

Andrea Zaccaro: conceptualization, methodology, software, validation, formal analysis, investigation, data curation, visualization, writing original draft, and writing-review and editing. Francesca della Penna: investigation, formal analysis, data curation, and writing-review and editing. Francesco Bubbico: software, data curation, and writing-review and editing. Başak Bayram: software, data curation, and writing-review and editing. Eleonora Parrotta: software, data curation, and writing-review and editing. Mauro Gianni Perrucci: methodology, software, data curation, resources, and writing-review and editing. Marcello Costantini: supervision, funding acquisition, and writing-review and editing. Francesca Ferri: conceptualization, methodology, resources, supervision, project administration, funding acquisition, visualization, and writing-review and editing.

## Data and code availability

The code used for experiment analysis, along with the raw behavioural and physiological data, will be made available upon publication.

