## Supplementary material for "Cardio-respiratory interactions in interoceptive perception: The role of heartbeat-modulated cortical oscillations"

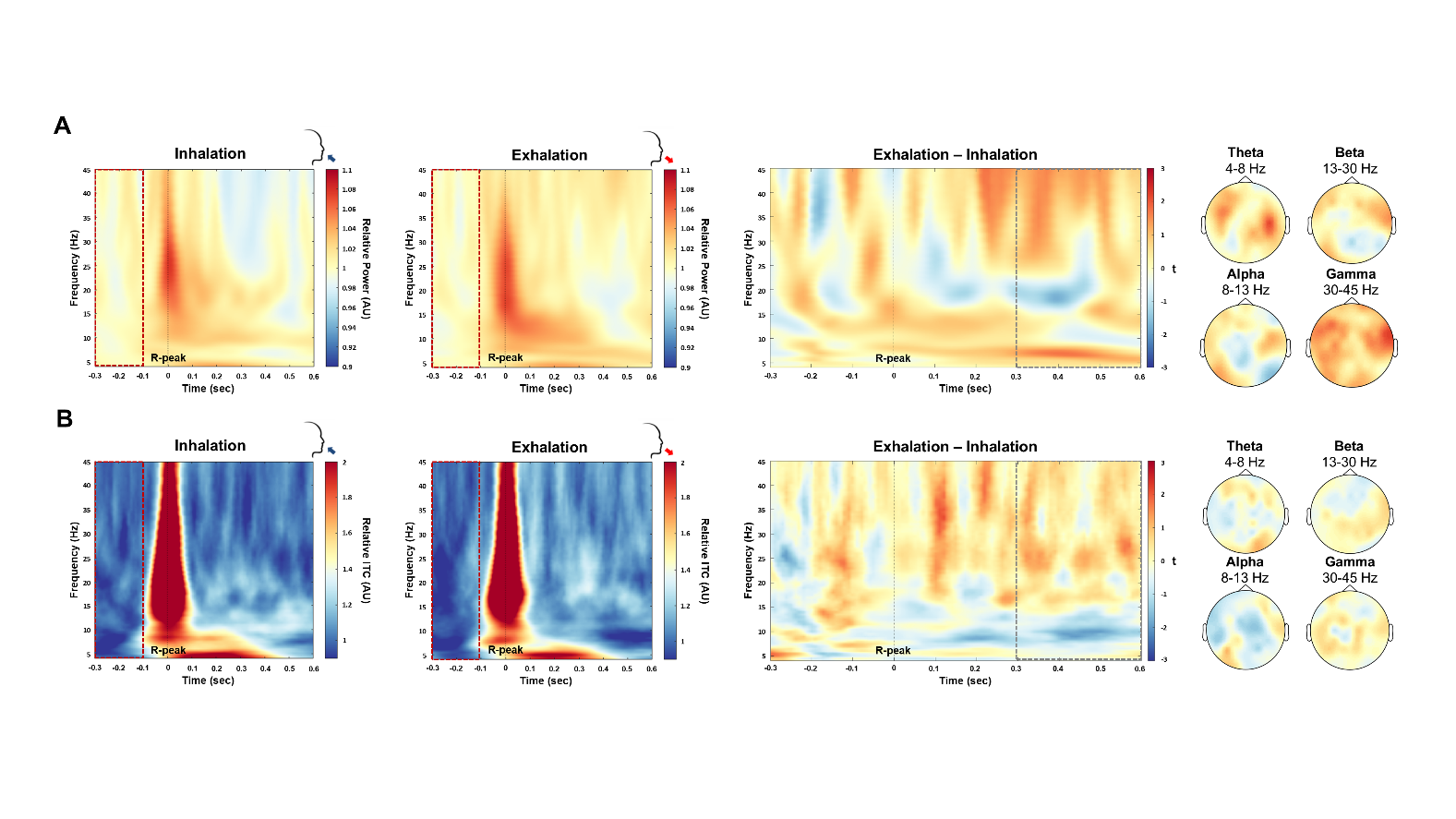

**Supplementary Figure 1. Heartbeat-related power and inter-trial coherence changes across respiratory phases during the resting-state.** (A) Grand-averaged heartbeat-related power during the resting-state during inhalation (left), exhalation (middle), and the exhalation-minus-inhalation difference (right). Topographical distributions illustrate non-significant differences. (B) Grand-averaged heartbeat-related inter-trial coherence during the resting-state during inhalation (left), exhalation (middle), and the exhalation-minus-inhalation difference (right). Topographical distributions illustrate non-significant differences. The red dotted area represents the baseline window (-300 to -100 msec relative to R-peak onset). The grey dotted area marks the temporal window of interest for statistical analysis (300-600 msec after the R-peak). Abbreviations: ITC = inter-trial coherence, AU = arbitrary unit.

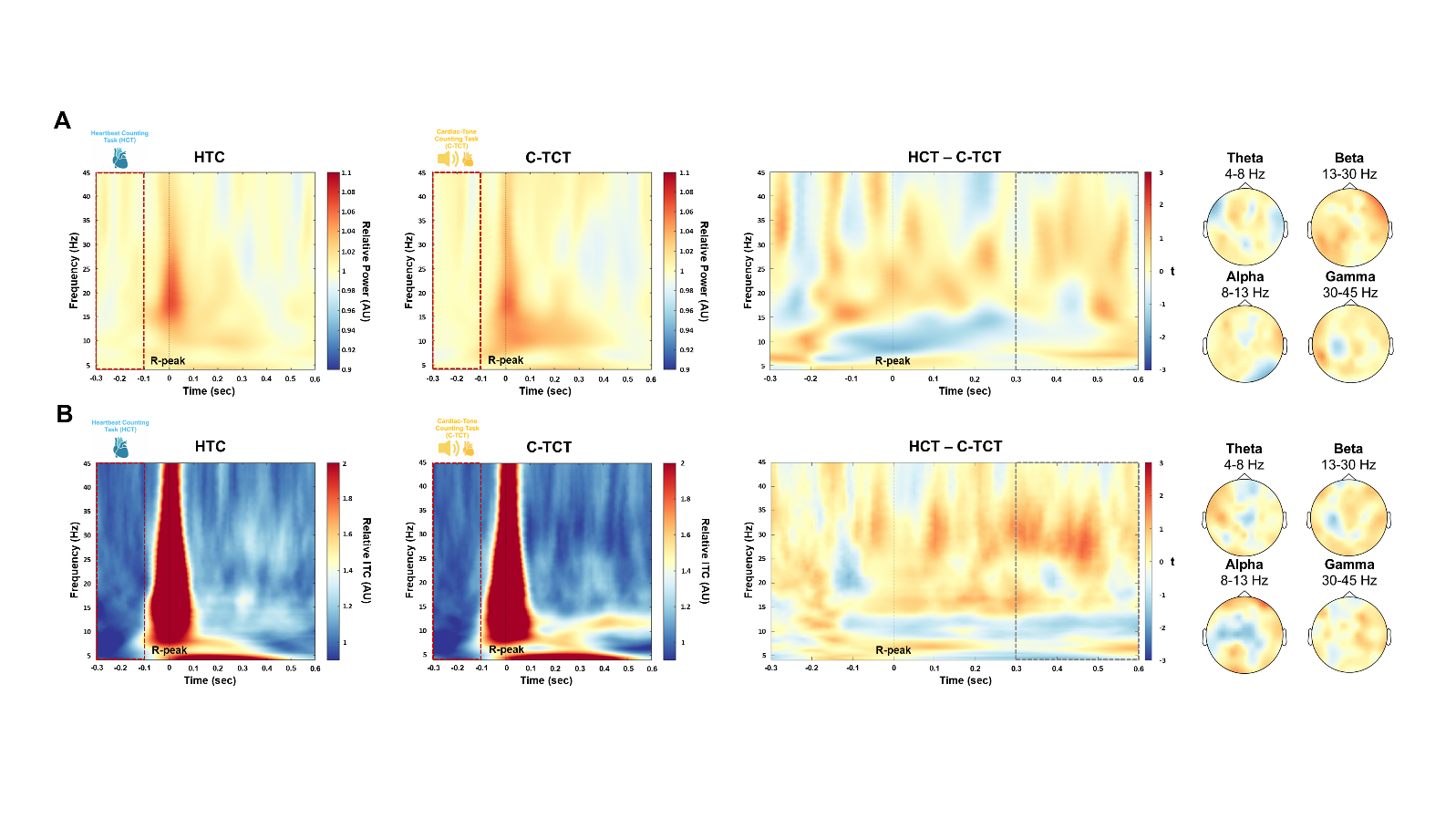

**Supplementary Figure 2. Heartbeat-related power and inter-trial coherence changes across tasks without considering respiratory phases.** (A) Grand-averaged heartbeat-related power during the heartbeat counting task (left), cardiac-tone counting task (middle), and their difference (right). Topographical distributions illustrate non-significant differences. (B) Grand-averaged heartbeat-related inter-trial coherence during the heartbeat counting task (left), cardiac-tone counting task (middle), and their difference (right). Topographical distributions illustrate non-significant differences. The red dotted area represents the baseline window (-300 to -100 msec relative to R-peak onset). The grey dotted area marks the temporal window of interest for statistical analysis (300-600 msec after the R-peak). Abbreviations: HCT = heartbeat counting task, C-TCT = cardiac-tone counting task, ITC = inter-trial coherence, AU = arbitrary unit.

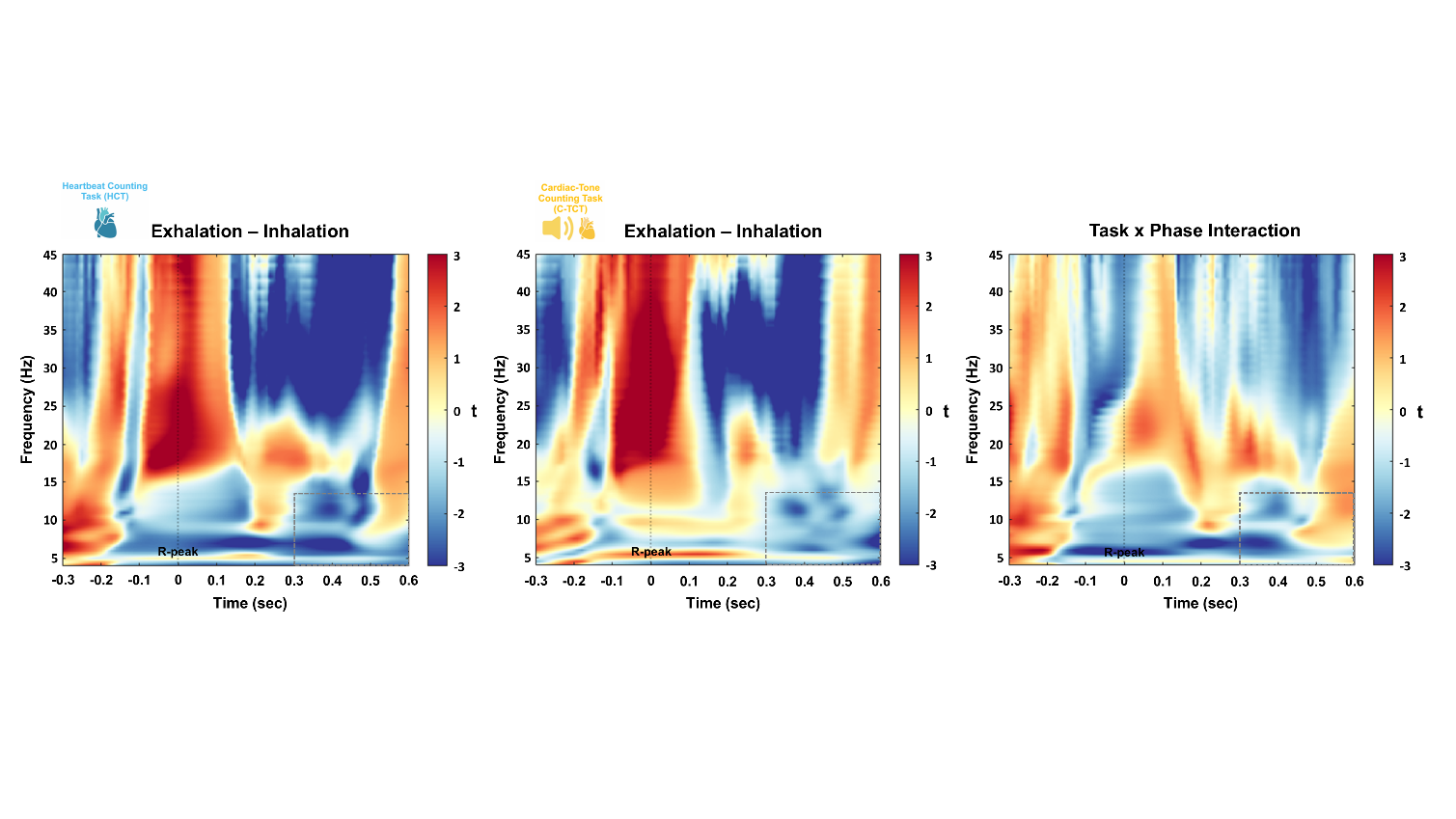

**Supplementary Figure 3. ECG signal power changes across respiratory phases and tasks.** Grand-averaged ECG signal power changes between respiratory phases (exhalation vs. inhalation) during the heartbeat counting task (left), the cardiac-tone counting task (middle), and their task × phase interaction (right). The grey dotted area marks the temporal window of interest for statistical analysis (300-600 msec after the R-peak, within the 4–13 Hz frequency range).

| **Channels** | **t** | **p** |
| --- | --- | --- |
| Fp1-P1 | 1.0639 | 0.0036 |
| Fp2-FCz | -3.0062 | 0.0014 |
| F4-P6 | 2.6667 | 0.0072 |
| C3-P3 | 1.7642 | 0.0018 |
| C3-CP6 | 2.1115 | 0.0064 |
| C4-CPz | 2.9565 | 0.0010 |
| P4-CP1 | 2.3860 | 0.0050 |
| P4-POz | 2.2980 | 0.0100 |
| Fz-PO3 | -2.5047 | 0.0054 |
| Cz-F2 | -2.1558 | 0.0002 |
| FC1-F5 | -1.6780 | 0.0046 |
| FC1-F6 | -2.4412 | 0.0090 |
| CP2-C6 | 1.4629 | 0.0056 |
| FC5-F6 | 2.3034 | 0.0082 |
| F2-AF3 | -2.6663 | 0.0054 |
| F2-AF4 | -1.5709 | 0.0074 |
| F2-FCz | -1.0245 | 0.0004 |
| C2-AF3 | -2.7037 | 0.0048 |
| AF4-FCz | -2.3720 | 0.0014 |
| PO3-P6 | 2.9372 | 0.0016 |
| F6-Fpz | -2.5218 | 0.0004 |

**Supplementary Table 1. Heartbeat-related functional connectivity changes across respiratory phases and tasks in the theta band.** T-statistics and p-values are reported for channel pairs exhibiting a significant (p < .01, uncorrected) task × phase interaction in the theta band.

| **Channels** | **t** | **p** |
| --- | --- | --- |
| Fp1-PO4 | 2.2274 | 0.0084 |
| Fp1-P5 | 2.5921 | 0.0050 |
| Fp2-FCz | -1.5365 | 0.0074 |
| F3-Pz | 1.3712 | 0.0082 |
| F4-P4 | 1.4461 | 0.0092 |
| C3-CP6 | 1.8603 | 0.0062 |
| C4-PO4 | 2.2560 | 0.0098 |
| C4-CPz | 3.0402 | 0.0020 |
| P3-P4 | 2.2167 | 0.0076 |
| P3-AF3 | 1.8756 | 0.0004 |
| P4-Fz | 1.5843 | 0.0046 |
| P4-Fpz | 1.1767 | 0.0026 |
| Cz-PO3 | 2.0234 | 0.0074 |
| Pz-F1 | 2.0303 | 0.0070 |
| FC1-CP4 | 2.3402 | 0.0100 |
| CP5-AF3 | -2.2601 | 0.0088 |
| F1-FC3 | 2.1905 | 0.0072 |
| AF4-CP4 | 2.8005 | 0.0006 |
| AF4-PO4 | 1.7049 | 0.0068 |
| FC3-CP4 | 2.7218 | 0.0066 |
| FC4-F6 | -2.0105 | 0.0034 |
| PO3-P6 | 2.3152 | 0.0096 |
| F6-Fpz | -2.0017 | 0.0076 |
| F6-POz | 1.6183 | 0.0034 |
| P6-POz | 2.2161 | 0.0010 |

**Supplementary Table 2. Heartbeat-related functional connectivity changes across respiratory phases during the heartbeat counting task in the theta band.** T-statistics and p-values are reported for channel pairs exhibiting significant (p < .01, uncorrected) differences between exhalation and inhalation in the theta band.

| **Channels** | **t** | **p** |
| --- | --- | --- |
| Fp1-P5 | 2.2113 | 0.0072 |
| Fp2-C3 | -2.3723 | 0.0076 |
| F4-Fz | 2.2906 | 0.0066 |
| C3-F6 | -2.2969 | 0.0088 |
| P4-P2 | -2.0752 | 0.0020 |
| Fz-C1 | 1.4491 | 0.0060 |
| Fz-PO3 | 2.8650 | 0.0006 |
| Cz-F2 | 1.5592 | 0.0056 |
| AFz-PO3 | 1.3346 | 0.0064 |
| FC1-C6 | 2.4685 | 0.0040 |
| F2-CP4 | 2.3480 | 0.0092 |
| F2-PO4 | -2.3669 | 0.0036 |
| F2-FCz | 2.3177 | 0.0004 |
| C1-P2 | -2.3253 | 0.0084 |
| C2-AF3 | 2.3567 | 0.0050 |
| C2-P5 | 2.1003 | 0.0034 |
| P1-FC4 | 2.0144 | 0.0072 |
| AF4-FCz | 2.0729 | 0.0050 |
| F6-Fpz | 1.3493 | 0.0032 |

**Supplementary Table 3. Heartbeat-related functional connectivity changes across respiratory phases during the cardiac-tone counting task in the theta band.** T-statistics and p-values are reported for channel pairs exhibiting significant (p < .01, uncorrected) differences between exhalation and inhalation in the theta band.

| **Channels** | **t** | **p** |
| --- | --- | --- |
| Fp1-Fp2 | -1.6513 | 0.0096 |
| Fp2-FC6 | 2.4993 | 0.0050 |
| Fp2-CP4 | -2.5038 | 0.0072 |
| F3-C3 | -2.2806 | 0.0080 |
| F4-CP2 | -2.7027 | 0.0050 |
| F4-C6 | 2.4396 | 0.0002 |
| C3-F5 | -2.3810 | 0.0032 |
| P4-CP2 | -2.8421 | 0.0006 |
| Fz-C6 | 3.1867 | 0.0010 |
| FC2-C6 | 2.4970 | 0.0022 |
| CP2-C1 | 1.3770 | 0.0012 |
| FC6-C6 | 2.5168 | 0.0038 |
| CP6-C2 | 2.3217 | 0.0030 |
| F1-F6 | -1.4752 | 0.0094 |
| F2-PO4 | -2.6374 | 0.0050 |
| F2-C6 | 1.9080 | 0.0076 |
| CP3-F5 | -1.7434 | 0.0018 |
| CP4-CPz | 2.9836 | 0.0030 |
| F6-P5 | 2.9873 | 0.0020 |
| C6-POz | -2.1920 | 0.0092 |
| Fpz-POz | -2.1106 | 0.0048 |

**Supplementary Table 4. Heartbeat-related functional connectivity changes across respiratory phases and tasks in the alpha band.** T-statistics and p-values are reported for channel pairs exhibiting a significant (p < .01, uncorrected) task × phase interaction in the alpha band.

| **Channels** | **t** | **p** |
| --- | --- | --- |
| Fp2-C6 | 2.7914 | 0.0014 |
| F4-C3 | 1.3470 | 0.0046 |
| F4-C6 | 1.9252 | 0.0002 |
| P4-CP2 | -2.4151 | 0.0030 |
| Cz-C6 | 1.2626 | 0.0040 |
| Pz-C1 | 1.1550 | 0.0078 |
| AFz-C6 | 2.5774 | 0.0018 |
| CP1-F2 | 2.0631 | 0.0088 |
| FC6-C6 | 2.1268 | 0.0082 |
| C1-P1 | 1.5225 | 0.0062 |
| C2-AF4 | -1.5986 | 0.0050 |
| AF4-C6 | 1.0433 | 0.0064 |
| CP3-F5 | -2.4871 | 0.0084 |
| C6-FCz | 2.1529 | 0.0040 |

**Supplementary Table 5. Heartbeat-related functional connectivity changes across respiratory phases during the heartbeat counting task in the alpha band.** T-statistics and p-values are reported for channel pairs exhibiting significant (p < .01, uncorrected) differences between exhalation and inhalation in the alpha band.

| **Channels** | **t** | **p** |
| --- | --- | --- |
| Fp1-AFz | -1.4756 | 0.0016 |
| Fp1-F2 | -2.8644 | 0.0002 |
| Fp1-C6 | -2.2656 | 0.0036 |
| Fp2-FC6 | -2.2117 | 0.0068 |
| Fp2-F2 | -2.1386 | 0.0084 |
| C4-FC5 | -1.1679 | 0.0100 |
| P3-Cz | -1.7612 | 0.0046 |
| P3-CP1 | -1.1166 | 0.0028 |
| Fz-AF4 | -2.2418 | 0.0010 |
| Fz-C6 | -2.2673 | 0.0048 |
| Cz-C2 | -2.5111 | 0.0032 |
| FC2-C6 | -2.2149 | 0.0066 |
| CP1-CP6 | 2.2169 | 0.0092 |
| FC5-FC6 | 2.6170 | 0.0046 |
| F2-C6 | -1.9814 | 0.0078 |
| FC3-FCz | -2.5262 | 0.0040 |
| C6-POz | 2.3036 | 0.0036 |

**Supplementary Table 6. Heartbeat-related functional connectivity changes across respiratory phases during the cardiac-tone counting task in the alpha band.** T-statistics and p-values are reported for channel pairs exhibiting significant (p < .01, uncorrected) differences between exhalation and inhalation in the alpha band.

|  | **Estimate** | **SE** | **t** | **p** |
| --- | --- | --- | --- | --- |
| Intercept | 0.2475 | 0.0056 | 44.024 | 1.12e-26 |
| Attention | 0.0378 | 0.0189 | 1.997 | 0.0560 |
| Time | 0.0364 | 0.0151 | 2.402 | 0.0234 |
| Phase | -0.0097 | 0.0135 | -0.719 | 0.4785 |
| Threshold 2 - 1 | 0.0528 | 0.0018 | 28.874 | 2.42e-137 |
| Threshold 3 - 1 | 0.0948 | 0.0018 | 51.868 | 1.75e-299 |
| Threshold 4 - 1 | 0.1323 | 0.0018 | 72.319 | 0.0000 |
| Threshold 5 - 1 | 0.1647 | 0.0018 | 90.042 | 0.0000 |
| Threshold 6 - 1 | 0.1970 | 0.0018 | 107.722 | 0.0000 |
| Attention * Time | 0.0029 | 0.0156 | 0.189 | 0.8514 |
| Attention * Phase | -0.0552 | 0.0206 | -2.671 | 0.0126 |
| Time * Phase | 0.0387 | 0.0258 | 1.501 | 0.1449 |
| Attention * Time * Phase | 0.0413 | 0.0623 | 0.664 | 0.5124 |

**Supplementary Table 7. Parameter estimates of the linear mixed-effects model for theta-band graph density.** Abbreviations: SE = standard error.

|  | **Estimate** | **SE** | **t** | **p** |
| --- | --- | --- | --- | --- |
| Intercept | 0.2785 | 0.0099 | 28.088 | 1.60e-21 |
| Attention | 0.0355 | 0.0225 | 1.580 | 0.1257 |
| Time | 0.0397 | 0.0143 | 2.781 | 0.0097 |
| Phase | -0.0075 | 0.0139 | -0.540 | 0.5936 |
| Threshold 2 - 1 | 0.0613 | 0.0027 | 22.395 | 5.05e-92 |
| Threshold 3 - 1 | 0.1079 | 0.0027 | 39.396 | 2.18e-213 |
| Threshold 4 - 1 | 0.1496 | 0.0027 | 54.584 | 3.19e-317 |
| Threshold 5 - 1 | 0.1845 | 0.0027 | 67.308 | 0.0000 |
| Threshold 6 - 1 | 0.2191 | 0.0027 | 79.948 | 0.0000 |
| Attention * Time | 0.0225 | 0.0167 | 1.347 | 0.1893 |
| Attention * Phase | -0.0509 | 0.0220 | -2.307 | 0.0289 |
| Time * Phase | 0.0233 | 0.0241 | 0.967 | 0.3420 |
| Attention * Time * Phase | 0.0491 | 0.0641 | 0.767 | 0.4499 |

**Supplementary Table 8. Parameter estimates of the linear mixed-effects model for theta-band graph clustering coefficient.** Abbreviations: SE = standard error.

|  | **Estimate** | **SE** | **t** | **p** |
| --- | --- | --- | --- | --- |
| Intercept | 0.4639 | 0.0157 | 29.403 | 4.8e-22 |
| Attention | 0.0567 | 0.0333 | 1.702 | 0.1001 |
| Time | 0.0648 | 0.0228 | 2.834 | 0.0085 |
| Phase | -0.0057 | 0.0197 | -0.291 | 0.7730 |
| Threshold 2 - 1 | 0.1073 | 0.0052 | 20.630 | 2.3e-80 |
| Threshold 3 - 1 | 0.1835 | 0.0052 | 35.257 | 1.5e-183 |
| Threshold 4 - 1 | 0.2467 | 0.0052 | 47.407 | 1.6e-269 |
| Threshold 5 - 1 | 0.2954 | 0.0052 | 56.771 | 0.0000 |
| Threshold 6 - 1 | 0.3404 | 0.0052 | 65.405 | 0.0000 |
| Attention * Time | 0.0243 | 0.0242 | 1.004 | 0.3244 |
| Attention * Phase | -0.0577 | 0.0331 | -1.743 | 0.0927 |
| Time * Phase | 0.0431 | 0.0384 | 1.121 | 0.2721 |
| Attention * Time * Phase | 0.0642 | 0.0991 | 0.648 | 0.5224 |

**Supplementary Table 9. Parameter estimates of the linear mixed-effects model for theta-band graph local efficiency.** Abbreviations: SE = standard error.

|  | **Estimate** | **SE** | **t** | **p** |
| --- | --- | --- | --- | --- |
| Intercept | 0.5378 | 0.0119 | 45.099 | 5.87e-27 |
| Attention | 0.0456 | 0.0228 | 1.993 | 0.0564 |
| Time | 0.0476 | 0.0190 | 2.503 | 0.0186 |
| Phase | -0.0035 | 0.0138 | -0.2526 | 0.8025 |
| Threshold 2 - 1 | 0.0891 | 0.0039 | 22.640 | 1.11e-93 |
| Threshold 3 - 1 | 0.1472 | 0.0039 | 37.383 | 6.31e-199 |
| Threshold 4 - 1 | 0.1909 | 0.0039 | 48.470 | 9.55e-277 |
| Threshold 5 - 1 | 0.2228 | 0.0039 | 56.587 | 0.0000 |
| Threshold 6 - 1 | 0.2517 | 0.0039 | 63.918 | 0.0000 |
| Attention * Time | 0.0047 | 0.0175 | 0.269 | 0.7900 |
| Attention * Phase | -0.0325 | 0.0235 | -1.382 | 0.1781 |
| Time * Phase | 0.0435 | 0.0277 | 1.568 | 0.1284 |
| Attention * Time * Phase | 0.0032 | 0.0668 | 0.048 | 0.9614 |

**Supplementary Table 10. Parameter estimates of the linear mixed-effects model for theta-band graph global efficiency.** Abbreviations: SE = standard error.

| **Comparison** | **Difference** | **SE** | **t** | **p** | **p_FDR_** |
| --- | --- | --- | --- | --- | --- |
| C-TCT-active-exhalation vs. C-TCT-active-inhalation | -0.0088 | 0.0255 | -0.3460 | 0.7320 | 0.732 |
| C-TCT-active-exhalation vs. C-TCT-baseline-exhalation | -0.0259 | 0.0274 | -0.9466 | 0.3522 | 0.4697 |
| C-TCT-active-inhalation vs. C-TCT-baseline-inhalation | -0.0440 | 0.0262 | -16.816 | 0.1041 | 0.4167 |
| C-TCT-baseline-exhalation vs. C-TCT-baseline-inhalation | -0.0268 | 0.0284 | -0.9478 | 0.3516 | 0.4697 |
| HCT-active-exhalation vs. HCT-active-inhalation | 0.0670 | 0.0251 | 2.6717 | 0.0126 | **0.0253** |
| HCT-active-exhalation vs. HCT-baseline-exhalation | -0.0082 | 0.0317 | -0.2605 | 0.7964 | 0.7964 |
| HCT-active-inhalation vs. HCT-baseline-inhalation | -0.0676 | 0.0190 | -3.5660 | 0.0013 | **0.0055** |
| HCT-baseline-exhalation vs. HCT-baseline-inhalation | 0.0076 | 0.0267 | 0.2862 | 0.7769 | 0.7964 |

**Supplementary Table 11. Planned comparisons of theta-band graph density.** Abbreviations: HCT = heartbeat counting task, C-TCT = cardiac-tone counting task, SE = standard error, FDR = false discovery rate.

| **Comparison** | **Difference** | **SE** | **t** | **p** | **p_FDR_** |
| --- | --- | --- | --- | --- | --- |
| C-TCT-active-exhalation vs. C-TCT-active-inhalation | -0.0185 | 0.0286 | -0.648 | 0.5223 | 0.5384 |
| C-TCT-active-exhalation vs. C-TCT-baseline-exhalation | -0.0291 | 0.0254 | -1.148 | 0.2609 | 0.5384 |
| C-TCT-active-inhalation vs. C-TCT-baseline-inhalation | -0.0279 | 0.0257 | -1.085 | 0.2876 | 0.5384 |
| C-TCT-baseline-exhalation vs. C-TCT-baseline-inhalation | -0.0172 | 0.0277 | -0.623 | 0.5384 | 0.5384 |
| HCT-active-exhalation vs. HCT-active-inhalation | 0.0569 | 0.0239 | 2.383 | 0.0244 | **0.0490** |
| HCT-active-exhalation vs. HCT-baseline-exhalation | -0.0271 | 0.0324 | -0.836 | 0.4105 | 0.5473 |
| HCT-active-inhalation vs. HCT-baseline-inhalation | -0.0750 | 0.0187 | -4.010 | 4.31e-4 | **0.0020** |
| HCT-baseline-exhalation vs. HCT-baseline-inhalation | 0.0090 | 0.0267 | 0.338 | 0.7377 | 0.7377 |

**Supplementary Table 12. Planned comparisons of theta-band graph clustering coefficient.** Abbreviations: HCT = heartbeat counting task, C-TCT = cardiac-tone counting task, SE = standard error, FDR = false discovery rate.

| **Comparison** | **Difference** | **SE** | **t** | **p** | **p_FDR_** |
| --- | --- | --- | --- | --- | --- |
| C-TCT-active-exhalation vs. C-TCT-active-inhalation | -0.0176 | 0.0447 | -0.3938 | 0.6967 | 0.6968 |
| C-TCT-active-exhalation vs. C-TCT-baseline-exhalation | -0.0471 | 0.0410 | -1.1507 | 0.2599 | 0.5199 |
| C-TCT-active-inhalation vs. C-TCT-baseline-inhalation | -0.0581 | 0.0414 | -1.4040 | 0.1716 | 0.5199 |
| C-TCT-baseline-exhalation vs. C-TCT-baseline-inhalation | -0.0286 | 0.0407 | -0.7032 | 0.4879 | 0.6505 |
| HCT-active-exhalation vs. HCT-active-inhalation | 0.0722 | 0.0367 | 1.9674 | 0.0594 | 0.1190 |
| HCT-active-exhalation vs. HCT-baseline-exhalation | -0.0393 | 0.0490 | -0.8042 | 0.4282 | 0.5710 |
| HCT-active-inhalation vs. HCT-baseline-inhalation | -0.1146 | 0.0287 | -3.9880 | 4.57e-4 | **0.0018** |
| HCT-baseline-exhalation vs. HCT-baseline-inhalation | -0.0030 | 0.0398 | -0.0752 | 0.9405 | 0.9406 |

**Supplementary Table 13. Planned comparisons of theta-band graph local efficiency.** Abbreviations: HCT = heartbeat counting task, C-TCT = cardiac-tone counting task, SE = standard error, FDR = false discovery rate.

| **Comparison** | **Difference** | **SE** | **t** | **p** | **p_FDR_** |
| --- | --- | --- | --- | --- | --- |
| C-TCT-active-exhalation vs. C-TCT-active-inhalation | 0.0081 | 0.0302 | 0.2712 | 0.7882 | 0.7883 |
| C-TCT-active-exhalation vs. C-TCT-baseline-exhalation | -0.0243 | 0.0311 | -0.7826 | 0.4406 | 0.5875 |
| C-TCT-active-inhalation vs. C-TCT-baseline-inhalation | -0.0662 | 0.0308 | -2.1547 | 0.0402 | 0.1611 |
| C-TCT-baseline-exhalation vs. C-TCT-baseline-inhalation | -0.0337 | 0.0279 | -1.2065 | 0.2380 | 0.4762 |
| HCT-active-exhalation vs. HCT-active-inhalation | 0.0423 | 0.0293 | 1.4438 | 0.1603 | 0.3206 |
| HCT-active-exhalation vs. HCT-baseline-exhalation | -0.0274 | 0.0369 | -0.7430 | 0.4638 | 0.6185 |
| HCT-active-inhalation vs. HCT-baseline-inhalation | -0.0726 | 0.0191 | -3.7934 | 7.63e-4 | **0.0031** |
| HCT-baseline-exhalation vs. HCT-baseline-inhalation | -0.0028 | 0.0257 | -0.1096 | 0.9135 | 0.9135 |

**Supplementary Table 14. Planned comparisons of theta-band graph global efficiency.** Abbreviations: HCT = heartbeat counting task, C-TCT = cardiac-tone counting task, SE = standard error, FDR = false discovery rate.

|  | **Estimate** | **SE** | **t** | **p** |
| --- | --- | --- | --- | --- |
| Intercept | 0.2342 | 0.0049 | 47.308 | 1.64e-27 |
| Attention | 0.0558 | 0.0262 | 2.129 | 0.0425 |
| Time | -0.0360 | 0.0076 | -4.720 | 6.46e-5 |
| Phase | -0.0184 | 0.0096 | -1.904 | 0.0676 |
| Threshold 2 - 1 | 0.0558 | 0.0020 | 27.945 | 1.10e-130 |
| Threshold 3 - 1 | 0.1017 | 0.0020 | 50.879 | 6.27e-293 |
| Threshold 4 - 1 | 0.1412 | 0.0020 | 70.640 | 0.0000 |
| Threshold 5 - 1 | 0.1763 | 0.0020 | 88.149 | 0.0000 |
| Threshold 6 - 1 | 0.2104 | 0.0020 | 105.232 | 0.0000 |
| Attention * Time | 0.0096 | 0.0148 | 0.648 | 0.5228 |
| Attention * Phase | -0.0335 | 0.0190 | -1.760 | 0.0898 |
| Time * Phase | 0.0127 | 0.0206 | 0.620 | 0.5408 |
| Attention * Time * Phase | 0.0412 | 0.0390 | 1.055 | 0.3009 |

**Supplementary Table 15. Parameter estimates of the linear mixed-effects model for alpha-band graph density.** Abbreviations: SE = standard error.

|  | **Estimate** | **SE** | **t** | **p** |
| --- | --- | --- | --- | --- |
| Intercept | 0.2781 | 0.0107 | 25.782 | 1.50e-20 |
| Attention | 0.0607 | 0.0295 | 2.059 | 0.0493 |
| Time | -0.0335 | 0.0090 | -3.732 | 8.95e-4 |
| Phase | -0.0122 | 0.0096 | -1.274 | 0.2137 |
| Threshold 2 - 1 | 0.0631 | 0.0030 | 20.477 | 2.33e-79 |
| Threshold 3 - 1 | 0.1125 | 0.0030 | 36.483 | 1.98e-192 |
| Threshold 4 - 1 | 0.1568 | 0.0030 | 50.815 | 1.67e-292 |
| Threshold 5 - 1 | 0.1946 | 0.0030 | 63.082 | 0.0000 |
| Threshold 6 - 1 | 0.2292 | 0.0030 | 74.281 | 0.0000 |
| Attention * Time | 0.0118 | 0.0172 | 0.688 | 0.4972 |
| Attention * Phase | -0.0209 | 0.0202 | -1.036 | 0.3095 |
| Time * Phase | 0.0090 | 0.0206 | 0.440 | 0.6636 |
| Attention * Time * Phase | 0.0404 | 0.0468 | 0.864 | 0.3949 |

**Supplementary Table 16. Parameter estimates of the linear mixed-effects model for alpha-band graph clustering coefficient.** Abbreviations: SE = standard error.

|  | **Estimate** | **SE** | **t** | **p** |
| --- | --- | --- | --- | --- |
| Intercept | 0.4566 | 0.0165 | 27.632 | 2.46e-21 |
| Attention | 0.0969 | 0.0444 | 2.178 | 0.0383 |
| Time | -0.0520 | 0.0126 | -4.115 | 3.26e-4 |
| Phase | -0.0168 | 0.0145 | -1.155 | 0.2581 |
| Threshold 2 - 1 | 0.1126 | 0.0055 | 20.213 | 1.18e-77 |
| Threshold 3 - 1 | 0.1960 | 0.0055 | 35.198 | 3.99e-183 |
| Threshold 4 - 1 | 0.2640 | 0.0055 | 47.397 | 1.91e-269 |
| Threshold 5 - 1 | 0.3179 | 0.0055 | 57.080 | 0.0000 |
| Threshold 6 - 1 | 0.3630 | 0.0055 | 65.180 | 0.0000 |
| Attention * Time | 0.0319 | 0.0232 | 1.372 | 0.1813 |
| Attention * Phase | -0.0284 | 0.0284 | -1.000 | 0.3260 |
| Time * Phase | 0.0194 | 0.0290 | 0.667 | 0.5106 |
| Attention * Time * Phase | 0.0649 | 0.0632 | 1.027 | 0.3137 |

**Supplementary Table 17. Parameter estimates of the linear mixed-effects model for alpha-band graph local efficiency.** Abbreviations: SE = standard error.

|  | **Estimate** | **SE** | **t** | **p** |
| --- | --- | --- | --- | --- |
| Intercept | 0.5182 | 0.0126 | 41.075 | 7.07e-26 |
| Attention | 0.0763 | 0.0324 | 2.353 | 0.0262 |
| Time | -0.0408 | 0.0094 | -4.343 | 1.78e-4 |
| Phase | -0.0165 | 0.0106 | -1.550 | 0.1328 |
| Threshold 2 - 1 | 0.1033 | 0.0040 | 25.763 | 3.25e-115 |
| Threshold 3 - 1 | 0.1708 | 0.0040 | 42.603 | 3.62e-236 |
| Threshold 4 - 1 | 0.2166 | 0.0040 | 54.014 | 1.58e-313 |
| Threshold 5 - 1 | 0.2537 | 0.0040 | 63.269 | 0.0000 |
| Threshold 6 - 1 | 0.2859 | 0.0040 | 71.300 | 0.0000 |
| Attention * Time | 0.0309 | 0.0165 | 1.861 | 0.0737 |
| Attention * Phase | -0.0321 | 0.0204 | -1.569 | 0.1283 |
| Time * Phase | 0.0150 | 0.0210 | 0.714 | 0.4816 |
| Attention * Time * Phase | 0.0519 | 0.0406 | 1.276 | 0.2127 |

**Supplementary Table 18. Parameter estimates of the linear mixed-effects model for alpha-band graph global efficiency.** Abbreviations: SE = standard error.

| **Comparison** | **Difference** | **SE** | **t** | **p** | **p_FDR_** |
| --- | --- | --- | --- | --- | --- |
| C-TCT-active-exhalation vs. C-TCT-active-inhalation | -0.0022 | 0.0169 | -0.132 | 0.8962 | 0.8962 |
| C-TCT-active-exhalation vs. C-TCT-baseline-exhalation | 0.0369 | 0.0195 | 1.896 | 0.0686 | 0.1373 |
| C-TCT-active-inhalation vs. C-TCT-baseline-inhalation | 0.0447 | 0.0140 | 3.188 | 0.0036 | **0.0144** |
| C-TCT-baseline-exhalation vs. C-TCT-baseline-inhalation | 0.0056 | 0.0207 | 0.271 | 0.7885 | 0.8963 |
| HCT-active-exhalation vs. HCT-active-inhalation | 0.0518 | 0.0208 | 2.497 | 0.0189 | **0.0378** |
| HCT-active-exhalation vs. HCT-baseline-exhalation | 0.0479 | 0.0170 | 2.820 | 0.0089 | **0.0356** |
| HCT-active-inhalation vs. HCT-baseline-inhalation | 0.0145 | 0.0199 | 0.730 | 0.4718 | 0.4718 |
| HCT-baseline-exhalation vs. HCT-baseline-inhalation | 0.0184 | 0.0200 | 0.924 | 0.3635 | 0.4718 |

**Supplementary Table 19. Planned comparisons of alpha-band graph density.** Abbreviations: HCT = heartbeat counting task, C-TCT = cardiac-tone counting task, SE = standard error, FDR = false discovery rate.

| **Comparison** | **Difference** | **SE** | **t** | **p** | **p_FDR_** |
| --- | --- | --- | --- | --- | --- |
| C-TCT-active-exhalation vs. C-TCT-active-inhalation | -0.0038 | 0.0178 | -0.214 | 0.8325 | 0.8325 |
| C-TCT-active-exhalation vs. C-TCT-baseline-exhalation | 0.0339 | 0.0190 | 1.782 | 0.0860 | 0.1721 |
| C-TCT-active-inhalation vs. C-TCT-baseline-inhalation | 0.0451 | 0.0184 | 2.448 | 0.0211 | 0.0846 |
| C-TCT-baseline-exhalation vs. C-TCT-baseline-inhalation | 0.0073 | 0.0213 | 0.346 | 0.7320 | 0.8325 |
| HCT-active-exhalation vs. HCT-active-inhalation | 0.0374 | 0.0232 | 1.615 | 0.1179 | 0.2359 |
| HCT-active-exhalation vs. HCT-baseline-exhalation | 0.0423 | 0.0199 | 2.121 | 0.0432 | 0.1730 |
| HCT-active-inhalation vs. HCT-baseline-inhalation | 0.0129 | 0.0223 | 0.583 | 0.5645 | 0.7052 |
| HCT-baseline-exhalation vs. HCT-baseline-inhalation | 0.0080 | 0.0212 | 0.382 | 0.7052 | 0.7052 |

**Supplementary Table 20. Planned comparisons of alpha-band graph clustering coefficient.** Abbreviations: HCT = heartbeat counting task, C-TCT = cardiac-tone counting task, SE = standard error, FDR = false discovery rate.

| **Comparison** | **Difference** | **SE** | **t** | **p** | **p_FDR_** |
| --- | --- | --- | --- | --- | --- |
| C-TCT-active-exhalation vs. C-TCT-active-inhalation | -0.0040 | 0.0238 | -0.168 | 0.8674 | 0.8675 |
| C-TCT-active-exhalation vs. C-TCT-baseline-exhalation | 0.0614 | 0.0264 | 2.326 | 0.0277 | 0.0556 |
| C-TCT-active-inhalation vs. C-TCT-baseline-inhalation | 0.0745 | 0.0259 | 2.871 | 0.0078 | **0.0314** |
| C-TCT-baseline-exhalation vs. C-TCT-baseline-inhalation | 0.0090 | 0.0308 | 0.295 | 0.7702 | 0.8675 |
| HCT-active-exhalation vs. HCT-active-inhalation | 0.0568 | 0.0324 | 1.758 | 0.0900 | 0.1800 |
| HCT-active-exhalation vs. HCT-baseline-exhalation | 0.0619 | 0.0269 | 2.301 | 0.0293 | 0.1174 |
| HCT-active-inhalation vs. HCT-baseline-inhalation | 0.0101 | 0.0305 | 0.332 | 0.7423 | 0.8696 |
| HCT-baseline-exhalation vs. HCT-baseline-inhalation | 0.0050 | 0.0305 | 0.166 | 0.8696 | 0.8696 |

**Supplementary Table 21. Planned comparisons of alpha-band graph local efficiency.** Abbreviations: HCT = heartbeat counting task, C-TCT = cardiac-tone counting task, SE = standard error, FDR = false discovery rate.

| **Comparison** | **Difference** | **SE** | **t** | **p** | **p_FDR_** |
| --- | --- | --- | --- | --- | --- |
| C-TCT-active-exhalation vs. C-TCT-active-inhalation | -0.0049 | 0.0170 | -0.292 | 0.77225 | 0.8098 |
| C-TCT-active-exhalation vs. C-TCT-baseline-exhalation | 0.0508 | 0.0224 | 2.272 | 0.03127 | 0.0625 |
| C-TCT-active-inhalation vs. C-TCT-baseline-inhalation | 0.0617 | 0.0165 | 3.751 | 8.53e-4 | **0.0034** |
| C-TCT-baseline-exhalation vs. C-TCT-baseline-inhalation | 0.0059 | 0.0244 | 0.243 | 0.80978 | 0.8098 |
| HCT-active-exhalation vs. HCT-active-inhalation | 0.0530 | 0.0212 | 2.499 | 0.01884 | **0.0377** |
| HCT-active-exhalation vs. HCT-baseline-exhalation | 0.0458 | 0.0175 | 2.620 | 0.01424 | **0.0377** |
| HCT-active-inhalation vs. HCT-baseline-inhalation | 0.0049 | 0.0202 | 0.243 | 0.81013 | 0.8101 |
| HCT-baseline-exhalation vs. HCT-baseline-inhalation | 0.0120 | 0.0198 | 0.610 | 0.54702 | 0.7294 |

**Supplementary Table 22. Planned comparisons of alpha-band graph global efficiency.** Abbreviations: HCT = heartbeat counting task, C-TCT = cardiac-tone counting task, SE = standard error, FDR = false discovery rate.

|  | **HCT** | | | | | | | | | | | | | | | | | |
| --- | --- | --- | --- | --- | --- | --- | --- | --- | --- | --- | --- | --- | --- | --- | --- | --- | --- | --- |
|  | **Theta** | | | | | | | | | **Alpha** | | | | | | | | |
|  | **ΔHEP and Δpower** | | | **ΔHEP and ΔITC** | | | **Δpower and ΔITC** | | | **ΔHEP and Δpower** | | | **ΔHEP and ΔITC** | | | **Δpower and ΔITC** | | |
| **Channel** | **rho** | **p** | **p_FDR_** | **rho** | **p** | **p_FDR_** | **rho** | **p** | **p_FDR_** | **rho** | **p** | **p_FDR_** | **rho** | **p** | **p_FDR_** | **rho** | **p** | **p_FDR_** |
| **Fp1** | 0.114 | 0.563 | 0.963 | -0.149 | 0.448 | 0.752 | -0.269 | 0.166 | 0.963 | 0.367 | 0.055 | 0.179 | 0.025 | 0.899 | 0.920 | -0.089 | 0.651 | 0.962 |
| **Fp2** | -0.112 | 0.568 | 0.963 | -0.086 | 0.663 | 0.883 | 0.103 | 0.601 | 0.963 | 0.437 | 0.021 | 0.132 | -0.105 | 0.595 | 0.722 | -0.176 | 0.370 | 0.849 |
| **F3** | -0.005 | 0.981 | 0.981 | -0.223 | 0.253 | 0.625 | -0.218 | 0.264 | 0.963 | 0.317 | 0.101 | 0.246 | -0.233 | 0.231 | 0.509 | -0.260 | 0.181 | 0.667 |
| **F4** | 0.248 | 0.202 | 0.963 | -0.236 | 0.225 | 0.621 | 0.085 | 0.667 | 0.963 | 0.577 | 0.002 | **0.038** | -0.588 | 0.001 | 0.054 | -0.267 | 0.170 | 0.667 |
| **C3** | -0.138 | 0.482 | 0.963 | 0.020 | 0.919 | 0.952 | 0.017 | 0.932 | 0.997 | -0.160 | 0.415 | 0.598 | -0.018 | 0.930 | 0.930 | 0.273 | 0.159 | 0.667 |
| **C4** | 0.258 | 0.184 | 0.963 | -0.413 | 0.030 | 0.325 | -0.240 | 0.217 | 0.963 | 0.005 | 0.981 | 0.999 | -0.132 | 0.502 | 0.722 | -0.243 | 0.212 | 0.717 |
| **P3** | -0.014 | 0.943 | 0.965 | 0.138 | 0.481 | 0.755 | 0.057 | 0.773 | 0.963 | -0.258 | 0.184 | 0.352 | 0.551 | 0.003 | 0.055 | 0.049 | 0.805 | 0.962 |
| **P4** | -0.163 | 0.407 | 0.963 | -0.213 | 0.275 | 0.638 | -0.044 | 0.823 | 0.963 | -0.355 | 0.064 | 0.189 | 0.108 | 0.584 | 0.722 | -0.259 | 0.182 | 0.667 |
| **Fz** | 0.264 | 0.173 | 0.963 | -0.337 | 0.080 | 0.438 | 0.042 | 0.831 | 0.963 | 0.573 | 0.002 | **0.038** | -0.182 | 0.352 | 0.590 | -0.003 | 0.988 | 0.990 |
| **Cz** | 0.267 | 0.169 | 0.963 | -0.108 | 0.582 | 0.853 | -0.018 | 0.930 | 0.997 | 0.292 | 0.131 | 0.275 | -0.111 | 0.574 | 0.722 | -0.156 | 0.426 | 0.849 |
| **Pz** | -0.094 | 0.633 | 0.963 | -0.054 | 0.786 | 0.883 | 0.083 | 0.673 | 0.963 | -0.186 | 0.341 | 0.537 | -0.187 | 0.339 | 0.590 | 0.141 | 0.472 | 0.849 |
| **AFz** | 0.061 | 0.758 | 0.965 | -0.398 | 0.037 | 0.325 | 0.047 | 0.812 | 0.963 | 0.519 | 0.005 | **0.044** | -0.156 | 0.426 | 0.670 | -0.188 | 0.336 | 0.849 |
| **FC1** | 0.101 | 0.607 | 0.963 | -0.129 | 0.511 | 0.775 | -0.134 | 0.495 | 0.963 | 0.264 | 0.173 | 0.347 | -0.383 | 0.045 | 0.219 | -0.158 | 0.421 | 0.849 |
| **FC2** | 0.172 | 0.379 | 0.963 | -0.505 | 0.007 | 0.299 | -0.042 | 0.831 | 0.963 | 0.300 | 0.120 | 0.275 | -0.479 | 0.011 | 0.112 | -0.278 | 0.152 | 0.667 |
| **CP1** | 0.111 | 0.572 | 0.963 | -0.296 | 0.127 | 0.465 | 0.061 | 0.758 | 0.963 | 0.138 | 0.482 | 0.631 | 0.146 | 0.458 | 0.695 | 0.003 | 0.990 | 0.990 |
| **CP2** | 0.100 | 0.611 | 0.963 | -0.326 | 0.091 | 0.438 | -0.194 | 0.320 | 0.963 | 0.348 | 0.071 | 0.194 | -0.215 | 0.272 | 0.543 | -0.286 | 0.140 | 0.667 |
| **FC5** | -0.080 | 0.685 | 0.963 | -0.157 | 0.425 | 0.752 | -0.097 | 0.623 | 0.963 | 0.080 | 0.683 | 0.771 | -0.222 | 0.256 | 0.536 | 0.038 | 0.849 | 0.962 |
| **FC6** | -0.044 | 0.825 | 0.965 | -0.002 | 0.992 | 0.992 | 0.170 | 0.386 | 0.963 | 0.032 | 0.871 | 0.912 | -0.469 | 0.013 | 0.112 | 0.034 | 0.864 | 0.962 |
| **CP5** | -0.184 | 0.347 | 0.963 | 0.018 | 0.930 | 0.952 | -0.177 | 0.365 | 0.963 | -0.116 | 0.555 | 0.660 | -0.030 | 0.882 | 0.920 | -0.042 | 0.833 | 0.962 |
| **CP6** | -0.072 | 0.714 | 0.963 | 0.084 | 0.669 | 0.883 | -0.384 | 0.044 | 0.963 | -0.044 | 0.823 | 0.883 | -0.055 | 0.782 | 0.860 | -0.464 | 0.014 | 0.286 |
| **F1** | 0.015 | 0.941 | 0.965 | -0.417 | 0.028 | 0.325 | -0.148 | 0.450 | 0.963 | 0.410 | 0.031 | 0.171 | -0.412 | 0.030 | 0.191 | -0.219 | 0.261 | 0.819 |
| **F2** | 0.081 | 0.681 | 0.963 | -0.377 | 0.049 | 0.360 | 0.056 | 0.775 | 0.963 | 0.366 | 0.056 | 0.179 | -0.449 | 0.017 | 0.127 | -0.055 | 0.780 | 0.962 |
| **C1** | 0.248 | 0.202 | 0.963 | -0.332 | 0.085 | 0.438 | 0.006 | 0.977 | 0.997 | 0.211 | 0.280 | 0.457 | -0.253 | 0.193 | 0.475 | -0.066 | 0.737 | 0.962 |
| **C2** | 0.127 | 0.518 | 0.963 | -0.194 | 0.320 | 0.671 | -0.058 | 0.769 | 0.963 | 0.132 | 0.502 | 0.631 | -0.178 | 0.362 | 0.590 | 0.147 | 0.453 | 0.849 |
| **P1** | 0.021 | 0.917 | 0.965 | -0.151 | 0.441 | 0.752 | 0.059 | 0.767 | 0.963 | -0.089 | 0.651 | 0.753 | 0.187 | 0.340 | 0.590 | -0.025 | 0.899 | 0.965 |
| **P2** | -0.218 | 0.263 | 0.963 | -0.179 | 0.361 | 0.690 | -0.091 | 0.644 | 0.963 | -0.340 | 0.077 | 0.200 | 0.094 | 0.634 | 0.735 | -0.031 | 0.875 | 0.962 |
| **AF3** | 0.160 | 0.413 | 0.963 | -0.438 | 0.021 | 0.325 | -0.010 | 0.961 | 0.997 | 0.521 | 0.005 | **0.044** | -0.386 | 0.043 | 0.219 | -0.442 | 0.020 | 0.286 |
| **AF4** | 0.102 | 0.605 | 0.963 | -0.236 | 0.226 | 0.621 | 0.292 | 0.132 | 0.963 | 0.522 | 0.005 | **0.044** | -0.284 | 0.144 | 0.421 | -0.157 | 0.425 | 0.849 |
| **FC3** | -0.025 | 0.899 | 0.965 | 0.049 | 0.803 | 0.883 | 0.096 | 0.625 | 0.963 | 0.236 | 0.226 | 0.413 | -0.375 | 0.050 | 0.219 | -0.055 | 0.780 | 0.962 |
| **FC4** | 0.269 | 0.165 | 0.963 | -0.206 | 0.292 | 0.643 | 0.025 | 0.901 | 0.997 | 0.158 | 0.421 | 0.598 | -0.300 | 0.120 | 0.378 | -0.461 | 0.014 | 0.286 |
| **CP3** | -0.053 | 0.788 | 0.965 | -0.309 | 0.109 | 0.438 | 0.259 | 0.182 | 0.963 | -0.067 | 0.733 | 0.806 | 0.252 | 0.194 | 0.475 | 0.183 | 0.349 | 0.849 |
| **CP4** | -0.070 | 0.722 | 0.963 | 0.073 | 0.712 | 0.883 | -0.437 | 0.021 | 0.925 | 0.001 | 0.999 | 0.999 | -0.122 | 0.536 | 0.722 | -0.421 | 0.026 | 0.291 |
| **PO3** | 0.161 | 0.412 | 0.963 | 0.042 | 0.833 | 0.894 | -0.108 | 0.582 | 0.963 | -0.371 | 0.053 | 0.179 | 0.181 | 0.355 | 0.590 | 0.299 | 0.122 | 0.667 |
| **PO4** | 0.030 | 0.882 | 0.965 | 0.067 | 0.735 | 0.883 | -0.099 | 0.615 | 0.963 | -0.365 | 0.057 | 0.179 | 0.088 | 0.655 | 0.738 | -0.181 | 0.356 | 0.849 |
| **F5** | -0.037 | 0.853 | 0.965 | 0.051 | 0.795 | 0.883 | -0.222 | 0.256 | 0.963 | 0.153 | 0.436 | 0.600 | 0.101 | 0.607 | 0.722 | -0.068 | 0.729 | 0.962 |
| **F6** | 0.227 | 0.245 | 0.963 | -0.309 | 0.109 | 0.438 | -0.076 | 0.702 | 0.963 | 0.367 | 0.055 | 0.179 | -0.536 | 0.004 | 0.055 | -0.138 | 0.482 | 0.849 |
| **C5** | -0.296 | 0.127 | 0.963 | -0.222 | 0.256 | 0.625 | -0.137 | 0.486 | 0.963 | -0.164 | 0.402 | 0.598 | -0.103 | 0.601 | 0.722 | -0.339 | 0.078 | 0.667 |
| **C6** | -0.088 | 0.657 | 0.963 | -0.188 | 0.336 | 0.672 | -0.142 | 0.470 | 0.963 | 0.227 | 0.244 | 0.413 | -0.244 | 0.211 | 0.488 | -0.057 | 0.771 | 0.962 |
| **P5** | 0.079 | 0.689 | 0.963 | 0.144 | 0.462 | 0.752 | 0.151 | 0.441 | 0.963 | -0.229 | 0.241 | 0.413 | 0.300 | 0.120 | 0.378 | 0.038 | 0.849 | 0.962 |
| **P6** | -0.190 | 0.332 | 0.963 | 0.060 | 0.760 | 0.883 | -0.216 | 0.269 | 0.963 | -0.512 | 0.006 | **0.044** | 0.318 | 0.099 | 0.364 | -0.185 | 0.344 | 0.849 |
| **Fpz** | 0.038 | 0.849 | 0.965 | -0.238 | 0.222 | 0.621 | 0.221 | 0.258 | 0.963 | 0.377 | 0.049 | 0.179 | -0.259 | 0.183 | 0.475 | -0.049 | 0.803 | 0.962 |
| **CPz** | 0.276 | 0.154 | 0.963 | -0.279 | 0.151 | 0.510 | 0.001 | 0.997 | 0.997 | 0.128 | 0.516 | 0.631 | 0.047 | 0.814 | 0.874 | -0.094 | 0.633 | 0.962 |
| **POz** | 0.078 | 0.691 | 0.963 | 0.057 | 0.771 | 0.883 | 0.096 | 0.625 | 0.963 | -0.128 | 0.514 | 0.631 | 0.118 | 0.549 | 0.722 | -0.003 | 0.990 | 0.990 |
| **FCz** | 0.248 | 0.203 | 0.963 | -0.084 | 0.669 | 0.883 | -0.050 | 0.799 | 0.963 | 0.294 | 0.128 | 0.275 | -0.330 | 0.087 | 0.347 | -0.124 | 0.529 | 0.895 |
|  | **HCT** | | | | | | | | |  | | | | | | | | |
|  | **Beta** | | | | | | | | | **Gamma** | | | | | | | | |
|  | **ΔHEP and Δpower** | | | **ΔHEP and ΔITC** | | | **Δpower and ΔITC** | | | **ΔHEP and Δpower** | | | **ΔHEP and ΔITC** | | | **Δpower and ΔITC** | | |
| **Channel** | **rho** | **p** | **p_FDR_** | **rho** | **p** | **p_FDR_** | **rho** | **p** | **p_FDR_** | **rho** | **p** | **p_FDR_** | **rho** | **p** | **p_FDR_** | **rho** | **p** | **p_FDR_** |
| **Fp1** | 0.011 | 0.954 | 0.954 | 0.388 | 0.042 | 0.617 | 0.143 | 0.465 | 0.775 | -0.134 | 0.495 | 0.732 | 0.094 | 0.633 | 0.947 | -0.104 | 0.597 | 0.847 |
| **Fp2** | 0.184 | 0.347 | 0.689 | 0.074 | 0.706 | 0.841 | 0.031 | 0.875 | 0.930 | 0.097 | 0.623 | 0.732 | -0.093 | 0.636 | 0.947 | -0.367 | 0.055 | 0.434 |
| **F3** | 0.250 | 0.199 | 0.554 | 0.282 | 0.146 | 0.617 | 0.178 | 0.364 | 0.775 | 0.169 | 0.388 | 0.732 | 0.337 | 0.080 | 0.707 | 0.146 | 0.458 | 0.793 |
| **F4** | 0.266 | 0.171 | 0.554 | -0.193 | 0.323 | 0.646 | -0.082 | 0.677 | 0.860 | -0.120 | 0.540 | 0.732 | -0.005 | 0.981 | 0.981 | 0.193 | 0.324 | 0.680 |
| **C3** | -0.195 | 0.318 | 0.665 | -0.065 | 0.743 | 0.841 | -0.116 | 0.555 | 0.814 | -0.183 | 0.349 | 0.732 | -0.189 | 0.334 | 0.893 | -0.195 | 0.319 | 0.680 |
| **C4** | -0.016 | 0.935 | 0.954 | 0.144 | 0.463 | 0.703 | 0.076 | 0.700 | 0.860 | -0.237 | 0.224 | 0.703 | -0.165 | 0.399 | 0.893 | -0.280 | 0.148 | 0.666 |
| **P3** | -0.323 | 0.093 | 0.533 | 0.066 | 0.737 | 0.841 | 0.288 | 0.136 | 0.690 | -0.129 | 0.513 | 0.732 | -0.027 | 0.890 | 0.947 | 0.254 | 0.192 | 0.666 |
| **P4** | 0.196 | 0.315 | 0.665 | -0.008 | 0.968 | 0.990 | 0.351 | 0.067 | 0.690 | 0.045 | 0.820 | 0.880 | -0.162 | 0.408 | 0.893 | 0.074 | 0.708 | 0.873 |
| **Fz** | 0.269 | 0.165 | 0.554 | 0.281 | 0.148 | 0.617 | -0.249 | 0.201 | 0.706 | 0.147 | 0.453 | 0.732 | 0.256 | 0.189 | 0.893 | 0.206 | 0.292 | 0.680 |
| **Cz** | -0.048 | 0.807 | 0.888 | -0.207 | 0.288 | 0.617 | 0.178 | 0.364 | 0.775 | -0.121 | 0.538 | 0.732 | 0.166 | 0.397 | 0.893 | -0.041 | 0.838 | 0.899 |
| **Pz** | -0.281 | 0.148 | 0.554 | -0.144 | 0.463 | 0.703 | 0.270 | 0.165 | 0.690 | -0.077 | 0.695 | 0.785 | -0.095 | 0.629 | 0.947 | -0.072 | 0.714 | 0.873 |
| **AFz** | 0.248 | 0.202 | 0.554 | 0.165 | 0.400 | 0.653 | -0.199 | 0.309 | 0.775 | -0.130 | 0.507 | 0.732 | 0.024 | 0.904 | 0.947 | 0.290 | 0.134 | 0.666 |
| **FC1** | 0.088 | 0.655 | 0.888 | 0.206 | 0.292 | 0.617 | 0.048 | 0.807 | 0.930 | 0.217 | 0.265 | 0.732 | 0.131 | 0.504 | 0.930 | 0.019 | 0.923 | 0.952 |
| **FC2** | 0.438 | 0.021 | 0.303 | 0.246 | 0.207 | 0.617 | -0.075 | 0.704 | 0.860 | 0.242 | 0.214 | 0.703 | 0.144 | 0.463 | 0.926 | 0.055 | 0.780 | 0.899 |
| **CP1** | -0.096 | 0.625 | 0.888 | 0.175 | 0.371 | 0.653 | 0.298 | 0.123 | 0.690 | -0.107 | 0.587 | 0.732 | 0.049 | 0.805 | 0.947 | 0.194 | 0.320 | 0.680 |
| **CP2** | 0.118 | 0.549 | 0.863 | -0.239 | 0.220 | 0.617 | 0.174 | 0.376 | 0.775 | 0.238 | 0.223 | 0.703 | -0.303 | 0.118 | 0.862 | 0.092 | 0.640 | 0.847 |
| **FC5** | 0.168 | 0.391 | 0.689 | -0.205 | 0.295 | 0.617 | -0.158 | 0.421 | 0.775 | 0.377 | 0.049 | 0.536 | 0.156 | 0.426 | 0.893 | -0.005 | 0.979 | 0.979 |
| **FC6** | 0.157 | 0.423 | 0.689 | -0.285 | 0.142 | 0.617 | -0.018 | 0.930 | 0.930 | 0.102 | 0.603 | 0.732 | 0.032 | 0.871 | 0.947 | -0.381 | 0.046 | 0.434 |
| **CP5** | 0.070 | 0.722 | 0.888 | -0.365 | 0.057 | 0.617 | -0.057 | 0.773 | 0.919 | 0.149 | 0.446 | 0.732 | -0.032 | 0.873 | 0.947 | 0.409 | 0.031 | 0.434 |
| **CP6** | 0.178 | 0.364 | 0.689 | -0.207 | 0.288 | 0.617 | -0.314 | 0.104 | 0.690 | -0.058 | 0.769 | 0.846 | -0.118 | 0.549 | 0.930 | 0.046 | 0.816 | 0.899 |
| **F1** | 0.157 | 0.423 | 0.689 | 0.109 | 0.580 | 0.808 | -0.209 | 0.286 | 0.775 | 0.288 | 0.137 | 0.658 | 0.425 | 0.025 | 0.366 | 0.431 | 0.023 | 0.434 |
| **F2** | 0.158 | 0.421 | 0.689 | 0.216 | 0.269 | 0.617 | -0.158 | 0.421 | 0.775 | -0.033 | 0.868 | 0.910 | 0.437 | 0.021 | 0.366 | -0.124 | 0.529 | 0.817 |
| **C1** | -0.063 | 0.750 | 0.888 | -0.059 | 0.765 | 0.841 | 0.339 | 0.078 | 0.690 | 0.135 | 0.491 | 0.732 | -0.071 | 0.720 | 0.947 | -0.018 | 0.930 | 0.952 |
| **C2** | 0.016 | 0.935 | 0.954 | -0.054 | 0.784 | 0.841 | 0.034 | 0.864 | 0.930 | -0.138 | 0.482 | 0.732 | 0.127 | 0.518 | 0.930 | 0.049 | 0.805 | 0.899 |
| **P1** | -0.333 | 0.084 | 0.533 | 0.010 | 0.961 | 0.990 | 0.311 | 0.108 | 0.690 | -0.403 | 0.034 | 0.536 | -0.228 | 0.242 | 0.893 | 0.115 | 0.557 | 0.817 |
| **P2** | -0.092 | 0.640 | 0.888 | -0.073 | 0.712 | 0.841 | 0.265 | 0.173 | 0.690 | -0.015 | 0.941 | 0.941 | -0.068 | 0.729 | 0.947 | 0.088 | 0.655 | 0.847 |
| **AF3** | 0.072 | 0.716 | 0.888 | 0.070 | 0.722 | 0.841 | -0.135 | 0.493 | 0.775 | -0.094 | 0.633 | 0.732 | 0.045 | 0.820 | 0.947 | 0.041 | 0.838 | 0.899 |
| **AF4** | 0.320 | 0.097 | 0.533 | 0.177 | 0.365 | 0.653 | -0.134 | 0.496 | 0.775 | -0.339 | 0.078 | 0.571 | 0.016 | 0.935 | 0.956 | -0.376 | 0.049 | 0.434 |
| **FC3** | 0.068 | 0.731 | 0.888 | -0.217 | 0.265 | 0.617 | -0.194 | 0.322 | 0.775 | 0.170 | 0.385 | 0.732 | 0.056 | 0.777 | 0.947 | 0.097 | 0.623 | 0.847 |
| **FC4** | 0.053 | 0.790 | 0.888 | 0.213 | 0.274 | 0.617 | 0.414 | 0.029 | 0.690 | 0.158 | 0.421 | 0.732 | 0.220 | 0.259 | 0.893 | -0.115 | 0.557 | 0.817 |
| **CP3** | -0.059 | 0.767 | 0.888 | -0.246 | 0.206 | 0.617 | -0.245 | 0.209 | 0.706 | -0.360 | 0.061 | 0.536 | -0.085 | 0.665 | 0.947 | 0.136 | 0.489 | 0.797 |
| **CP4** | 0.241 | 0.215 | 0.557 | -0.167 | 0.393 | 0.653 | -0.101 | 0.607 | 0.834 | -0.025 | 0.899 | 0.920 | -0.206 | 0.292 | 0.893 | -0.142 | 0.468 | 0.793 |
| **PO3** | -0.522 | 0.005 | 0.216 | 0.269 | 0.166 | 0.617 | -0.130 | 0.509 | 0.775 | -0.365 | 0.057 | 0.536 | -0.229 | 0.241 | 0.893 | 0.211 | 0.279 | 0.680 |
| **PO4** | -0.067 | 0.735 | 0.888 | -0.132 | 0.502 | 0.736 | 0.084 | 0.669 | 0.860 | -0.278 | 0.152 | 0.658 | -0.246 | 0.206 | 0.893 | 0.174 | 0.376 | 0.751 |
| **F5** | 0.207 | 0.288 | 0.665 | 0.107 | 0.587 | 0.808 | -0.301 | 0.120 | 0.690 | 0.295 | 0.127 | 0.658 | -0.216 | 0.268 | 0.893 | -0.275 | 0.157 | 0.666 |
| **F6** | 0.259 | 0.182 | 0.554 | -0.283 | 0.144 | 0.617 | 0.142 | 0.468 | 0.775 | -0.149 | 0.446 | 0.732 | 0.122 | 0.535 | 0.930 | 0.201 | 0.304 | 0.680 |
| **C5** | -0.103 | 0.601 | 0.888 | -0.063 | 0.748 | 0.841 | 0.129 | 0.511 | 0.775 | 0.270 | 0.165 | 0.658 | 0.182 | 0.352 | 0.893 | 0.160 | 0.413 | 0.791 |
| **C6** | 0.276 | 0.154 | 0.554 | -0.233 | 0.231 | 0.617 | -0.025 | 0.901 | 0.930 | 0.152 | 0.440 | 0.732 | 0.059 | 0.765 | 0.947 | -0.298 | 0.123 | 0.666 |
| **P5** | -0.359 | 0.061 | 0.533 | 0.054 | 0.784 | 0.841 | 0.110 | 0.576 | 0.817 | 0.126 | 0.522 | 0.732 | -0.165 | 0.399 | 0.893 | 0.362 | 0.059 | 0.434 |
| **P6** | -0.032 | 0.873 | 0.937 | -0.263 | 0.175 | 0.617 | 0.269 | 0.165 | 0.690 | -0.138 | 0.482 | 0.732 | -0.443 | 0.019 | 0.366 | 0.209 | 0.286 | 0.680 |
| **Fpz** | 0.348 | 0.071 | 0.533 | 0.224 | 0.250 | 0.617 | -0.132 | 0.500 | 0.775 | -0.097 | 0.623 | 0.732 | 0.068 | 0.729 | 0.947 | 0.146 | 0.456 | 0.793 |
| **CPz** | -0.218 | 0.263 | 0.643 | -0.001 | 0.999 | 0.999 | 0.129 | 0.511 | 0.775 | -0.103 | 0.601 | 0.732 | 0.198 | 0.312 | 0.893 | 0.257 | 0.187 | 0.666 |
| **POz** | -0.251 | 0.197 | 0.554 | 0.172 | 0.379 | 0.653 | 0.021 | 0.915 | 0.930 | -0.426 | 0.025 | 0.536 | -0.044 | 0.825 | 0.947 | -0.243 | 0.212 | 0.666 |
| **FCz** | 0.463 | 0.014 | 0.303 | 0.223 | 0.253 | 0.617 | -0.034 | 0.862 | 0.930 | 0.294 | 0.128 | 0.658 | 0.365 | 0.057 | 0.630 | 0.245 | 0.208 | 0.666 |
|  | **C-TCT** | | | | | | | | | | | | | | | | | |
|  | **Theta** | | | | | | | | | **Alpha** | | | | | | | | |
|  | **ΔHEP and Δpower** | | | **ΔHEP and ΔITC** | | | **Δpower and ΔITC** | | | **ΔHEP and Δpower** | | | **ΔHEP and ΔITC** | | | **Δpower and ΔITC** | | |
| **Channel** | **rho** | **p** | **p_FDR_** | **rho** | **p** | **p_FDR_** | **rho** | **p** | **p_FDR_** | **rho** | **p** | **p_FDR_** | **rho** | **p** | **p_FDR_** | **rho** | **p** | **p_FDR_** |
| **Fp1** | -0.248 | 0.202 | 0.868 | 0.118 | 0.548 | 0.983 | -0.137 | 0.486 | 0.913 | -0.359 | 0.062 | 0.679 | -0.063 | 0.750 | 0.947 | -0.190 | 0.332 | 0.949 |
| **Fp2** | -0.143 | 0.465 | 0.868 | -0.197 | 0.313 | 0.983 | -0.049 | 0.805 | 0.973 | -0.306 | 0.113 | 0.887 | -0.142 | 0.470 | 0.947 | 0.016 | 0.937 | 1.000 |
| **F3** | -0.030 | 0.879 | 0.967 | -0.074 | 0.708 | 0.983 | -0.019 | 0.923 | 0.979 | 0.074 | 0.706 | 0.997 | 0.056 | 0.777 | 0.947 | -0.051 | 0.795 | 1.000 |
| **F4** | 0.161 | 0.412 | 0.868 | -0.372 | 0.052 | 0.983 | 0.271 | 0.162 | 0.728 | -0.028 | 0.886 | 0.997 | -0.087 | 0.659 | 0.947 | 0.053 | 0.788 | 1.000 |
| **C3** | -0.024 | 0.906 | 0.972 | 0.008 | 0.968 | 0.983 | 0.315 | 0.103 | 0.728 | -0.109 | 0.578 | 0.997 | 0.328 | 0.089 | 0.652 | 0.230 | 0.238 | 0.949 |
| **C4** | -0.147 | 0.453 | 0.868 | 0.080 | 0.685 | 0.983 | -0.120 | 0.540 | 0.914 | 0.189 | 0.333 | 0.997 | 0.001 | 0.999 | 0.999 | 0.069 | 0.727 | 1.000 |
| **P3** | -0.052 | 0.792 | 0.967 | -0.261 | 0.179 | 0.983 | 0.410 | 0.031 | 0.457 | 0.042 | 0.831 | 0.997 | 0.388 | 0.042 | 0.547 | -0.233 | 0.231 | 0.949 |
| **P4** | -0.057 | 0.771 | 0.967 | -0.057 | 0.773 | 0.983 | 0.075 | 0.704 | 0.968 | 0.036 | 0.855 | 0.997 | -0.048 | 0.807 | 0.947 | -0.157 | 0.425 | 0.949 |
| **Fz** | 0.050 | 0.799 | 0.967 | -0.235 | 0.227 | 0.983 | 0.154 | 0.431 | 0.913 | 0.225 | 0.249 | 0.997 | -0.014 | 0.943 | 0.988 | -0.001 | 0.999 | 1.000 |
| **Cz** | -0.286 | 0.140 | 0.868 | 0.129 | 0.511 | 0.983 | 0.216 | 0.268 | 0.842 | -0.050 | 0.799 | 0.997 | 0.275 | 0.157 | 0.928 | 0.037 | 0.853 | 1.000 |
| **Pz** | -0.124 | 0.529 | 0.895 | -0.004 | 0.983 | 0.983 | 0.158 | 0.420 | 0.913 | -0.108 | 0.582 | 0.997 | 0.196 | 0.316 | 0.947 | -0.133 | 0.498 | 0.983 |
| **AFz** | 0.257 | 0.186 | 0.868 | -0.214 | 0.273 | 0.983 | -0.021 | 0.917 | 0.979 | 0.131 | 0.505 | 0.997 | -0.073 | 0.712 | 0.947 | -0.158 | 0.420 | 0.949 |
| **FC1** | -0.459 | 0.015 | 0.657 | 0.024 | 0.906 | 0.983 | 0.296 | 0.126 | 0.728 | 0.389 | 0.042 | 0.679 | -0.019 | 0.923 | 0.988 | -0.331 | 0.086 | 0.949 |
| **FC2** | -0.191 | 0.329 | 0.868 | 0.094 | 0.633 | 0.983 | 0.269 | 0.165 | 0.728 | -0.082 | 0.679 | 0.997 | 0.103 | 0.599 | 0.947 | -0.117 | 0.551 | 0.983 |
| **CP1** | -0.068 | 0.729 | 0.967 | 0.039 | 0.844 | 0.983 | 0.186 | 0.343 | 0.888 | 0.022 | 0.910 | 0.997 | 0.358 | 0.062 | 0.547 | -0.154 | 0.431 | 0.949 |
| **CP2** | -0.366 | 0.056 | 0.822 | -0.009 | 0.966 | 0.983 | -0.030 | 0.879 | 0.979 | -0.072 | 0.716 | 0.997 | 0.159 | 0.418 | 0.947 | -0.259 | 0.183 | 0.949 |
| **FC5** | 0.145 | 0.460 | 0.868 | 0.117 | 0.551 | 0.983 | 0.097 | 0.621 | 0.948 | 0.014 | 0.943 | 0.997 | -0.059 | 0.767 | 0.947 | 0.302 | 0.118 | 0.949 |
| **FC6** | -0.034 | 0.864 | 0.967 | -0.252 | 0.194 | 0.983 | -0.004 | 0.986 | 0.994 | -0.001 | 0.997 | 0.997 | 0.090 | 0.647 | 0.947 | 0.198 | 0.311 | 0.949 |
| **CP5** | -0.100 | 0.611 | 0.960 | -0.219 | 0.261 | 0.983 | 0.207 | 0.288 | 0.845 | 0.073 | 0.712 | 0.997 | 0.371 | 0.053 | 0.547 | -0.010 | 0.961 | 1.000 |
| **CP6** | -0.048 | 0.810 | 0.967 | -0.040 | 0.840 | 0.983 | -0.198 | 0.312 | 0.858 | -0.300 | 0.121 | 0.887 | -0.051 | 0.795 | 0.947 | -0.144 | 0.463 | 0.971 |
| **F1** | 0.128 | 0.516 | 0.895 | -0.250 | 0.199 | 0.983 | 0.124 | 0.527 | 0.914 | 0.372 | 0.052 | 0.679 | -0.213 | 0.274 | 0.928 | -0.190 | 0.330 | 0.949 |
| **F2** | 0.178 | 0.362 | 0.868 | -0.413 | 0.030 | 0.983 | 0.114 | 0.563 | 0.917 | 0.004 | 0.986 | 0.997 | 0.045 | 0.818 | 0.947 | -0.108 | 0.582 | 0.983 |
| **C1** | -0.195 | 0.318 | 0.868 | 0.294 | 0.129 | 0.983 | -0.078 | 0.693 | 0.968 | -0.045 | 0.818 | 0.997 | 0.091 | 0.642 | 0.947 | 0.023 | 0.908 | 1.000 |
| **C2** | -0.320 | 0.098 | 0.868 | -0.143 | 0.467 | 0.983 | 0.002 | 0.994 | 0.994 | 0.012 | 0.952 | 0.997 | 0.038 | 0.849 | 0.958 | -0.079 | 0.689 | 1.000 |
| **P1** | -0.226 | 0.246 | 0.868 | -0.217 | 0.267 | 0.983 | 0.421 | 0.026 | 0.457 | -0.062 | 0.754 | 0.997 | 0.364 | 0.058 | 0.547 | 0.211 | 0.280 | 0.949 |
| **P2** | -0.176 | 0.370 | 0.868 | 0.030 | 0.882 | 0.983 | 0.320 | 0.098 | 0.728 | -0.070 | 0.724 | 0.997 | 0.073 | 0.712 | 0.947 | -0.240 | 0.217 | 0.949 |
| **AF3** | 0.056 | 0.777 | 0.967 | -0.028 | 0.886 | 0.983 | -0.058 | 0.769 | 0.973 | -0.025 | 0.901 | 0.997 | 0.026 | 0.895 | 0.984 | -0.229 | 0.241 | 0.949 |
| **AF4** | 0.141 | 0.474 | 0.868 | 0.065 | 0.743 | 0.983 | 0.220 | 0.259 | 0.842 | -0.212 | 0.278 | 0.997 | -0.224 | 0.250 | 0.928 | -0.102 | 0.603 | 0.983 |
| **FC3** | -0.094 | 0.634 | 0.963 | 0.063 | 0.750 | 0.983 | -0.045 | 0.818 | 0.973 | 0.032 | 0.871 | 0.997 | 0.079 | 0.689 | 0.947 | 0.017 | 0.932 | 1.000 |
| **FC4** | -0.014 | 0.946 | 0.991 | 0.005 | 0.981 | 0.983 | -0.024 | 0.906 | 0.979 | 0.215 | 0.272 | 0.997 | 0.241 | 0.216 | 0.928 | 0.304 | 0.116 | 0.949 |
| **CP3** | 0.250 | 0.199 | 0.868 | -0.236 | 0.225 | 0.983 | 0.133 | 0.498 | 0.913 | -0.058 | 0.769 | 0.997 | 0.231 | 0.236 | 0.928 | 0.170 | 0.386 | 0.949 |
| **CP4** | -0.381 | 0.046 | 0.822 | -0.137 | 0.486 | 0.983 | -0.079 | 0.687 | 0.968 | -0.129 | 0.513 | 0.997 | 0.005 | 0.981 | 0.999 | -0.024 | 0.904 | 1.000 |
| **PO3** | -0.003 | 0.988 | 0.994 | -0.033 | 0.866 | 0.983 | 0.524 | 0.005 | 0.207 | -0.006 | 0.977 | 0.997 | 0.224 | 0.251 | 0.928 | -0.037 | 0.853 | 1.000 |
| **PO4** | 0.081 | 0.681 | 0.967 | -0.030 | 0.882 | 0.983 | 0.216 | 0.268 | 0.842 | -0.108 | 0.584 | 0.997 | 0.079 | 0.687 | 0.947 | 0.269 | 0.166 | 0.949 |
| **F5** | 0.193 | 0.323 | 0.868 | -0.022 | 0.910 | 0.983 | -0.060 | 0.762 | 0.973 | 0.072 | 0.716 | 0.997 | 0.152 | 0.440 | 0.947 | -0.120 | 0.542 | 0.983 |
| **F6** | 0.149 | 0.448 | 0.868 | -0.163 | 0.405 | 0.983 | 0.289 | 0.136 | 0.728 | 0.072 | 0.716 | 0.997 | -0.222 | 0.255 | 0.928 | -0.008 | 0.970 | 1.000 |
| **C5** | 0.309 | 0.109 | 0.868 | 0.080 | 0.685 | 0.983 | 0.096 | 0.625 | 0.948 | 0.065 | 0.741 | 0.997 | -0.097 | 0.623 | 0.947 | 0.201 | 0.304 | 0.949 |
| **C6** | -0.036 | 0.855 | 0.967 | -0.014 | 0.943 | 0.983 | 0.136 | 0.489 | 0.913 | -0.006 | 0.977 | 0.997 | 0.054 | 0.786 | 0.947 | 0.081 | 0.681 | 1.000 |
| **P5** | -0.261 | 0.179 | 0.868 | -0.093 | 0.638 | 0.983 | 0.338 | 0.079 | 0.728 | -0.147 | 0.455 | 0.997 | 0.188 | 0.336 | 0.947 | 0.169 | 0.389 | 0.949 |
| **P6** | 0.102 | 0.603 | 0.960 | -0.165 | 0.400 | 0.983 | 0.172 | 0.379 | 0.913 | -0.090 | 0.649 | 0.997 | -0.157 | 0.425 | 0.947 | 0.000 | 1.000 | 1.000 |
| **Fpz** | -0.032 | 0.873 | 0.967 | 0.221 | 0.257 | 0.983 | -0.144 | 0.462 | 0.913 | -0.211 | 0.280 | 0.997 | -0.065 | 0.743 | 0.947 | -0.288 | 0.136 | 0.949 |
| **CPz** | -0.183 | 0.350 | 0.868 | 0.123 | 0.531 | 0.983 | -0.053 | 0.790 | 0.973 | -0.179 | 0.361 | 0.997 | 0.402 | 0.035 | 0.547 | -0.051 | 0.797 | 1.000 |
| **POz** | -0.002 | 0.994 | 0.994 | -0.071 | 0.718 | 0.983 | -0.016 | 0.935 | 0.979 | -0.039 | 0.842 | 0.997 | 0.182 | 0.353 | 0.947 | 0.114 | 0.561 | 0.983 |
| **FCz** | -0.152 | 0.440 | 0.868 | 0.043 | 0.827 | 0.983 | 0.234 | 0.229 | 0.842 | 0.423 | 0.026 | 0.679 | 0.134 | 0.496 | 0.947 | -0.091 | 0.644 | 1.000 |
|  | **C-TCT** | | | | | | | | |  | | | | | | | | |
|  | **Beta** | | | | | | | | | **Gamma** | | | | | | | | |
|  | **ΔHEP and Δpower** | | | **ΔHEP and ΔITC** | | | **Δpower and ΔITC** | | | **ΔHEP and Δpower** | | | **ΔHEP and ΔITC** | | | **Δpower and ΔITC** | | |
| **Channel** | **rho** | **p** | **p_FDR_** | **rho** | **p** | **p_FDR_** | **rho** | **p** | **p_FDR_** | **rho** | **p** | **p_FDR_** | **rho** | **p** | **p_FDR_** | **rho** | **p** | **p_FDR_** |
| **Fp1** | 0.091 | 0.642 | 0.857 | -0.068 | 0.731 | 0.961 | 0.181 | 0.355 | 0.871 | 0.206 | 0.292 | 0.847 | 0.041 | 0.836 | 0.972 | -0.040 | 0.840 | 0.962 |
| **Fp2** | -0.036 | 0.855 | 0.953 | 0.132 | 0.502 | 0.961 | 0.056 | 0.777 | 0.904 | 0.085 | 0.665 | 0.943 | 0.055 | 0.782 | 0.972 | -0.002 | 0.992 | 0.992 |
| **F3** | 0.452 | 0.017 | 0.213 | 0.165 | 0.399 | 0.961 | -0.124 | 0.529 | 0.904 | 0.024 | 0.904 | 0.992 | -0.175 | 0.373 | 0.863 | -0.079 | 0.689 | 0.962 |
| **F4** | -0.033 | 0.868 | 0.953 | 0.018 | 0.930 | 0.961 | -0.099 | 0.615 | 0.904 | 0.049 | 0.805 | 0.992 | 0.013 | 0.950 | 0.995 | -0.206 | 0.292 | 0.838 |
| **C3** | 0.119 | 0.546 | 0.774 | -0.258 | 0.185 | 0.961 | 0.110 | 0.576 | 0.904 | -0.174 | 0.374 | 0.847 | -0.278 | 0.152 | 0.750 | 0.196 | 0.315 | 0.838 |
| **C4** | 0.305 | 0.115 | 0.460 | 0.408 | 0.032 | 0.632 | 0.180 | 0.359 | 0.871 | -0.029 | 0.884 | 0.992 | 0.277 | 0.153 | 0.750 | -0.074 | 0.708 | 0.962 |
| **P3** | -0.270 | 0.164 | 0.460 | 0.125 | 0.525 | 0.961 | -0.148 | 0.451 | 0.904 | 0.178 | 0.364 | 0.847 | -0.025 | 0.901 | 0.972 | -0.036 | 0.855 | 0.962 |
| **P4** | -0.227 | 0.244 | 0.480 | 0.172 | 0.379 | 0.961 | -0.228 | 0.242 | 0.815 | -0.367 | 0.056 | 0.426 | 0.100 | 0.611 | 0.949 | -0.180 | 0.358 | 0.838 |
| **Fz** | 0.456 | 0.016 | 0.213 | -0.386 | 0.043 | 0.632 | 0.042 | 0.831 | 0.904 | 0.307 | 0.112 | 0.548 | -0.234 | 0.229 | 0.791 | -0.418 | 0.028 | 0.609 |
| **Cz** | 0.240 | 0.218 | 0.480 | 0.043 | 0.829 | 0.961 | -0.055 | 0.782 | 0.904 | -0.146 | 0.456 | 0.899 | -0.206 | 0.292 | 0.803 | 0.082 | 0.677 | 0.962 |
| **Pz** | -0.442 | 0.019 | 0.213 | -0.179 | 0.361 | 0.961 | -0.051 | 0.797 | 0.904 | -0.255 | 0.191 | 0.816 | -0.072 | 0.716 | 0.960 | 0.234 | 0.230 | 0.838 |
| **AFz** | 0.403 | 0.034 | 0.285 | 0.011 | 0.954 | 0.961 | 0.212 | 0.278 | 0.815 | 0.189 | 0.333 | 0.847 | -0.190 | 0.332 | 0.849 | -0.313 | 0.106 | 0.774 |
| **FC1** | 0.224 | 0.251 | 0.480 | 0.163 | 0.405 | 0.961 | -0.124 | 0.529 | 0.904 | -0.115 | 0.559 | 0.899 | 0.148 | 0.450 | 0.942 | -0.024 | 0.906 | 0.972 |
| **FC2** | -0.033 | 0.866 | 0.953 | -0.074 | 0.708 | 0.961 | -0.050 | 0.799 | 0.904 | -0.049 | 0.805 | 0.992 | 0.120 | 0.542 | 0.949 | 0.032 | 0.873 | 0.962 |
| **CP1** | 0.074 | 0.706 | 0.887 | 0.092 | 0.640 | 0.961 | -0.001 | 0.999 | 0.999 | 0.422 | 0.026 | 0.426 | -0.250 | 0.199 | 0.791 | 0.048 | 0.807 | 0.962 |
| **CP2** | -0.028 | 0.888 | 0.953 | -0.010 | 0.961 | 0.961 | -0.244 | 0.210 | 0.815 | -0.314 | 0.104 | 0.548 | -0.027 | 0.893 | 0.972 | 0.038 | 0.849 | 0.962 |
| **FC5** | -0.044 | 0.825 | 0.953 | -0.094 | 0.633 | 0.961 | -0.252 | 0.195 | 0.815 | -0.013 | 0.950 | 0.992 | 0.393 | 0.039 | 0.750 | -0.182 | 0.352 | 0.838 |
| **FC6** | -0.302 | 0.119 | 0.460 | -0.021 | 0.915 | 0.961 | -0.166 | 0.396 | 0.871 | -0.149 | 0.446 | 0.899 | 0.240 | 0.217 | 0.791 | -0.447 | 0.018 | 0.609 |
| **CP5** | 0.346 | 0.072 | 0.396 | -0.030 | 0.879 | 0.961 | -0.027 | 0.890 | 0.913 | -0.079 | 0.689 | 0.948 | -0.306 | 0.113 | 0.750 | -0.017 | 0.932 | 0.977 |
| **CP6** | -0.279 | 0.151 | 0.460 | 0.066 | 0.739 | 0.961 | -0.181 | 0.355 | 0.871 | -0.188 | 0.337 | 0.847 | -0.149 | 0.446 | 0.942 | -0.004 | 0.983 | 0.992 |
| **F1** | 0.285 | 0.141 | 0.460 | -0.049 | 0.805 | 0.961 | -0.213 | 0.274 | 0.815 | 0.238 | 0.223 | 0.816 | -0.044 | 0.825 | 0.972 | -0.344 | 0.073 | 0.749 |
| **F2** | 0.351 | 0.067 | 0.396 | -0.120 | 0.542 | 0.961 | -0.065 | 0.743 | 0.904 | 0.238 | 0.222 | 0.816 | 0.028 | 0.888 | 0.972 | -0.031 | 0.875 | 0.962 |
| **C1** | -0.050 | 0.799 | 0.953 | -0.078 | 0.691 | 0.961 | 0.227 | 0.244 | 0.815 | -0.126 | 0.522 | 0.899 | -0.209 | 0.286 | 0.803 | -0.039 | 0.842 | 0.962 |
| **C2** | 0.193 | 0.324 | 0.595 | 0.192 | 0.327 | 0.961 | 0.253 | 0.193 | 0.815 | -0.368 | 0.055 | 0.426 | -0.369 | 0.054 | 0.750 | 0.207 | 0.288 | 0.838 |
| **P1** | -0.499 | 0.008 | 0.213 | 0.144 | 0.463 | 0.961 | 0.057 | 0.771 | 0.904 | -0.030 | 0.882 | 0.992 | 0.071 | 0.720 | 0.960 | 0.066 | 0.737 | 0.962 |
| **P2** | -0.328 | 0.089 | 0.435 | 0.013 | 0.950 | 0.961 | -0.259 | 0.183 | 0.815 | -0.334 | 0.083 | 0.521 | -0.024 | 0.906 | 0.972 | -0.269 | 0.166 | 0.838 |
| **AF3** | 0.227 | 0.245 | 0.480 | 0.072 | 0.716 | 0.961 | -0.221 | 0.257 | 0.815 | 0.041 | 0.838 | 0.992 | 0.005 | 0.981 | 0.997 | -0.190 | 0.330 | 0.838 |
| **AF4** | 0.011 | 0.954 | 0.954 | 0.157 | 0.425 | 0.961 | 0.040 | 0.840 | 0.904 | 0.133 | 0.498 | 0.899 | 0.118 | 0.549 | 0.949 | -0.332 | 0.085 | 0.749 |
| **FC3** | -0.017 | 0.932 | 0.954 | -0.118 | 0.548 | 0.961 | -0.300 | 0.121 | 0.815 | -0.008 | 0.970 | 0.992 | -0.293 | 0.130 | 0.750 | 0.245 | 0.208 | 0.838 |
| **FC4** | 0.076 | 0.702 | 0.887 | 0.210 | 0.283 | 0.961 | -0.114 | 0.563 | 0.904 | -0.119 | 0.546 | 0.899 | 0.133 | 0.498 | 0.949 | 0.044 | 0.825 | 0.962 |
| **CP3** | -0.016 | 0.935 | 0.954 | 0.183 | 0.350 | 0.961 | -0.039 | 0.842 | 0.904 | 0.373 | 0.052 | 0.426 | 0.050 | 0.801 | 0.972 | 0.138 | 0.482 | 0.923 |
| **CP4** | 0.167 | 0.394 | 0.619 | 0.562 | 0.002 | 0.096 | 0.042 | 0.831 | 0.904 | -0.175 | 0.373 | 0.847 | 0.001 | 0.997 | 0.997 | -0.079 | 0.687 | 0.962 |
| **PO3** | -0.231 | 0.236 | 0.480 | 0.154 | 0.433 | 0.961 | -0.067 | 0.733 | 0.904 | -0.101 | 0.609 | 0.924 | -0.285 | 0.142 | 0.750 | -0.178 | 0.362 | 0.838 |
| **PO4** | -0.256 | 0.189 | 0.471 | -0.028 | 0.888 | 0.961 | 0.069 | 0.727 | 0.904 | -0.363 | 0.058 | 0.426 | 0.088 | 0.657 | 0.949 | -0.252 | 0.195 | 0.838 |
| **F5** | 0.108 | 0.582 | 0.800 | 0.111 | 0.574 | 0.961 | -0.100 | 0.613 | 0.904 | -0.140 | 0.475 | 0.899 | -0.232 | 0.234 | 0.791 | 0.054 | 0.786 | 0.962 |
| **F6** | -0.253 | 0.193 | 0.471 | -0.010 | 0.959 | 0.961 | 0.312 | 0.106 | 0.815 | -0.111 | 0.572 | 0.899 | 0.089 | 0.651 | 0.949 | -0.154 | 0.433 | 0.866 |
| **C5** | -0.178 | 0.362 | 0.619 | -0.030 | 0.882 | 0.961 | -0.217 | 0.265 | 0.815 | 0.027 | 0.890 | 0.992 | -0.304 | 0.116 | 0.750 | -0.062 | 0.754 | 0.962 |
| **C6** | -0.128 | 0.514 | 0.754 | 0.116 | 0.555 | 0.961 | -0.027 | 0.893 | 0.913 | -0.093 | 0.636 | 0.934 | 0.184 | 0.347 | 0.849 | -0.198 | 0.312 | 0.838 |
| **P5** | -0.268 | 0.167 | 0.460 | 0.342 | 0.076 | 0.834 | -0.223 | 0.253 | 0.815 | -0.047 | 0.812 | 0.992 | -0.085 | 0.667 | 0.949 | -0.048 | 0.810 | 0.962 |
| **P6** | -0.394 | 0.039 | 0.285 | 0.056 | 0.775 | 0.961 | -0.090 | 0.649 | 0.904 | 0.002 | 0.992 | 0.992 | 0.206 | 0.292 | 0.803 | -0.164 | 0.402 | 0.866 |
| **Fpz** | 0.153 | 0.436 | 0.662 | -0.047 | 0.814 | 0.961 | 0.213 | 0.274 | 0.815 | 0.170 | 0.385 | 0.847 | -0.084 | 0.669 | 0.949 | -0.157 | 0.423 | 0.866 |
| **CPz** | -0.170 | 0.386 | 0.619 | -0.092 | 0.640 | 0.961 | 0.172 | 0.379 | 0.871 | 0.016 | 0.935 | 0.992 | -0.461 | 0.014 | 0.629 | 0.389 | 0.042 | 0.609 |
| **POz** | -0.285 | 0.142 | 0.460 | -0.136 | 0.488 | 0.961 | -0.111 | 0.574 | 0.904 | -0.411 | 0.031 | 0.426 | 0.120 | 0.542 | 0.949 | 0.078 | 0.691 | 0.962 |
| **FCz** | 0.169 | 0.389 | 0.619 | 0.059 | 0.767 | 0.961 | -0.219 | 0.262 | 0.815 | -0.220 | 0.259 | 0.847 | 0.085 | 0.665 | 0.949 | 0.245 | 0.209 | 0.838 |

**Supplementary Table 23. Correlations between respiratory phase-related differences (Δ = exhalation − inhalation) in heartbeat-evoked potential, heartbeat-related power, and heartbeat-related inter-trial coherence across frequency bands and tasks.** Abbreviations: HCT = heartbeat counting task, C-TCT = cardiac-tone counting task, HEP = heartbeat-evoked potential, ITC = inter-trial coherence, FDR = false discovery rate.

|  | **HCT** | | | | | | | | | | | | | | | | | |
| --- | --- | --- | --- | --- | --- | --- | --- | --- | --- | --- | --- | --- | --- | --- | --- | --- | --- | --- |
|  | **Theta** | | | | | | | | | | | | | | | | | |
|  | **ΔHEP and Δpower** | | | | | | **ΔHEP and ΔITC** | | | | | | **Δpower and ΔITC** | | | | | |
|  | **Inhalation** | | | **Exhalation** | | | **Inhalation** | | | **Exhalation** | | | **Inhalation** | | | **Exhalation** | | |
| **Channel** | **rho** | **p** | **p_FDR_** | **rho** | **p** | **p_FDR_** | **rho** | **p** | **p_FDR_** | **rho** | **p** | **p_FDR_** | **rho** | **p** | **p_FDR_** | **rho** | **p** | **p_FDR_** |
| **Fp1** | 0.090 | 0.647 | 0.977 | -0.015 | 0.939 | 0.963 | -0.026 | 0.897 | 0.940 | -0.106 | 0.589 | 0.836 | 0.185 | 0.344 | 0.737 | -0.154 | 0.433 | 0.901 |
| **Fp2** | -0.048 | 0.810 | 0.977 | -0.109 | 0.578 | 0.959 | 0.203 | 0.300 | 0.596 | -0.259 | 0.182 | 0.699 | 0.175 | 0.371 | 0.737 | 0.263 | 0.175 | 0.878 |
| **F3** | -0.195 | 0.319 | 0.787 | 0.118 | 0.549 | 0.959 | -0.337 | 0.080 | 0.476 | -0.125 | 0.525 | 0.836 | 0.198 | 0.312 | 0.737 | 0.060 | 0.760 | 0.918 |
| **F4** | 0.078 | 0.691 | 0.977 | -0.112 | 0.570 | 0.959 | -0.336 | 0.081 | 0.476 | -0.112 | 0.570 | 0.836 | 0.141 | 0.472 | 0.737 | 0.041 | 0.836 | 0.943 |
| **C3** | 0.039 | 0.844 | 0.977 | -0.373 | 0.051 | 0.747 | 0.209 | 0.284 | 0.596 | -0.061 | 0.756 | 0.894 | 0.109 | 0.578 | 0.778 | -0.002 | 0.994 | 0.994 |
| **C4** | 0.085 | 0.667 | 0.977 | 0.000 | 1.000 | 1.000 | -0.187 | 0.339 | 0.596 | 0.085 | 0.665 | 0.860 | 0.022 | 0.912 | 0.934 | -0.255 | 0.190 | 0.878 |
| **P3** | -0.105 | 0.595 | 0.977 | -0.117 | 0.551 | 0.959 | 0.139 | 0.479 | 0.702 | -0.167 | 0.393 | 0.836 | 0.144 | 0.463 | 0.737 | 0.229 | 0.239 | 0.878 |
| **P4** | 0.163 | 0.407 | 0.814 | -0.186 | 0.341 | 0.959 | 0.007 | 0.972 | 0.972 | -0.087 | 0.659 | 0.860 | 0.362 | 0.059 | 0.351 | -0.184 | 0.347 | 0.878 |
| **Fz** | -0.204 | 0.296 | 0.787 | -0.077 | 0.697 | 0.959 | -0.118 | 0.549 | 0.711 | -0.092 | 0.640 | 0.860 | 0.185 | 0.344 | 0.737 | -0.075 | 0.704 | 0.918 |
| **Cz** | 0.243 | 0.212 | 0.777 | -0.093 | 0.638 | 0.959 | -0.221 | 0.257 | 0.596 | -0.133 | 0.498 | 0.836 | -0.296 | 0.127 | 0.558 | 0.026 | 0.895 | 0.960 |
| **Pz** | 0.368 | 0.055 | 0.674 | -0.372 | 0.052 | 0.747 | 0.105 | 0.593 | 0.746 | -0.255 | 0.191 | 0.699 | 0.412 | 0.030 | 0.332 | -0.180 | 0.359 | 0.878 |
| **AFz** | -0.259 | 0.183 | 0.731 | -0.152 | 0.440 | 0.959 | -0.084 | 0.671 | 0.820 | -0.400 | 0.036 | 0.562 | 0.245 | 0.209 | 0.706 | 0.016 | 0.937 | 0.981 |
| **FC1** | -0.305 | 0.115 | 0.706 | 0.077 | 0.695 | 0.959 | 0.222 | 0.255 | 0.596 | -0.055 | 0.780 | 0.894 | -0.258 | 0.185 | 0.706 | -0.057 | 0.773 | 0.918 |
| **FC2** | 0.041 | 0.838 | 0.977 | -0.211 | 0.280 | 0.959 | -0.302 | 0.119 | 0.476 | 0.165 | 0.399 | 0.836 | 0.198 | 0.311 | 0.737 | 0.107 | 0.587 | 0.918 |
| **CP1** | 0.048 | 0.810 | 0.977 | -0.366 | 0.056 | 0.747 | -0.303 | 0.117 | 0.476 | -0.177 | 0.367 | 0.836 | 0.106 | 0.591 | 0.778 | 0.167 | 0.394 | 0.901 |
| **CP2** | -0.117 | 0.553 | 0.974 | 0.314 | 0.104 | 0.747 | -0.045 | 0.820 | 0.899 | -0.139 | 0.479 | 0.836 | 0.037 | 0.853 | 0.934 | -0.218 | 0.263 | 0.878 |
| **FC5** | -0.423 | 0.026 | 0.674 | -0.045 | 0.818 | 0.963 | 0.019 | 0.923 | 0.945 | -0.395 | 0.038 | 0.562 | 0.028 | 0.886 | 0.934 | 0.002 | 0.992 | 0.994 |
| **FC6** | -0.169 | 0.388 | 0.813 | 0.275 | 0.156 | 0.747 | -0.288 | 0.137 | 0.503 | 0.179 | 0.361 | 0.836 | 0.185 | 0.344 | 0.737 | 0.323 | 0.093 | 0.878 |
| **CP5** | 0.006 | 0.977 | 0.977 | -0.114 | 0.563 | 0.959 | -0.066 | 0.739 | 0.834 | -0.052 | 0.792 | 0.894 | 0.123 | 0.533 | 0.778 | -0.129 | 0.513 | 0.901 |
| **CP6** | -0.183 | 0.350 | 0.787 | -0.175 | 0.373 | 0.959 | -0.123 | 0.533 | 0.711 | 0.130 | 0.507 | 0.836 | -0.159 | 0.418 | 0.737 | -0.123 | 0.533 | 0.901 |
| **F1** | -0.386 | 0.043 | 0.674 | 0.023 | 0.908 | 0.963 | -0.076 | 0.702 | 0.827 | 0.109 | 0.580 | 0.836 | -0.101 | 0.607 | 0.778 | -0.132 | 0.502 | 0.901 |
| **F2** | -0.205 | 0.295 | 0.787 | -0.096 | 0.625 | 0.959 | -0.374 | 0.051 | 0.476 | -0.135 | 0.493 | 0.836 | -0.022 | 0.910 | 0.934 | 0.066 | 0.739 | 0.918 |
| **C1** | -0.038 | 0.847 | 0.977 | -0.031 | 0.877 | 0.963 | -0.191 | 0.329 | 0.596 | -0.361 | 0.060 | 0.656 | -0.137 | 0.486 | 0.737 | 0.079 | 0.687 | 0.918 |
| **C2** | -0.032 | 0.871 | 0.977 | 0.077 | 0.697 | 0.959 | -0.328 | 0.089 | 0.476 | 0.019 | 0.926 | 0.988 | -0.097 | 0.623 | 0.778 | -0.187 | 0.339 | 0.878 |
| **P1** | 0.146 | 0.458 | 0.876 | -0.308 | 0.111 | 0.747 | 0.168 | 0.391 | 0.637 | 0.004 | 0.986 | 0.988 | 0.423 | 0.026 | 0.332 | -0.065 | 0.741 | 0.918 |
| **P2** | 0.182 | 0.352 | 0.787 | -0.152 | 0.440 | 0.959 | 0.189 | 0.333 | 0.596 | 0.111 | 0.572 | 0.836 | 0.344 | 0.074 | 0.361 | -0.458 | 0.015 | 0.669 |
| **AF3** | -0.130 | 0.509 | 0.933 | 0.269 | 0.165 | 0.747 | 0.153 | 0.436 | 0.662 | -0.282 | 0.146 | 0.664 | 0.378 | 0.048 | 0.351 | 0.064 | 0.746 | 0.918 |
| **AF4** | 0.009 | 0.963 | 0.977 | -0.055 | 0.780 | 0.963 | -0.257 | 0.187 | 0.548 | -0.290 | 0.134 | 0.664 | 0.174 | 0.376 | 0.737 | 0.243 | 0.212 | 0.878 |
| **FC3** | -0.090 | 0.647 | 0.977 | 0.077 | 0.697 | 0.959 | 0.172 | 0.379 | 0.637 | -0.184 | 0.346 | 0.836 | 0.050 | 0.799 | 0.925 | 0.094 | 0.634 | 0.918 |
| **FC4** | 0.046 | 0.816 | 0.977 | -0.015 | 0.941 | 0.963 | -0.357 | 0.063 | 0.476 | 0.397 | 0.037 | 0.562 | 0.157 | 0.425 | 0.737 | -0.259 | 0.182 | 0.878 |
| **CP3** | 0.269 | 0.165 | 0.728 | -0.344 | 0.073 | 0.747 | 0.041 | 0.838 | 0.899 | -0.333 | 0.083 | 0.664 | -0.016 | 0.935 | 0.935 | 0.052 | 0.792 | 0.918 |
| **CP4** | 0.316 | 0.102 | 0.706 | -0.050 | 0.799 | 0.963 | 0.305 | 0.115 | 0.476 | 0.293 | 0.130 | 0.664 | 0.036 | 0.857 | 0.934 | -0.366 | 0.056 | 0.878 |
| **PO3** | 0.307 | 0.113 | 0.706 | -0.104 | 0.597 | 0.959 | 0.472 | 0.012 | 0.476 | -0.145 | 0.460 | 0.836 | 0.356 | 0.064 | 0.351 | -0.221 | 0.257 | 0.878 |
| **PO4** | 0.283 | 0.144 | 0.706 | -0.099 | 0.617 | 0.959 | 0.372 | 0.052 | 0.476 | 0.279 | 0.151 | 0.664 | 0.235 | 0.228 | 0.717 | -0.183 | 0.350 | 0.878 |
| **F5** | -0.359 | 0.061 | 0.674 | 0.277 | 0.153 | 0.747 | -0.124 | 0.529 | 0.711 | -0.181 | 0.355 | 0.836 | 0.166 | 0.396 | 0.737 | -0.264 | 0.174 | 0.878 |
| **F6** | 0.006 | 0.977 | 0.977 | 0.267 | 0.170 | 0.747 | -0.417 | 0.028 | 0.476 | -0.003 | 0.988 | 0.988 | 0.111 | 0.572 | 0.778 | 0.026 | 0.895 | 0.960 |
| **C5** | 0.070 | 0.722 | 0.977 | -0.034 | 0.862 | 0.963 | 0.264 | 0.174 | 0.548 | -0.221 | 0.257 | 0.808 | -0.146 | 0.456 | 0.737 | -0.198 | 0.311 | 0.878 |
| **C6** | -0.008 | 0.968 | 0.977 | -0.063 | 0.750 | 0.963 | -0.204 | 0.297 | 0.596 | 0.006 | 0.977 | 0.988 | -0.063 | 0.750 | 0.892 | 0.147 | 0.455 | 0.901 |
| **P5** | -0.184 | 0.347 | 0.787 | 0.036 | 0.855 | 0.963 | 0.072 | 0.714 | 0.827 | -0.070 | 0.722 | 0.891 | 0.248 | 0.202 | 0.706 | 0.181 | 0.356 | 0.878 |
| **P6** | 0.180 | 0.358 | 0.787 | -0.251 | 0.196 | 0.786 | 0.118 | 0.549 | 0.711 | 0.292 | 0.131 | 0.664 | 0.093 | 0.636 | 0.778 | -0.195 | 0.318 | 0.878 |
| **Fpz** | 0.022 | 0.910 | 0.977 | -0.084 | 0.669 | 0.959 | 0.194 | 0.320 | 0.596 | -0.224 | 0.250 | 0.808 | 0.375 | 0.050 | 0.351 | 0.135 | 0.491 | 0.901 |
| **CPz** | -0.077 | 0.695 | 0.977 | 0.107 | 0.587 | 0.959 | -0.269 | 0.165 | 0.548 | 0.018 | 0.928 | 0.988 | 0.513 | 0.006 | 0.257 | -0.129 | 0.513 | 0.901 |
| **POz** | 0.229 | 0.239 | 0.787 | -0.097 | 0.623 | 0.959 | 0.157 | 0.425 | 0.662 | -0.131 | 0.504 | 0.836 | 0.444 | 0.019 | 0.332 | -0.073 | 0.712 | 0.918 |
| **FCz** | 0.285 | 0.142 | 0.706 | -0.044 | 0.825 | 0.963 | 0.249 | 0.201 | 0.551 | -0.068 | 0.729 | 0.891 | -0.161 | 0.410 | 0.737 | -0.101 | 0.609 | 0.918 |
|  | **HCT** | | | | | | | | | | | | | | | | | |
|  | **Alpha** | | | | | | | | | | | | | | | | | |
|  | **ΔHEP and Δpower** | | | | | | **ΔHEP and ΔITC** | | | | | | **Δpower and ΔITC** | | | | | |
|  | **Inhalation** | | | **Exhalation** | | | **Inhalation** | | | **Exhalation** | | | **Inhalation** | | | **Exhalation** | | |
| **Channel** | **rho** | **p** | **p_FDR_** | **rho** | **p** | **p_FDR_** | **rho** | **p** | **p_FDR_** | **rho** | **p** | **p_FDR_** | **rho** | **p** | **p_FDR_** | **rho** | **p** | **p_FDR_** |
| **Fp1** | 0.143 | 0.467 | 0.819 | -0.113 | 0.564 | 0.990 | 0.018 | 0.930 | 0.948 | 0.010 | 0.961 | 0.981 | -0.055 | 0.780 | 0.972 | 0.085 | 0.665 | 0.947 |
| **Fp2** | 0.194 | 0.320 | 0.819 | -0.015 | 0.941 | 0.997 | -0.078 | 0.693 | 0.847 | -0.078 | 0.693 | 0.981 | -0.201 | 0.303 | 0.972 | 0.070 | 0.724 | 0.947 |
| **F3** | 0.220 | 0.259 | 0.819 | 0.084 | 0.669 | 0.990 | 0.132 | 0.500 | 0.818 | -0.069 | 0.727 | 0.981 | -0.009 | 0.966 | 0.972 | -0.028 | 0.888 | 0.947 |
| **F4** | 0.207 | 0.288 | 0.819 | 0.170 | 0.386 | 0.990 | -0.302 | 0.119 | 0.606 | -0.043 | 0.829 | 0.981 | -0.339 | 0.078 | 0.972 | 0.188 | 0.336 | 0.947 |
| **C3** | -0.071 | 0.720 | 0.906 | -0.061 | 0.756 | 0.990 | 0.234 | 0.229 | 0.704 | 0.178 | 0.364 | 0.981 | 0.090 | 0.647 | 0.972 | 0.130 | 0.509 | 0.947 |
| **C4** | 0.319 | 0.098 | 0.819 | -0.019 | 0.923 | 0.997 | -0.208 | 0.287 | 0.704 | -0.106 | 0.591 | 0.981 | 0.313 | 0.105 | 0.972 | -0.233 | 0.231 | 0.947 |
| **P3** | -0.284 | 0.143 | 0.819 | -0.136 | 0.489 | 0.990 | 0.402 | 0.035 | 0.409 | -0.013 | 0.950 | 0.981 | -0.061 | 0.756 | 0.972 | 0.074 | 0.708 | 0.947 |
| **P4** | -0.150 | 0.445 | 0.819 | -0.054 | 0.786 | 0.990 | 0.155 | 0.430 | 0.785 | 0.163 | 0.407 | 0.981 | -0.009 | 0.966 | 0.972 | -0.093 | 0.636 | 0.947 |
| **Fz** | -0.099 | 0.615 | 0.906 | 0.059 | 0.767 | 0.990 | -0.332 | 0.085 | 0.606 | 0.363 | 0.058 | 0.918 | -0.020 | 0.921 | 0.972 | 0.143 | 0.465 | 0.947 |
| **Cz** | -0.119 | 0.546 | 0.889 | -0.164 | 0.404 | 0.990 | 0.013 | 0.948 | 0.948 | -0.005 | 0.981 | 0.981 | -0.107 | 0.585 | 0.972 | 0.027 | 0.890 | 0.947 |
| **Pz** | -0.050 | 0.799 | 0.922 | -0.360 | 0.061 | 0.990 | 0.114 | 0.561 | 0.847 | -0.357 | 0.063 | 0.918 | 0.054 | 0.784 | 0.972 | 0.223 | 0.253 | 0.947 |
| **AFz** | 0.047 | 0.814 | 0.922 | -0.077 | 0.695 | 0.990 | -0.269 | 0.165 | 0.606 | 0.169 | 0.388 | 0.981 | -0.026 | 0.897 | 0.972 | 0.028 | 0.888 | 0.947 |
| **FC1** | 0.083 | 0.675 | 0.906 | -0.105 | 0.593 | 0.990 | 0.105 | 0.595 | 0.847 | -0.168 | 0.391 | 0.981 | 0.041 | 0.836 | 0.972 | 0.073 | 0.712 | 0.947 |
| **FC2** | -0.165 | 0.399 | 0.819 | -0.070 | 0.724 | 0.990 | -0.132 | 0.502 | 0.818 | -0.033 | 0.868 | 0.981 | -0.174 | 0.374 | 0.972 | 0.165 | 0.400 | 0.947 |
| **CP1** | -0.061 | 0.758 | 0.922 | 0.092 | 0.640 | 0.990 | -0.026 | 0.895 | 0.940 | -0.171 | 0.382 | 0.981 | -0.148 | 0.451 | 0.972 | -0.039 | 0.844 | 0.947 |
| **CP2** | -0.184 | 0.347 | 0.819 | 0.351 | 0.068 | 0.990 | -0.201 | 0.304 | 0.704 | -0.065 | 0.743 | 0.981 | -0.033 | 0.868 | 0.972 | -0.108 | 0.582 | 0.947 |
| **FC5** | -0.274 | 0.158 | 0.819 | 0.210 | 0.282 | 0.990 | 0.176 | 0.370 | 0.730 | -0.069 | 0.727 | 0.981 | 0.102 | 0.605 | 0.972 | -0.142 | 0.468 | 0.947 |
| **FC6** | 0.072 | 0.716 | 0.906 | 0.173 | 0.377 | 0.990 | -0.563 | 0.002 | 0.094 | -0.173 | 0.377 | 0.981 | 0.158 | 0.421 | 0.972 | 0.186 | 0.341 | 0.947 |
| **CP5** | 0.180 | 0.358 | 0.819 | -0.107 | 0.587 | 0.990 | -0.092 | 0.640 | 0.847 | -0.221 | 0.258 | 0.981 | 0.099 | 0.617 | 0.972 | 0.003 | 0.990 | 0.990 |
| **CP6** | 0.080 | 0.685 | 0.906 | -0.026 | 0.895 | 0.997 | -0.049 | 0.805 | 0.932 | -0.063 | 0.750 | 0.981 | 0.036 | 0.855 | 0.972 | -0.169 | 0.389 | 0.947 |
| **F1** | 0.019 | 0.926 | 0.935 | -0.039 | 0.842 | 0.997 | -0.270 | 0.165 | 0.606 | -0.154 | 0.431 | 0.981 | -0.037 | 0.853 | 0.972 | 0.045 | 0.820 | 0.947 |
| **F2** | 0.215 | 0.272 | 0.819 | -0.092 | 0.640 | 0.990 | -0.397 | 0.037 | 0.409 | -0.083 | 0.675 | 0.981 | -0.075 | 0.704 | 0.972 | 0.239 | 0.220 | 0.947 |
| **C1** | 0.025 | 0.899 | 0.935 | -0.002 | 0.992 | 0.997 | -0.026 | 0.895 | 0.940 | -0.223 | 0.252 | 0.981 | -0.226 | 0.246 | 0.972 | 0.027 | 0.893 | 0.947 |
| **C2** | -0.195 | 0.319 | 0.819 | 0.007 | 0.972 | 0.997 | -0.171 | 0.382 | 0.730 | 0.169 | 0.389 | 0.981 | -0.158 | 0.421 | 0.972 | -0.045 | 0.820 | 0.947 |
| **P1** | -0.204 | 0.296 | 0.819 | -0.048 | 0.810 | 0.990 | 0.271 | 0.163 | 0.606 | 0.031 | 0.877 | 0.981 | -0.190 | 0.330 | 0.972 | 0.024 | 0.904 | 0.947 |
| **P2** | 0.043 | 0.827 | 0.922 | -0.249 | 0.201 | 0.990 | 0.205 | 0.293 | 0.704 | 0.272 | 0.161 | 0.981 | 0.120 | 0.542 | 0.972 | -0.259 | 0.183 | 0.947 |
| **AF3** | 0.138 | 0.481 | 0.819 | 0.121 | 0.538 | 0.990 | -0.183 | 0.350 | 0.730 | 0.049 | 0.803 | 0.981 | -0.125 | 0.524 | 0.972 | -0.048 | 0.807 | 0.947 |
| **AF4** | 0.111 | 0.574 | 0.902 | 0.001 | 0.997 | 0.997 | -0.274 | 0.158 | 0.606 | -0.119 | 0.546 | 0.981 | -0.066 | 0.739 | 0.972 | 0.083 | 0.675 | 0.947 |
| **FC3** | 0.305 | 0.114 | 0.819 | -0.084 | 0.669 | 0.990 | -0.080 | 0.685 | 0.847 | 0.414 | 0.029 | 0.918 | 0.136 | 0.489 | 0.972 | -0.072 | 0.716 | 0.947 |
| **FC4** | 0.225 | 0.249 | 0.819 | 0.094 | 0.633 | 0.990 | -0.219 | 0.261 | 0.704 | 0.141 | 0.472 | 0.981 | -0.118 | 0.549 | 0.972 | -0.136 | 0.488 | 0.947 |
| **CP3** | 0.212 | 0.278 | 0.819 | -0.089 | 0.653 | 0.990 | 0.026 | 0.897 | 0.940 | -0.007 | 0.974 | 0.981 | -0.303 | 0.117 | 0.972 | 0.172 | 0.380 | 0.947 |
| **CP4** | 0.143 | 0.465 | 0.819 | 0.219 | 0.262 | 0.990 | 0.028 | 0.888 | 0.940 | -0.039 | 0.842 | 0.981 | -0.078 | 0.693 | 0.972 | -0.370 | 0.053 | 0.947 |
| **PO3** | -0.374 | 0.051 | 0.746 | -0.234 | 0.229 | 0.990 | 0.149 | 0.446 | 0.785 | -0.162 | 0.408 | 0.981 | -0.169 | 0.389 | 0.972 | 0.239 | 0.219 | 0.947 |
| **PO4** | -0.137 | 0.484 | 0.819 | -0.185 | 0.344 | 0.990 | 0.238 | 0.223 | 0.704 | -0.127 | 0.518 | 0.981 | -0.227 | 0.244 | 0.972 | -0.184 | 0.346 | 0.947 |
| **F5** | 0.078 | 0.691 | 0.906 | 0.211 | 0.280 | 0.990 | 0.050 | 0.801 | 0.932 | 0.014 | 0.943 | 0.981 | -0.115 | 0.557 | 0.972 | -0.112 | 0.568 | 0.947 |
| **F6** | 0.227 | 0.245 | 0.819 | 0.316 | 0.102 | 0.990 | -0.462 | 0.014 | 0.311 | -0.167 | 0.394 | 0.981 | -0.275 | 0.156 | 0.972 | 0.338 | 0.079 | 0.947 |
| **C5** | -0.016 | 0.935 | 0.935 | 0.308 | 0.111 | 0.990 | 0.093 | 0.636 | 0.847 | 0.234 | 0.229 | 0.981 | -0.178 | 0.364 | 0.972 | -0.219 | 0.262 | 0.947 |
| **C6** | 0.024 | 0.904 | 0.935 | 0.053 | 0.790 | 0.990 | -0.275 | 0.156 | 0.606 | -0.023 | 0.908 | 0.981 | 0.261 | 0.180 | 0.972 | 0.018 | 0.928 | 0.949 |
| **P5** | -0.493 | 0.008 | 0.371 | 0.172 | 0.379 | 0.990 | -0.324 | 0.093 | 0.606 | -0.083 | 0.675 | 0.981 | -0.007 | 0.972 | 0.972 | 0.133 | 0.498 | 0.947 |
| **P6** | -0.086 | 0.663 | 0.906 | -0.196 | 0.316 | 0.990 | 0.214 | 0.273 | 0.704 | 0.261 | 0.180 | 0.981 | 0.074 | 0.708 | 0.972 | -0.142 | 0.468 | 0.947 |
| **Fpz** | 0.137 | 0.484 | 0.819 | -0.012 | 0.952 | 0.997 | 0.095 | 0.631 | 0.847 | -0.033 | 0.868 | 0.981 | -0.074 | 0.708 | 0.972 | 0.135 | 0.493 | 0.947 |
| **CPz** | -0.137 | 0.484 | 0.819 | 0.252 | 0.195 | 0.990 | -0.087 | 0.659 | 0.847 | 0.065 | 0.743 | 0.981 | -0.050 | 0.801 | 0.972 | -0.070 | 0.722 | 0.947 |
| **POz** | -0.041 | 0.838 | 0.922 | -0.152 | 0.438 | 0.990 | 0.183 | 0.349 | 0.730 | 0.026 | 0.895 | 0.981 | -0.058 | 0.769 | 0.972 | 0.193 | 0.323 | 0.947 |
| **FCz** | -0.378 | 0.048 | 0.746 | -0.199 | 0.308 | 0.990 | -0.121 | 0.538 | 0.846 | -0.061 | 0.758 | 0.981 | -0.160 | 0.413 | 0.972 | 0.148 | 0.451 | 0.947 |
|  | **HCT** | | | | | | | | | | | | | | | | | |
|  | **Beta** | | | | | | | | | | | | | | | | | |
|  | **ΔHEP and Δpower** | | | | | | **ΔHEP and ΔITC** | | | | | | **Δpower and ΔITC** | | | | | |
|  | **Inhalation** | | | **Exhalation** | | | **Inhalation** | | | **Exhalation** | | | **Inhalation** | | | **Exhalation** | | |
| **Channel** | **rho** | **p** | **p_FDR_** | **rho** | **p** | **p_FDR_** | **rho** | **p** | **p_FDR_** | **rho** | **p** | **p_FDR_** | **rho** | **p** | **p_FDR_** | **rho** | **p** | **p_FDR_** |
| **Fp1** | 0.085 | 0.665 | 0.900 | 0.275 | 0.156 | 0.919 | -0.020 | 0.921 | 0.965 | 0.244 | 0.210 | 0.862 | 0.095 | 0.631 | 0.946 | 0.011 | 0.954 | 0.992 |
| **Fp2** | 0.289 | 0.136 | 0.686 | 0.130 | 0.509 | 0.954 | 0.012 | 0.952 | 0.974 | -0.193 | 0.323 | 0.888 | -0.102 | 0.605 | 0.946 | 0.045 | 0.818 | 0.992 |
| **F3** | 0.020 | 0.919 | 0.986 | -0.068 | 0.729 | 0.954 | -0.053 | 0.788 | 0.913 | -0.143 | 0.465 | 0.957 | 0.084 | 0.669 | 0.946 | 0.091 | 0.644 | 0.992 |
| **F4** | 0.069 | 0.727 | 0.900 | -0.054 | 0.784 | 0.954 | 0.093 | 0.636 | 0.893 | 0.244 | 0.211 | 0.862 | 0.016 | 0.937 | 0.946 | -0.205 | 0.295 | 0.992 |
| **C3** | 0.197 | 0.313 | 0.708 | -0.174 | 0.376 | 0.954 | -0.119 | 0.546 | 0.893 | 0.060 | 0.762 | 0.957 | -0.085 | 0.667 | 0.946 | -0.102 | 0.603 | 0.992 |
| **C4** | 0.255 | 0.191 | 0.686 | -0.238 | 0.222 | 0.954 | 0.105 | 0.595 | 0.893 | 0.037 | 0.851 | 0.957 | 0.094 | 0.633 | 0.946 | -0.157 | 0.423 | 0.992 |
| **P3** | -0.346 | 0.072 | 0.686 | -0.043 | 0.829 | 0.954 | -0.171 | 0.383 | 0.893 | 0.219 | 0.261 | 0.862 | 0.126 | 0.522 | 0.946 | 0.384 | 0.044 | 0.992 |
| **P4** | -0.366 | 0.056 | 0.686 | 0.028 | 0.886 | 0.954 | -0.078 | 0.691 | 0.893 | 0.292 | 0.132 | 0.862 | 0.140 | 0.477 | 0.946 | 0.251 | 0.197 | 0.992 |
| **Fz** | 0.114 | 0.561 | 0.893 | -0.398 | 0.037 | 0.919 | 0.337 | 0.080 | 0.893 | 0.285 | 0.141 | 0.862 | 0.059 | 0.767 | 0.946 | -0.349 | 0.070 | 0.992 |
| **Cz** | 0.135 | 0.493 | 0.883 | 0.065 | 0.743 | 0.954 | 0.079 | 0.687 | 0.893 | -0.213 | 0.274 | 0.862 | 0.016 | 0.935 | 0.946 | 0.030 | 0.879 | 0.992 |
| **Pz** | -0.072 | 0.716 | 0.900 | -0.271 | 0.162 | 0.919 | 0.270 | 0.164 | 0.893 | -0.276 | 0.154 | 0.862 | 0.123 | 0.531 | 0.946 | 0.237 | 0.224 | 0.992 |
| **AFz** | 0.433 | 0.022 | 0.490 | -0.142 | 0.470 | 0.954 | 0.072 | 0.716 | 0.893 | 0.133 | 0.498 | 0.957 | -0.062 | 0.754 | 0.946 | -0.090 | 0.649 | 0.992 |
| **FC1** | 0.015 | 0.941 | 0.986 | -0.015 | 0.941 | 0.957 | 0.143 | 0.467 | 0.893 | 0.079 | 0.687 | 0.957 | -0.151 | 0.443 | 0.946 | -0.063 | 0.748 | 0.992 |
| **FC2** | -0.213 | 0.274 | 0.686 | 0.027 | 0.893 | 0.954 | 0.152 | 0.440 | 0.893 | 0.418 | 0.028 | 0.862 | 0.433 | 0.022 | 0.490 | 0.038 | 0.847 | 0.992 |
| **CP1** | 0.058 | 0.769 | 0.900 | 0.140 | 0.475 | 0.954 | -0.240 | 0.218 | 0.893 | 0.058 | 0.769 | 0.957 | -0.215 | 0.270 | 0.946 | 0.164 | 0.404 | 0.992 |
| **CP2** | 0.004 | 0.986 | 0.986 | -0.028 | 0.888 | 0.954 | -0.169 | 0.388 | 0.893 | -0.054 | 0.784 | 0.957 | -0.111 | 0.572 | 0.946 | 0.143 | 0.465 | 0.992 |
| **FC5** | -0.492 | 0.009 | 0.378 | 0.198 | 0.312 | 0.954 | 0.088 | 0.657 | 0.893 | 0.070 | 0.722 | 0.957 | -0.240 | 0.218 | 0.946 | 0.002 | 0.992 | 0.992 |
| **FC6** | -0.122 | 0.536 | 0.893 | 0.176 | 0.368 | 0.954 | -0.113 | 0.566 | 0.893 | -0.145 | 0.460 | 0.957 | 0.277 | 0.153 | 0.946 | -0.118 | 0.549 | 0.992 |
| **CP5** | -0.044 | 0.825 | 0.931 | -0.278 | 0.152 | 0.919 | -0.111 | 0.574 | 0.893 | -0.297 | 0.125 | 0.862 | -0.056 | 0.777 | 0.946 | -0.032 | 0.871 | 0.992 |
| **CP6** | -0.175 | 0.373 | 0.781 | 0.140 | 0.477 | 0.954 | -0.117 | 0.551 | 0.893 | -0.039 | 0.842 | 0.957 | -0.201 | 0.304 | 0.946 | -0.263 | 0.175 | 0.992 |
| **F1** | -0.008 | 0.970 | 0.986 | -0.188 | 0.337 | 0.954 | -0.004 | 0.986 | 0.986 | 0.024 | 0.904 | 0.957 | -0.051 | 0.795 | 0.946 | -0.215 | 0.270 | 0.992 |
| **F2** | 0.136 | 0.489 | 0.883 | -0.156 | 0.426 | 0.954 | 0.068 | 0.731 | 0.893 | 0.274 | 0.158 | 0.862 | -0.018 | 0.930 | 0.946 | -0.162 | 0.408 | 0.992 |
| **C1** | 0.211 | 0.280 | 0.686 | 0.159 | 0.417 | 0.954 | -0.209 | 0.284 | 0.893 | -0.240 | 0.218 | 0.862 | -0.148 | 0.450 | 0.946 | 0.059 | 0.767 | 0.992 |
| **C2** | -0.274 | 0.158 | 0.686 | -0.011 | 0.957 | 0.957 | -0.020 | 0.919 | 0.965 | -0.014 | 0.943 | 0.957 | 0.101 | 0.607 | 0.946 | -0.051 | 0.795 | 0.992 |
| **P1** | -0.267 | 0.169 | 0.686 | -0.092 | 0.640 | 0.954 | 0.188 | 0.337 | 0.893 | -0.065 | 0.743 | 0.957 | 0.014 | 0.946 | 0.946 | 0.269 | 0.166 | 0.992 |
| **P2** | 0.005 | 0.981 | 0.986 | 0.024 | 0.906 | 0.954 | 0.393 | 0.039 | 0.893 | -0.219 | 0.262 | 0.862 | 0.215 | 0.270 | 0.946 | 0.093 | 0.638 | 0.992 |
| **AF3** | 0.274 | 0.158 | 0.686 | -0.022 | 0.910 | 0.954 | 0.055 | 0.782 | 0.913 | 0.054 | 0.784 | 0.957 | -0.025 | 0.901 | 0.946 | -0.008 | 0.968 | 0.992 |
| **AF4** | 0.303 | 0.118 | 0.686 | 0.104 | 0.597 | 0.954 | -0.166 | 0.397 | 0.893 | -0.061 | 0.758 | 0.957 | -0.191 | 0.329 | 0.946 | -0.102 | 0.603 | 0.992 |
| **FC3** | -0.088 | 0.657 | 0.900 | 0.270 | 0.165 | 0.919 | -0.215 | 0.272 | 0.893 | 0.084 | 0.671 | 0.957 | -0.250 | 0.198 | 0.946 | -0.140 | 0.477 | 0.992 |
| **FC4** | -0.066 | 0.737 | 0.900 | 0.191 | 0.329 | 0.954 | 0.310 | 0.109 | 0.893 | 0.257 | 0.187 | 0.862 | 0.442 | 0.019 | 0.490 | 0.205 | 0.293 | 0.992 |
| **CP3** | 0.221 | 0.258 | 0.686 | -0.104 | 0.597 | 0.954 | -0.325 | 0.092 | 0.893 | -0.038 | 0.849 | 0.957 | -0.314 | 0.104 | 0.946 | -0.010 | 0.961 | 0.992 |
| **CP4** | -0.059 | 0.765 | 0.900 | 0.285 | 0.141 | 0.919 | 0.120 | 0.542 | 0.893 | -0.011 | 0.957 | 0.957 | -0.322 | 0.095 | 0.946 | 0.060 | 0.762 | 0.992 |
| **PO3** | -0.226 | 0.246 | 0.686 | -0.102 | 0.603 | 0.954 | -0.020 | 0.921 | 0.965 | -0.170 | 0.386 | 0.957 | -0.263 | 0.175 | 0.946 | 0.085 | 0.667 | 0.992 |
| **PO4** | -0.262 | 0.177 | 0.686 | -0.067 | 0.735 | 0.954 | 0.173 | 0.377 | 0.893 | -0.201 | 0.304 | 0.888 | 0.036 | 0.855 | 0.946 | 0.069 | 0.727 | 0.992 |
| **F5** | -0.096 | 0.625 | 0.900 | 0.188 | 0.336 | 0.954 | -0.374 | 0.050 | 0.893 | -0.100 | 0.611 | 0.957 | -0.064 | 0.746 | 0.946 | -0.177 | 0.365 | 0.992 |
| **F6** | 0.132 | 0.502 | 0.883 | 0.051 | 0.795 | 0.954 | -0.204 | 0.297 | 0.893 | 0.143 | 0.465 | 0.957 | -0.078 | 0.691 | 0.946 | 0.267 | 0.170 | 0.992 |
| **C5** | -0.156 | 0.426 | 0.853 | 0.042 | 0.833 | 0.954 | 0.196 | 0.315 | 0.893 | -0.091 | 0.644 | 0.957 | -0.033 | 0.866 | 0.946 | -0.126 | 0.520 | 0.992 |
| **C6** | 0.327 | 0.089 | 0.686 | 0.173 | 0.377 | 0.954 | -0.077 | 0.695 | 0.893 | -0.032 | 0.871 | 0.957 | -0.078 | 0.693 | 0.946 | 0.058 | 0.769 | 0.992 |
| **P5** | -0.194 | 0.322 | 0.708 | -0.041 | 0.836 | 0.954 | -0.169 | 0.388 | 0.893 | -0.148 | 0.450 | 0.957 | 0.039 | 0.844 | 0.946 | 0.067 | 0.733 | 0.992 |
| **P6** | -0.088 | 0.657 | 0.900 | 0.074 | 0.708 | 0.954 | -0.250 | 0.198 | 0.893 | -0.017 | 0.932 | 0.957 | 0.179 | 0.361 | 0.946 | 0.147 | 0.455 | 0.992 |
| **Fpz** | 0.215 | 0.270 | 0.686 | 0.314 | 0.104 | 0.919 | -0.036 | 0.855 | 0.965 | -0.034 | 0.864 | 0.957 | -0.030 | 0.879 | 0.946 | 0.037 | 0.853 | 0.992 |
| **CPz** | 0.234 | 0.230 | 0.686 | -0.268 | 0.167 | 0.919 | -0.076 | 0.700 | 0.893 | -0.273 | 0.159 | 0.862 | 0.183 | 0.349 | 0.946 | 0.090 | 0.649 | 0.992 |
| **POz** | -0.112 | 0.568 | 0.893 | 0.032 | 0.871 | 0.954 | 0.106 | 0.589 | 0.893 | -0.071 | 0.720 | 0.957 | -0.158 | 0.420 | 0.946 | -0.004 | 0.983 | 0.992 |
| **FCz** | -0.056 | 0.777 | 0.900 | 0.176 | 0.370 | 0.954 | 0.250 | 0.198 | 0.893 | 0.024 | 0.906 | 0.957 | 0.055 | 0.780 | 0.946 | 0.021 | 0.915 | 0.992 |
|  | **HCT** | | | | | | | | | | | | | | | | | |
|  | **Gamma** | | | | | | | | | | | | | | | | | |
|  | **ΔHEP and Δpower** | | | | | | **ΔHEP and ΔITC** | | | | | | **Δpower and ΔITC** | | | | | |
|  | **Inhalation** | | | **Exhalation** | | | **Inhalation** | | | **Exhalation** | | | **Inhalation** | | | **Exhalation** | | |
| **Channel** | **rho** | **p** | **p_FDR_** | **rho** | **p** | **p_FDR_** | **rho** | **p** | **p_FDR_** | **rho** | **p** | **p_FDR_** | **rho** | **p** | **p_FDR_** | **rho** | **p** | **p_FDR_** |
| **Fp1** | -0.147 | 0.455 | 0.728 | -0.165 | 0.399 | 0.975 | 0.042 | 0.831 | 0.989 | -0.007 | 0.972 | 0.995 | -0.039 | 0.842 | 0.946 | 0.186 | 0.341 | 0.806 |
| **Fp2** | -0.004 | 0.986 | 0.986 | 0.327 | 0.089 | 0.835 | 0.409 | 0.031 | 0.232 | -0.165 | 0.399 | 0.995 | -0.187 | 0.339 | 0.850 | -0.101 | 0.609 | 0.806 |
| **F3** | 0.200 | 0.307 | 0.728 | -0.181 | 0.355 | 0.918 | 0.000 | 1.000 | 1.000 | 0.207 | 0.288 | 0.845 | -0.035 | 0.860 | 0.946 | 0.128 | 0.514 | 0.806 |
| **F4** | 0.031 | 0.877 | 0.898 | 0.266 | 0.171 | 0.835 | -0.057 | 0.771 | 0.989 | -0.251 | 0.197 | 0.830 | 0.364 | 0.058 | 0.530 | -0.132 | 0.502 | 0.806 |
| **C3** | -0.418 | 0.028 | 0.615 | -0.014 | 0.943 | 0.977 | -0.409 | 0.032 | 0.232 | -0.227 | 0.245 | 0.830 | 0.077 | 0.697 | 0.946 | -0.119 | 0.544 | 0.806 |
| **C4** | -0.195 | 0.318 | 0.728 | 0.003 | 0.988 | 0.988 | 0.450 | 0.017 | 0.232 | -0.374 | 0.051 | 0.677 | -0.070 | 0.722 | 0.946 | -0.111 | 0.572 | 0.806 |
| **P3** | -0.281 | 0.147 | 0.728 | -0.129 | 0.511 | 0.977 | -0.346 | 0.071 | 0.350 | 0.109 | 0.578 | 0.995 | 0.323 | 0.094 | 0.530 | -0.014 | 0.946 | 0.991 |
| **P4** | 0.147 | 0.455 | 0.728 | 0.016 | 0.937 | 0.977 | 0.197 | 0.313 | 0.849 | 0.374 | 0.050 | 0.677 | 0.286 | 0.140 | 0.566 | -0.112 | 0.568 | 0.806 |
| **Fz** | 0.188 | 0.336 | 0.728 | 0.244 | 0.211 | 0.884 | 0.027 | 0.890 | 0.993 | 0.086 | 0.663 | 0.995 | 0.043 | 0.827 | 0.946 | 0.014 | 0.946 | 0.991 |
| **Cz** | -0.164 | 0.404 | 0.728 | -0.031 | 0.877 | 0.977 | 0.080 | 0.683 | 0.989 | -0.050 | 0.799 | 0.995 | 0.037 | 0.851 | 0.946 | -0.097 | 0.623 | 0.806 |
| **Pz** | -0.129 | 0.513 | 0.776 | 0.030 | 0.882 | 0.977 | -0.084 | 0.671 | 0.989 | 0.102 | 0.603 | 0.995 | 0.097 | 0.623 | 0.946 | -0.170 | 0.385 | 0.806 |
| **AFz** | 0.084 | 0.669 | 0.815 | 0.029 | 0.884 | 0.977 | 0.217 | 0.265 | 0.779 | -0.122 | 0.535 | 0.995 | 0.102 | 0.605 | 0.946 | 0.144 | 0.463 | 0.806 |
| **FC1** | 0.168 | 0.391 | 0.728 | 0.348 | 0.071 | 0.835 | 0.247 | 0.205 | 0.750 | 0.000 | 1.000 | 1.000 | -0.028 | 0.886 | 0.951 | -0.217 | 0.265 | 0.806 |
| **FC2** | -0.048 | 0.810 | 0.864 | 0.216 | 0.268 | 0.884 | 0.289 | 0.136 | 0.543 | 0.195 | 0.319 | 0.877 | 0.051 | 0.797 | 0.946 | 0.323 | 0.093 | 0.806 |
| **CP1** | -0.044 | 0.825 | 0.864 | -0.040 | 0.840 | 0.977 | -0.101 | 0.607 | 0.989 | 0.019 | 0.926 | 0.995 | 0.328 | 0.089 | 0.530 | -0.240 | 0.217 | 0.806 |
| **CP2** | -0.200 | 0.307 | 0.728 | 0.296 | 0.126 | 0.835 | 0.048 | 0.810 | 0.989 | -0.134 | 0.496 | 0.995 | -0.044 | 0.823 | 0.946 | 0.112 | 0.570 | 0.806 |
| **FC5** | -0.241 | 0.216 | 0.728 | 0.088 | 0.655 | 0.977 | -0.026 | 0.897 | 0.993 | -0.085 | 0.667 | 0.995 | -0.268 | 0.167 | 0.566 | 0.132 | 0.500 | 0.806 |
| **FC6** | 0.046 | 0.816 | 0.864 | 0.022 | 0.912 | 0.977 | -0.049 | 0.805 | 0.989 | -0.133 | 0.498 | 0.995 | -0.275 | 0.156 | 0.566 | -0.149 | 0.446 | 0.806 |
| **CP5** | -0.151 | 0.443 | 0.728 | 0.125 | 0.524 | 0.977 | -0.219 | 0.262 | 0.779 | -0.325 | 0.092 | 0.677 | 0.402 | 0.035 | 0.530 | 0.053 | 0.790 | 0.940 |
| **CP6** | -0.201 | 0.304 | 0.728 | -0.083 | 0.673 | 0.977 | -0.148 | 0.450 | 0.899 | -0.218 | 0.264 | 0.830 | -0.063 | 0.748 | 0.946 | -0.081 | 0.681 | 0.856 |
| **F1** | 0.113 | 0.566 | 0.788 | -0.189 | 0.334 | 0.918 | -0.020 | 0.921 | 0.993 | 0.353 | 0.066 | 0.677 | 0.245 | 0.208 | 0.609 | 0.303 | 0.117 | 0.806 |
| **F2** | 0.144 | 0.463 | 0.728 | 0.206 | 0.291 | 0.884 | 0.007 | 0.974 | 0.997 | 0.059 | 0.767 | 0.995 | 0.095 | 0.629 | 0.946 | -0.140 | 0.475 | 0.806 |
| **C1** | -0.249 | 0.201 | 0.728 | 0.317 | 0.101 | 0.835 | -0.128 | 0.514 | 0.984 | -0.290 | 0.135 | 0.742 | 0.325 | 0.092 | 0.530 | -0.170 | 0.386 | 0.806 |
| **C2** | -0.064 | 0.746 | 0.863 | -0.093 | 0.638 | 0.977 | -0.400 | 0.036 | 0.232 | 0.015 | 0.939 | 0.995 | 0.106 | 0.591 | 0.946 | 0.003 | 0.988 | 0.994 |
| **P1** | -0.463 | 0.014 | 0.615 | -0.011 | 0.954 | 0.977 | -0.069 | 0.727 | 0.989 | -0.014 | 0.946 | 0.995 | 0.046 | 0.816 | 0.946 | -0.198 | 0.311 | 0.806 |
| **P2** | -0.193 | 0.323 | 0.728 | 0.288 | 0.137 | 0.835 | 0.160 | 0.413 | 0.899 | 0.100 | 0.613 | 0.995 | 0.021 | 0.917 | 0.961 | 0.137 | 0.484 | 0.806 |
| **AF3** | -0.155 | 0.428 | 0.728 | 0.114 | 0.563 | 0.977 | -0.062 | 0.752 | 0.989 | 0.258 | 0.184 | 0.830 | -0.174 | 0.376 | 0.850 | 0.140 | 0.475 | 0.806 |
| **AF4** | 0.168 | 0.391 | 0.728 | 0.078 | 0.693 | 0.977 | 0.190 | 0.330 | 0.849 | -0.300 | 0.121 | 0.742 | -0.058 | 0.769 | 0.946 | -0.424 | 0.026 | 0.806 |
| **FC3** | -0.088 | 0.657 | 0.815 | 0.099 | 0.617 | 0.977 | -0.011 | 0.957 | 0.997 | 0.089 | 0.651 | 0.995 | -0.134 | 0.495 | 0.946 | -0.076 | 0.702 | 0.857 |
| **FC4** | -0.317 | 0.100 | 0.728 | 0.445 | 0.019 | 0.805 | 0.184 | 0.347 | 0.849 | 0.219 | 0.262 | 0.830 | -0.101 | 0.607 | 0.946 | 0.261 | 0.179 | 0.806 |
| **CP3** | -0.059 | 0.765 | 0.863 | 0.202 | 0.301 | 0.884 | -0.438 | 0.021 | 0.232 | 0.333 | 0.083 | 0.677 | 0.179 | 0.361 | 0.850 | -0.022 | 0.912 | 0.991 |
| **CP4** | -0.170 | 0.385 | 0.728 | -0.060 | 0.762 | 0.977 | 0.370 | 0.053 | 0.294 | -0.401 | 0.036 | 0.677 | -0.172 | 0.380 | 0.850 | 0.044 | 0.825 | 0.955 |
| **PO3** | -0.155 | 0.428 | 0.728 | -0.093 | 0.636 | 0.977 | -0.558 | 0.002 | 0.106 | -0.035 | 0.860 | 0.995 | 0.321 | 0.096 | 0.530 | 0.144 | 0.462 | 0.806 |
| **PO4** | -0.268 | 0.167 | 0.728 | -0.270 | 0.165 | 0.835 | 0.227 | 0.244 | 0.779 | 0.060 | 0.760 | 0.995 | 0.272 | 0.161 | 0.566 | -0.223 | 0.252 | 0.806 |
| **F5** | 0.080 | 0.685 | 0.815 | 0.031 | 0.875 | 0.977 | -0.019 | 0.926 | 0.993 | -0.010 | 0.959 | 0.995 | -0.353 | 0.066 | 0.530 | -0.125 | 0.524 | 0.806 |
| **F6** | -0.201 | 0.303 | 0.728 | 0.398 | 0.037 | 0.805 | -0.114 | 0.561 | 0.987 | 0.053 | 0.790 | 0.995 | 0.157 | 0.423 | 0.886 | -0.105 | 0.595 | 0.806 |
| **C5** | -0.159 | 0.417 | 0.728 | 0.117 | 0.553 | 0.977 | -0.088 | 0.657 | 0.989 | 0.075 | 0.704 | 0.995 | -0.080 | 0.683 | 0.946 | 0.287 | 0.139 | 0.806 |
| **C6** | -0.106 | 0.589 | 0.788 | 0.107 | 0.585 | 0.977 | 0.049 | 0.805 | 0.989 | 0.138 | 0.481 | 0.995 | -0.170 | 0.386 | 0.850 | -0.259 | 0.182 | 0.806 |
| **P5** | 0.231 | 0.236 | 0.728 | -0.060 | 0.760 | 0.977 | -0.398 | 0.037 | 0.232 | -0.228 | 0.243 | 0.830 | 0.362 | 0.059 | 0.530 | 0.159 | 0.417 | 0.806 |
| **P6** | -0.083 | 0.675 | 0.815 | -0.095 | 0.631 | 0.977 | 0.114 | 0.561 | 0.987 | -0.056 | 0.777 | 0.995 | -0.009 | 0.966 | 0.966 | -0.028 | 0.886 | 0.991 |
| **Fpz** | 0.176 | 0.370 | 0.728 | -0.215 | 0.270 | 0.884 | 0.311 | 0.107 | 0.470 | 0.021 | 0.915 | 0.995 | 0.303 | 0.118 | 0.566 | 0.002 | 0.994 | 0.994 |
| **CPz** | 0.124 | 0.529 | 0.776 | -0.231 | 0.236 | 0.884 | 0.058 | 0.769 | 0.989 | 0.027 | 0.890 | 0.995 | 0.253 | 0.193 | 0.608 | 0.258 | 0.184 | 0.806 |
| **POz** | -0.274 | 0.158 | 0.728 | -0.119 | 0.546 | 0.977 | 0.155 | 0.430 | 0.899 | 0.056 | 0.775 | 0.995 | -0.082 | 0.679 | 0.946 | -0.124 | 0.529 | 0.806 |
| **FCz** | -0.106 | 0.591 | 0.788 | -0.049 | 0.803 | 0.977 | 0.170 | 0.386 | 0.895 | 0.058 | 0.769 | 0.995 | -0.014 | 0.946 | 0.966 | 0.241 | 0.215 | 0.806 |
|  | **C-TCT** | | | | | | | | | | | | | | | | | |
|  | **Theta** | | | | | | | | | | | | | | | | | |
|  | **ΔHEP and Δpower** | | | | | | **ΔHEP and ΔITC** | | | | | | **Δpower and ΔITC** | | | | | |
|  | **Inhalation** | | | **Exhalation** | | | **Inhalation** | | | **Exhalation** | | | **Inhalation** | | | **Exhalation** | | |
| **Channel** | **rho** | **p** | **p_FDR_** | **rho** | **p** | **p_FDR_** | **rho** | **p** | **p_FDR_** | **rho** | **p** | **p_FDR_** | **rho** | **p** | **p_FDR_** | **rho** | **p** | **p_FDR_** |
| **Fp1** | 0.299 | 0.122 | 0.714 | -0.367 | 0.055 | 0.692 | 0.104 | 0.597 | 0.938 | 0.085 | 0.665 | 0.869 | 0.210 | 0.282 | 0.631 | -0.357 | 0.063 | 0.693 |
| **Fp2** | 0.271 | 0.162 | 0.714 | -0.256 | 0.189 | 0.843 | -0.076 | 0.700 | 0.938 | -0.073 | 0.712 | 0.869 | 0.193 | 0.324 | 0.631 | -0.099 | 0.617 | 0.825 |
| **F3** | 0.213 | 0.274 | 0.714 | -0.176 | 0.370 | 0.878 | 0.051 | 0.795 | 0.938 | -0.111 | 0.572 | 0.812 | 0.026 | 0.895 | 0.942 | -0.147 | 0.455 | 0.825 |
| **F4** | -0.086 | 0.661 | 0.969 | 0.068 | 0.729 | 0.878 | 0.071 | 0.718 | 0.938 | -0.544 | 0.003 | 0.140 | 0.369 | 0.054 | 0.397 | 0.234 | 0.229 | 0.825 |
| **C3** | 0.281 | 0.147 | 0.714 | 0.063 | 0.750 | 0.878 | -0.279 | 0.150 | 0.938 | -0.056 | 0.775 | 0.879 | 0.195 | 0.318 | 0.631 | -0.048 | 0.810 | 0.891 |
| **C4** | -0.168 | 0.391 | 0.757 | -0.106 | 0.589 | 0.878 | 0.151 | 0.443 | 0.938 | -0.196 | 0.315 | 0.812 | 0.114 | 0.563 | 0.773 | -0.117 | 0.553 | 0.825 |
| **P3** | -0.166 | 0.396 | 0.757 | 0.100 | 0.611 | 0.878 | -0.212 | 0.277 | 0.938 | -0.218 | 0.263 | 0.812 | 0.683 | 0.000 | 0.004 | 0.107 | 0.587 | 0.825 |
| **P4** | -0.019 | 0.926 | 0.970 | 0.155 | 0.430 | 0.878 | -0.042 | 0.833 | 0.938 | -0.167 | 0.393 | 0.812 | -0.149 | 0.448 | 0.758 | 0.089 | 0.653 | 0.825 |
| **Fz** | -0.115 | 0.557 | 0.908 | 0.219 | 0.262 | 0.878 | 0.018 | 0.930 | 0.974 | 0.018 | 0.930 | 0.930 | 0.104 | 0.597 | 0.773 | 0.025 | 0.899 | 0.942 |
| **Cz** | -0.041 | 0.836 | 0.970 | -0.090 | 0.647 | 0.878 | -0.037 | 0.853 | 0.938 | 0.153 | 0.436 | 0.812 | -0.207 | 0.289 | 0.631 | 0.430 | 0.023 | 0.486 |
| **Pz** | -0.196 | 0.315 | 0.714 | 0.346 | 0.072 | 0.692 | 0.245 | 0.208 | 0.938 | -0.070 | 0.722 | 0.869 | 0.105 | 0.595 | 0.773 | 0.112 | 0.570 | 0.825 |
| **AFz** | 0.201 | 0.304 | 0.714 | 0.152 | 0.438 | 0.878 | 0.082 | 0.677 | 0.938 | -0.134 | 0.495 | 0.812 | 0.188 | 0.337 | 0.631 | -0.257 | 0.187 | 0.825 |
| **FC1** | 0.010 | 0.961 | 0.970 | 0.091 | 0.642 | 0.878 | 0.082 | 0.679 | 0.938 | 0.201 | 0.304 | 0.812 | 0.114 | 0.561 | 0.773 | 0.159 | 0.418 | 0.825 |
| **FC2** | 0.025 | 0.901 | 0.970 | -0.314 | 0.104 | 0.692 | -0.193 | 0.323 | 0.938 | -0.114 | 0.563 | 0.812 | 0.495 | 0.008 | 0.119 | 0.183 | 0.350 | 0.825 |
| **CP1** | 0.019 | 0.926 | 0.970 | 0.070 | 0.724 | 0.878 | -0.356 | 0.064 | 0.938 | 0.135 | 0.491 | 0.812 | 0.075 | 0.704 | 0.817 | 0.055 | 0.782 | 0.887 |
| **CP2** | -0.117 | 0.551 | 0.908 | 0.254 | 0.192 | 0.843 | 0.160 | 0.413 | 0.938 | 0.449 | 0.017 | 0.384 | 0.043 | 0.827 | 0.910 | 0.009 | 0.963 | 0.963 |
| **FC5** | 0.008 | 0.968 | 0.970 | 0.182 | 0.353 | 0.878 | -0.107 | 0.585 | 0.938 | 0.173 | 0.377 | 0.812 | 0.050 | 0.801 | 0.904 | -0.103 | 0.601 | 0.825 |
| **FC6** | 0.099 | 0.617 | 0.969 | 0.034 | 0.862 | 0.891 | 0.083 | 0.673 | 0.938 | -0.084 | 0.671 | 0.869 | 0.218 | 0.263 | 0.631 | 0.306 | 0.113 | 0.825 |
| **CP5** | 0.089 | 0.653 | 0.969 | -0.081 | 0.681 | 0.878 | -0.140 | 0.475 | 0.938 | 0.153 | 0.435 | 0.812 | 0.270 | 0.164 | 0.631 | 0.102 | 0.603 | 0.825 |
| **CP6** | -0.028 | 0.886 | 0.970 | -0.102 | 0.605 | 0.878 | -0.209 | 0.284 | 0.938 | -0.336 | 0.081 | 0.812 | -0.096 | 0.625 | 0.773 | 0.088 | 0.657 | 0.825 |
| **F1** | 0.264 | 0.174 | 0.714 | 0.143 | 0.467 | 0.878 | 0.083 | 0.673 | 0.938 | -0.211 | 0.280 | 0.812 | 0.217 | 0.267 | 0.631 | -0.054 | 0.786 | 0.887 |
| **F2** | 0.240 | 0.217 | 0.714 | 0.134 | 0.495 | 0.878 | 0.113 | 0.564 | 0.938 | -0.311 | 0.108 | 0.812 | 0.267 | 0.169 | 0.631 | -0.041 | 0.838 | 0.899 |
| **C1** | -0.333 | 0.084 | 0.714 | 0.105 | 0.593 | 0.878 | 0.112 | 0.568 | 0.938 | -0.210 | 0.282 | 0.812 | -0.074 | 0.706 | 0.817 | 0.095 | 0.629 | 0.825 |
| **C2** | -0.194 | 0.322 | 0.714 | -0.054 | 0.784 | 0.878 | 0.045 | 0.820 | 0.938 | 0.158 | 0.421 | 0.812 | -0.162 | 0.408 | 0.719 | 0.161 | 0.412 | 0.825 |
| **P1** | 0.077 | 0.695 | 0.970 | 0.181 | 0.356 | 0.878 | -0.144 | 0.463 | 0.938 | 0.022 | 0.912 | 0.930 | 0.334 | 0.082 | 0.518 | 0.217 | 0.267 | 0.825 |
| **P2** | 0.140 | 0.477 | 0.840 | 0.271 | 0.162 | 0.843 | -0.063 | 0.750 | 0.938 | 0.206 | 0.291 | 0.812 | 0.229 | 0.239 | 0.631 | -0.129 | 0.511 | 0.825 |
| **AF3** | 0.233 | 0.233 | 0.714 | -0.314 | 0.104 | 0.692 | 0.233 | 0.233 | 0.938 | 0.028 | 0.886 | 0.930 | 0.246 | 0.207 | 0.631 | -0.204 | 0.296 | 0.825 |
| **AF4** | 0.333 | 0.083 | 0.714 | -0.060 | 0.760 | 0.878 | 0.075 | 0.704 | 0.938 | 0.113 | 0.566 | 0.812 | 0.238 | 0.223 | 0.631 | 0.172 | 0.379 | 0.825 |
| **FC3** | -0.022 | 0.912 | 0.970 | 0.054 | 0.784 | 0.878 | -0.011 | 0.957 | 0.979 | 0.068 | 0.731 | 0.869 | 0.001 | 0.997 | 0.997 | -0.103 | 0.599 | 0.825 |
| **FC4** | 0.207 | 0.289 | 0.714 | 0.104 | 0.597 | 0.878 | -0.189 | 0.334 | 0.938 | -0.134 | 0.496 | 0.812 | 0.263 | 0.176 | 0.631 | -0.060 | 0.762 | 0.887 |
| **CP3** | -0.059 | 0.765 | 0.970 | 0.317 | 0.101 | 0.692 | -0.252 | 0.194 | 0.938 | 0.070 | 0.722 | 0.869 | 0.297 | 0.125 | 0.631 | 0.190 | 0.332 | 0.825 |
| **CP4** | -0.147 | 0.455 | 0.834 | 0.049 | 0.803 | 0.878 | 0.083 | 0.675 | 0.938 | -0.263 | 0.176 | 0.812 | 0.209 | 0.286 | 0.631 | -0.332 | 0.085 | 0.749 |
| **PO3** | 0.248 | 0.203 | 0.714 | -0.058 | 0.769 | 0.878 | 0.026 | 0.895 | 0.960 | 0.266 | 0.171 | 0.812 | 0.433 | 0.022 | 0.194 | 0.093 | 0.636 | 0.825 |
| **PO4** | 0.065 | 0.741 | 0.970 | 0.138 | 0.481 | 0.878 | -0.158 | 0.421 | 0.938 | -0.048 | 0.807 | 0.888 | 0.124 | 0.529 | 0.773 | 0.186 | 0.343 | 0.825 |
| **F5** | 0.262 | 0.177 | 0.714 | -0.003 | 0.990 | 0.990 | -0.169 | 0.389 | 0.938 | 0.118 | 0.548 | 0.812 | -0.185 | 0.344 | 0.631 | -0.136 | 0.489 | 0.825 |
| **F6** | 0.037 | 0.851 | 0.970 | -0.101 | 0.607 | 0.878 | -0.098 | 0.619 | 0.938 | -0.222 | 0.256 | 0.812 | 0.627 | 0.000 | 0.010 | 0.456 | 0.016 | 0.486 |
| **C5** | 0.193 | 0.324 | 0.714 | 0.193 | 0.324 | 0.878 | -0.136 | 0.489 | 0.938 | 0.269 | 0.165 | 0.812 | 0.025 | 0.899 | 0.942 | 0.204 | 0.296 | 0.825 |
| **C6** | 0.256 | 0.189 | 0.714 | -0.059 | 0.767 | 0.878 | 0.121 | 0.538 | 0.938 | 0.176 | 0.370 | 0.812 | 0.118 | 0.549 | 0.773 | 0.193 | 0.324 | 0.825 |
| **P5** | -0.196 | 0.316 | 0.714 | -0.092 | 0.640 | 0.878 | 0.045 | 0.820 | 0.938 | -0.055 | 0.780 | 0.879 | 0.446 | 0.018 | 0.194 | 0.239 | 0.220 | 0.825 |
| **P6** | -0.008 | 0.970 | 0.970 | 0.032 | 0.871 | 0.891 | 0.131 | 0.505 | 0.938 | -0.379 | 0.048 | 0.699 | 0.120 | 0.542 | 0.773 | 0.016 | 0.937 | 0.959 |
| **Fpz** | 0.269 | 0.165 | 0.714 | -0.369 | 0.054 | 0.692 | 0.157 | 0.423 | 0.938 | -0.024 | 0.906 | 0.930 | 0.094 | 0.633 | 0.773 | -0.406 | 0.033 | 0.486 |
| **CPz** | -0.059 | 0.767 | 0.970 | -0.108 | 0.582 | 0.878 | 0.051 | 0.795 | 0.938 | 0.247 | 0.204 | 0.812 | -0.095 | 0.629 | 0.773 | 0.174 | 0.376 | 0.825 |
| **POz** | -0.171 | 0.382 | 0.757 | 0.309 | 0.110 | 0.692 | 0.004 | 0.983 | 0.983 | -0.151 | 0.441 | 0.812 | 0.222 | 0.256 | 0.631 | 0.133 | 0.498 | 0.825 |
| **FCz** | -0.248 | 0.202 | 0.714 | 0.045 | 0.818 | 0.878 | 0.284 | 0.144 | 0.938 | 0.132 | 0.500 | 0.812 | -0.006 | 0.977 | 0.997 | -0.080 | 0.683 | 0.835 |
|  | **C-TCT** | | | | | | | | | | | | | | | | | |
|  | **Alpha** | | | | | | | | | | | | | | | | | |
|  | **ΔHEP and Δpower** | | | | | | **ΔHEP and ΔITC** | | | | | | **Δpower and ΔITC** | | | | | |
|  | **Inhalation** | | | **Exhalation** | | | **Inhalation** | | | **Exhalation** | | | **Inhalation** | | | **Exhalation** | | |
| **Channel** | **rho** | **p** | **p_FDR_** | **rho** | **p** | **p_FDR_** | **rho** | **p** | **p_FDR_** | **rho** | **p** | **p_FDR_** | **rho** | **p** | **p_FDR_** | **rho** | **p** | **p_FDR_** |
| **Fp1** | -0.022 | 0.912 | 0.997 | -0.180 | 0.358 | 0.721 | 0.336 | 0.081 | 0.553 | -0.268 | 0.167 | 0.751 | -0.142 | 0.470 | 0.980 | 0.091 | 0.642 | 0.908 |
| **Fp2** | -0.209 | 0.284 | 0.863 | -0.323 | 0.093 | 0.685 | 0.004 | 0.983 | 0.983 | -0.221 | 0.258 | 0.790 | 0.048 | 0.807 | 0.980 | 0.118 | 0.548 | 0.908 |
| **F3** | 0.063 | 0.750 | 0.970 | 0.024 | 0.904 | 0.943 | 0.059 | 0.767 | 0.955 | -0.192 | 0.326 | 0.797 | -0.140 | 0.475 | 0.980 | 0.019 | 0.923 | 0.945 |
| **F4** | -0.079 | 0.687 | 0.953 | -0.072 | 0.716 | 0.852 | -0.033 | 0.866 | 0.955 | -0.556 | 0.002 | 0.063 | -0.014 | 0.946 | 0.991 | 0.376 | 0.049 | 0.848 |
| **C3** | 0.015 | 0.939 | 0.997 | 0.469 | 0.013 | 0.263 | -0.115 | 0.557 | 0.955 | -0.153 | 0.435 | 0.889 | 0.062 | 0.752 | 0.980 | 0.217 | 0.265 | 0.908 |
| **C4** | -0.119 | 0.544 | 0.891 | 0.165 | 0.399 | 0.721 | -0.033 | 0.868 | 0.955 | -0.057 | 0.773 | 0.958 | 0.066 | 0.739 | 0.980 | 0.166 | 0.397 | 0.908 |
| **P3** | 0.005 | 0.981 | 0.997 | 0.142 | 0.470 | 0.721 | -0.444 | 0.019 | 0.553 | 0.066 | 0.737 | 0.958 | 0.082 | 0.677 | 0.980 | -0.276 | 0.154 | 0.908 |
| **P4** | 0.458 | 0.015 | 0.568 | -0.183 | 0.350 | 0.721 | 0.010 | 0.961 | 0.983 | 0.136 | 0.488 | 0.889 | 0.033 | 0.868 | 0.980 | -0.196 | 0.316 | 0.908 |
| **Fz** | 0.189 | 0.333 | 0.863 | 0.143 | 0.465 | 0.721 | -0.076 | 0.702 | 0.955 | -0.135 | 0.491 | 0.889 | -0.054 | 0.786 | 0.980 | -0.028 | 0.886 | 0.945 |
| **Cz** | 0.053 | 0.790 | 0.984 | -0.163 | 0.405 | 0.721 | -0.079 | 0.687 | 0.955 | 0.016 | 0.935 | 0.970 | 0.092 | 0.640 | 0.980 | 0.001 | 0.997 | 0.997 |
| **Pz** | 0.186 | 0.343 | 0.863 | -0.078 | 0.691 | 0.852 | -0.079 | 0.689 | 0.955 | 0.359 | 0.062 | 0.679 | -0.004 | 0.983 | 0.999 | 0.094 | 0.634 | 0.908 |
| **AFz** | 0.049 | 0.805 | 0.984 | -0.160 | 0.413 | 0.721 | 0.381 | 0.046 | 0.553 | -0.216 | 0.269 | 0.790 | -0.115 | 0.557 | 0.980 | 0.068 | 0.731 | 0.919 |
| **FC1** | -0.007 | 0.974 | 0.997 | 0.133 | 0.498 | 0.725 | 0.236 | 0.225 | 0.743 | -0.049 | 0.805 | 0.958 | -0.158 | 0.420 | 0.980 | -0.086 | 0.661 | 0.908 |
| **FC2** | -0.155 | 0.430 | 0.863 | 0.020 | 0.921 | 0.943 | -0.103 | 0.599 | 0.955 | -0.175 | 0.373 | 0.863 | 0.085 | 0.665 | 0.980 | -0.060 | 0.760 | 0.922 |
| **CP1** | -0.169 | 0.389 | 0.863 | 0.148 | 0.450 | 0.721 | -0.250 | 0.198 | 0.743 | 0.273 | 0.159 | 0.751 | -0.014 | 0.943 | 0.991 | -0.041 | 0.838 | 0.922 |
| **CP2** | -0.017 | 0.932 | 0.997 | 0.120 | 0.542 | 0.745 | 0.241 | 0.215 | 0.743 | -0.117 | 0.553 | 0.901 | -0.079 | 0.687 | 0.980 | -0.197 | 0.313 | 0.908 |
| **FC5** | -0.108 | 0.582 | 0.891 | 0.218 | 0.264 | 0.721 | -0.267 | 0.170 | 0.743 | -0.266 | 0.171 | 0.751 | 0.436 | 0.021 | 0.311 | -0.196 | 0.315 | 0.908 |
| **FC6** | 0.317 | 0.100 | 0.863 | -0.447 | 0.018 | 0.263 | -0.044 | 0.823 | 0.955 | 0.088 | 0.657 | 0.932 | 0.040 | 0.840 | 0.980 | 0.330 | 0.087 | 0.848 |
| **CP5** | -0.300 | 0.120 | 0.863 | 0.241 | 0.216 | 0.721 | 0.049 | 0.805 | 0.955 | -0.011 | 0.957 | 0.970 | 0.001 | 0.999 | 0.999 | -0.138 | 0.482 | 0.908 |
| **CP6** | 0.423 | 0.026 | 0.568 | -0.189 | 0.333 | 0.721 | -0.390 | 0.041 | 0.553 | -0.054 | 0.784 | 0.958 | -0.084 | 0.669 | 0.980 | -0.154 | 0.431 | 0.908 |
| **F1** | 0.114 | 0.563 | 0.891 | 0.407 | 0.033 | 0.359 | 0.349 | 0.069 | 0.553 | -0.088 | 0.655 | 0.932 | -0.211 | 0.280 | 0.943 | -0.043 | 0.827 | 0.922 |
| **F2** | -0.168 | 0.391 | 0.863 | 0.233 | 0.231 | 0.721 | -0.064 | 0.746 | 0.955 | -0.059 | 0.767 | 0.958 | 0.085 | 0.667 | 0.980 | -0.111 | 0.574 | 0.908 |
| **C1** | -0.275 | 0.156 | 0.863 | -0.102 | 0.605 | 0.783 | -0.149 | 0.446 | 0.892 | -0.414 | 0.029 | 0.430 | 0.266 | 0.171 | 0.848 | 0.050 | 0.799 | 0.922 |
| **C2** | -0.149 | 0.446 | 0.863 | -0.038 | 0.849 | 0.934 | 0.012 | 0.952 | 0.983 | -0.018 | 0.930 | 0.970 | 0.111 | 0.574 | 0.980 | -0.073 | 0.710 | 0.919 |
| **P1** | 0.169 | 0.389 | 0.863 | 0.032 | 0.871 | 0.934 | -0.151 | 0.443 | 0.892 | 0.251 | 0.197 | 0.790 | 0.269 | 0.166 | 0.848 | 0.089 | 0.653 | 0.908 |
| **P2** | 0.247 | 0.205 | 0.863 | 0.033 | 0.866 | 0.934 | -0.082 | 0.679 | 0.955 | 0.239 | 0.219 | 0.790 | -0.233 | 0.231 | 0.861 | -0.197 | 0.313 | 0.908 |
| **AF3** | 0.078 | 0.693 | 0.953 | -0.385 | 0.044 | 0.385 | 0.294 | 0.128 | 0.627 | -0.194 | 0.320 | 0.797 | -0.323 | 0.093 | 0.823 | 0.097 | 0.621 | 0.908 |
| **AF4** | -0.117 | 0.553 | 0.891 | -0.167 | 0.393 | 0.721 | 0.317 | 0.101 | 0.553 | -0.269 | 0.165 | 0.751 | 0.036 | 0.857 | 0.980 | 0.361 | 0.060 | 0.848 |
| **FC3** | -0.212 | 0.277 | 0.863 | 0.226 | 0.246 | 0.721 | 0.077 | 0.697 | 0.955 | 0.099 | 0.615 | 0.932 | 0.069 | 0.727 | 0.980 | -0.154 | 0.431 | 0.908 |
| **FC4** | -0.022 | 0.912 | 0.997 | 0.250 | 0.198 | 0.721 | -0.207 | 0.289 | 0.743 | 0.136 | 0.488 | 0.889 | -0.018 | 0.930 | 0.991 | 0.388 | 0.042 | 0.848 |
| **CP3** | -0.001 | 0.997 | 0.997 | 0.105 | 0.595 | 0.783 | -0.121 | 0.538 | 0.955 | 0.123 | 0.533 | 0.901 | 0.043 | 0.827 | 0.980 | -0.021 | 0.917 | 0.945 |
| **CP4** | 0.116 | 0.555 | 0.891 | 0.047 | 0.814 | 0.934 | -0.035 | 0.860 | 0.955 | 0.152 | 0.440 | 0.889 | -0.063 | 0.750 | 0.980 | 0.261 | 0.180 | 0.908 |
| **PO3** | 0.200 | 0.305 | 0.863 | -0.129 | 0.511 | 0.725 | 0.020 | 0.921 | 0.983 | 0.293 | 0.130 | 0.751 | 0.479 | 0.011 | 0.235 | -0.174 | 0.374 | 0.908 |
| **PO4** | 0.356 | 0.064 | 0.863 | -0.140 | 0.475 | 0.721 | -0.155 | 0.430 | 0.892 | 0.131 | 0.505 | 0.889 | 0.264 | 0.173 | 0.848 | -0.161 | 0.410 | 0.908 |
| **F5** | -0.177 | 0.365 | 0.863 | 0.194 | 0.322 | 0.721 | 0.118 | 0.548 | 0.955 | -0.015 | 0.941 | 0.970 | -0.088 | 0.655 | 0.980 | -0.141 | 0.474 | 0.908 |
| **F6** | 0.242 | 0.214 | 0.863 | -0.151 | 0.443 | 0.721 | -0.201 | 0.304 | 0.743 | -0.550 | 0.003 | 0.063 | 0.271 | 0.163 | 0.848 | 0.169 | 0.389 | 0.908 |
| **C5** | -0.165 | 0.399 | 0.863 | 0.291 | 0.133 | 0.721 | -0.190 | 0.332 | 0.768 | -0.010 | 0.961 | 0.970 | 0.513 | 0.006 | 0.235 | -0.071 | 0.720 | 0.919 |
| **C6** | 0.086 | 0.661 | 0.953 | 0.000 | 1.000 | 1.000 | -0.209 | 0.286 | 0.743 | -0.065 | 0.741 | 0.958 | -0.051 | 0.795 | 0.980 | 0.321 | 0.096 | 0.848 |
| **P5** | -0.196 | 0.315 | 0.863 | 0.187 | 0.340 | 0.721 | -0.212 | 0.278 | 0.743 | 0.105 | 0.593 | 0.932 | 0.395 | 0.038 | 0.421 | -0.049 | 0.803 | 0.922 |
| **P6** | 0.265 | 0.173 | 0.863 | -0.159 | 0.418 | 0.721 | -0.205 | 0.293 | 0.743 | -0.008 | 0.970 | 0.970 | -0.203 | 0.300 | 0.943 | 0.114 | 0.561 | 0.908 |
| **Fpz** | 0.029 | 0.884 | 0.997 | -0.476 | 0.011 | 0.263 | 0.366 | 0.056 | 0.553 | -0.021 | 0.915 | 0.970 | -0.251 | 0.196 | 0.861 | -0.114 | 0.561 | 0.908 |
| **CPz** | -0.107 | 0.587 | 0.891 | -0.262 | 0.178 | 0.721 | -0.035 | 0.860 | 0.955 | 0.281 | 0.147 | 0.751 | 0.162 | 0.408 | 0.980 | -0.144 | 0.462 | 0.908 |
| **POz** | 0.148 | 0.451 | 0.863 | -0.076 | 0.702 | 0.852 | -0.100 | 0.613 | 0.955 | 0.217 | 0.265 | 0.790 | 0.232 | 0.235 | 0.861 | 0.104 | 0.597 | 0.908 |
| **FCz** | 0.067 | 0.733 | 0.970 | 0.274 | 0.158 | 0.721 | 0.317 | 0.100 | 0.553 | -0.194 | 0.320 | 0.797 | -0.043 | 0.827 | 0.980 | -0.250 | 0.198 | 0.908 |
|  | **C-TCT** | | | | | | | | | | | | | | | | | |
|  | **Beta** | | | | | | | | | | | | | | | | | |
|  | **ΔHEP and Δpower** | | | | | | **ΔHEP and ΔITC** | | | | | | **Δpower and ΔITC** | | | | | |
|  | **Inhalation** | | | **Exhalation** | | | **Inhalation** | | | **Exhalation** | | | **Inhalation** | | | **Exhalation** | | |
| **Channel** | **rho** | **p** | **p_FDR_** | **rho** | **p** | **p_FDR_** | **rho** | **p** | **p_FDR_** | **rho** | **p** | **p_FDR_** | **rho** | **p** | **p_FDR_** | **rho** | **p** | **p_FDR_** |
| **Fp1** | -0.026 | 0.895 | 0.972 | -0.164 | 0.404 | 0.740 | 0.249 | 0.201 | 0.831 | 0.155 | 0.430 | 0.920 | 0.157 | 0.425 | 0.941 | 0.047 | 0.814 | 0.967 |
| **Fp2** | 0.048 | 0.807 | 0.972 | -0.337 | 0.080 | 0.644 | 0.025 | 0.901 | 0.967 | -0.199 | 0.309 | 0.920 | -0.019 | 0.926 | 0.994 | 0.024 | 0.906 | 0.967 |
| **F3** | 0.186 | 0.343 | 0.972 | -0.246 | 0.206 | 0.703 | 0.144 | 0.463 | 0.831 | -0.051 | 0.797 | 0.965 | 0.192 | 0.326 | 0.904 | -0.053 | 0.790 | 0.967 |
| **F4** | -0.166 | 0.397 | 0.972 | -0.215 | 0.272 | 0.703 | 0.171 | 0.383 | 0.831 | 0.001 | 0.999 | 0.999 | 0.075 | 0.704 | 0.994 | -0.114 | 0.561 | 0.967 |
| **C3** | -0.192 | 0.326 | 0.972 | -0.047 | 0.814 | 0.873 | -0.116 | 0.555 | 0.831 | -0.151 | 0.441 | 0.920 | 0.312 | 0.106 | 0.904 | 0.027 | 0.893 | 0.967 |
| **C4** | -0.031 | 0.877 | 0.972 | 0.066 | 0.739 | 0.873 | -0.483 | 0.010 | 0.220 | -0.226 | 0.247 | 0.920 | -0.010 | 0.961 | 0.994 | 0.263 | 0.175 | 0.967 |
| **P3** | -0.378 | 0.048 | 0.532 | -0.285 | 0.142 | 0.703 | 0.092 | 0.640 | 0.881 | 0.259 | 0.183 | 0.920 | -0.301 | 0.120 | 0.904 | -0.074 | 0.708 | 0.967 |
| **P4** | 0.102 | 0.605 | 0.972 | -0.226 | 0.246 | 0.703 | 0.100 | 0.613 | 0.870 | 0.172 | 0.379 | 0.920 | -0.097 | 0.623 | 0.945 | -0.192 | 0.326 | 0.967 |
| **Fz** | -0.014 | 0.946 | 0.972 | 0.065 | 0.743 | 0.873 | 0.178 | 0.364 | 0.831 | -0.135 | 0.491 | 0.920 | 0.046 | 0.816 | 0.994 | 0.024 | 0.906 | 0.967 |
| **Cz** | 0.012 | 0.952 | 0.972 | -0.181 | 0.356 | 0.740 | 0.151 | 0.441 | 0.831 | 0.077 | 0.695 | 0.965 | -0.125 | 0.524 | 0.941 | -0.205 | 0.295 | 0.967 |
| **Pz** | 0.007 | 0.972 | 0.972 | -0.369 | 0.054 | 0.644 | -0.162 | 0.408 | 0.831 | 0.001 | 0.999 | 0.999 | 0.200 | 0.305 | 0.904 | -0.290 | 0.135 | 0.967 |
| **AFz** | 0.114 | 0.561 | 0.972 | 0.022 | 0.910 | 0.910 | 0.330 | 0.087 | 0.424 | -0.206 | 0.292 | 0.920 | 0.262 | 0.178 | 0.904 | 0.037 | 0.851 | 0.967 |
| **FC1** | 0.036 | 0.855 | 0.972 | -0.144 | 0.462 | 0.807 | 0.028 | 0.886 | 0.967 | -0.137 | 0.484 | 0.920 | 0.359 | 0.062 | 0.904 | -0.311 | 0.107 | 0.967 |
| **FC2** | -0.183 | 0.349 | 0.972 | -0.051 | 0.797 | 0.873 | -0.124 | 0.527 | 0.831 | -0.065 | 0.741 | 0.965 | -0.058 | 0.769 | 0.994 | 0.077 | 0.695 | 0.967 |
| **CP1** | -0.091 | 0.644 | 0.972 | -0.056 | 0.775 | 0.873 | 0.171 | 0.383 | 0.831 | 0.132 | 0.502 | 0.920 | -0.002 | 0.994 | 0.994 | -0.050 | 0.799 | 0.967 |
| **CP2** | 0.305 | 0.115 | 0.822 | -0.201 | 0.304 | 0.740 | -0.119 | 0.544 | 0.831 | -0.179 | 0.361 | 0.920 | -0.204 | 0.296 | 0.904 | 0.091 | 0.642 | 0.967 |
| **FC5** | -0.125 | 0.525 | 0.972 | 0.059 | 0.767 | 0.873 | 0.115 | 0.557 | 0.831 | 0.065 | 0.741 | 0.965 | -0.106 | 0.589 | 0.941 | -0.157 | 0.423 | 0.967 |
| **FC6** | 0.070 | 0.722 | 0.972 | -0.166 | 0.396 | 0.740 | -0.007 | 0.974 | 0.994 | -0.215 | 0.270 | 0.920 | 0.109 | 0.578 | 0.941 | -0.149 | 0.446 | 0.967 |
| **CP5** | 0.046 | 0.816 | 0.972 | -0.055 | 0.780 | 0.873 | -0.035 | 0.860 | 0.967 | 0.118 | 0.549 | 0.945 | 0.045 | 0.820 | 0.994 | 0.084 | 0.671 | 0.967 |
| **CP6** | 0.195 | 0.318 | 0.972 | -0.054 | 0.784 | 0.873 | -0.028 | 0.888 | 0.967 | -0.044 | 0.825 | 0.965 | 0.126 | 0.522 | 0.941 | -0.238 | 0.223 | 0.967 |
| **F1** | -0.013 | 0.948 | 0.972 | 0.094 | 0.634 | 0.873 | 0.113 | 0.566 | 0.831 | 0.002 | 0.994 | 0.999 | -0.039 | 0.842 | 0.994 | -0.193 | 0.323 | 0.967 |
| **F2** | -0.010 | 0.961 | 0.972 | 0.082 | 0.679 | 0.873 | -0.161 | 0.412 | 0.831 | 0.218 | 0.264 | 0.920 | -0.305 | 0.114 | 0.904 | -0.059 | 0.767 | 0.967 |
| **C1** | -0.082 | 0.679 | 0.972 | -0.294 | 0.128 | 0.703 | -0.003 | 0.988 | 0.994 | -0.239 | 0.220 | 0.920 | 0.167 | 0.394 | 0.941 | 0.200 | 0.305 | 0.967 |
| **C2** | 0.038 | 0.849 | 0.972 | -0.078 | 0.693 | 0.873 | -0.189 | 0.334 | 0.831 | -0.028 | 0.886 | 0.996 | -0.146 | 0.456 | 0.941 | 0.207 | 0.288 | 0.967 |
| **P1** | 0.045 | 0.820 | 0.972 | -0.219 | 0.261 | 0.703 | -0.002 | 0.994 | 0.994 | 0.109 | 0.580 | 0.945 | 0.104 | 0.597 | 0.941 | -0.229 | 0.239 | 0.967 |
| **P2** | 0.049 | 0.805 | 0.972 | -0.101 | 0.609 | 0.873 | 0.136 | 0.489 | 0.831 | -0.263 | 0.176 | 0.920 | -0.234 | 0.229 | 0.904 | -0.199 | 0.308 | 0.967 |
| **AF3** | 0.117 | 0.551 | 0.972 | -0.242 | 0.214 | 0.703 | 0.417 | 0.028 | 0.335 | 0.152 | 0.440 | 0.920 | 0.103 | 0.599 | 0.941 | -0.223 | 0.252 | 0.967 |
| **AF4** | 0.161 | 0.410 | 0.972 | -0.079 | 0.687 | 0.873 | 0.360 | 0.060 | 0.424 | -0.085 | 0.665 | 0.965 | -0.007 | 0.972 | 0.994 | 0.054 | 0.784 | 0.967 |
| **FC3** | 0.055 | 0.780 | 0.972 | -0.248 | 0.203 | 0.703 | 0.412 | 0.030 | 0.335 | 0.042 | 0.833 | 0.965 | -0.134 | 0.495 | 0.941 | 0.038 | 0.849 | 0.967 |
| **FC4** | -0.122 | 0.536 | 0.972 | -0.037 | 0.853 | 0.873 | -0.135 | 0.493 | 0.831 | 0.111 | 0.572 | 0.945 | 0.042 | 0.831 | 0.994 | 0.019 | 0.923 | 0.967 |
| **CP3** | 0.282 | 0.145 | 0.822 | -0.269 | 0.166 | 0.703 | 0.042 | 0.833 | 0.967 | -0.097 | 0.623 | 0.965 | 0.067 | 0.733 | 0.994 | -0.118 | 0.549 | 0.967 |
| **CP4** | -0.060 | 0.762 | 0.972 | 0.329 | 0.088 | 0.644 | -0.182 | 0.352 | 0.831 | 0.157 | 0.423 | 0.920 | -0.109 | 0.580 | 0.941 | 0.070 | 0.722 | 0.967 |
| **PO3** | -0.009 | 0.966 | 0.972 | -0.230 | 0.237 | 0.703 | 0.190 | 0.332 | 0.831 | 0.317 | 0.100 | 0.920 | -0.191 | 0.329 | 0.904 | -0.012 | 0.952 | 0.974 |
| **PO4** | 0.138 | 0.481 | 0.972 | -0.175 | 0.371 | 0.740 | -0.331 | 0.086 | 0.424 | -0.091 | 0.642 | 0.965 | -0.014 | 0.946 | 0.994 | 0.138 | 0.481 | 0.967 |
| **F5** | 0.176 | 0.368 | 0.972 | -0.140 | 0.477 | 0.807 | 0.065 | 0.741 | 0.906 | -0.194 | 0.320 | 0.920 | 0.007 | 0.972 | 0.994 | -0.124 | 0.527 | 0.967 |
| **F6** | -0.268 | 0.168 | 0.822 | -0.353 | 0.066 | 0.644 | -0.542 | 0.003 | 0.145 | -0.380 | 0.047 | 0.920 | 0.352 | 0.066 | 0.904 | 0.141 | 0.472 | 0.967 |
| **C5** | -0.086 | 0.663 | 0.972 | 0.040 | 0.840 | 0.873 | 0.081 | 0.681 | 0.906 | 0.245 | 0.209 | 0.920 | -0.203 | 0.300 | 0.904 | -0.112 | 0.570 | 0.967 |
| **C6** | 0.519 | 0.005 | 0.227 | 0.226 | 0.246 | 0.703 | -0.374 | 0.051 | 0.424 | 0.023 | 0.908 | 0.996 | -0.280 | 0.148 | 0.904 | 0.331 | 0.086 | 0.967 |
| **P5** | -0.275 | 0.157 | 0.822 | -0.063 | 0.748 | 0.873 | -0.071 | 0.718 | 0.906 | 0.336 | 0.081 | 0.920 | -0.111 | 0.574 | 0.941 | -0.155 | 0.430 | 0.967 |
| **P6** | 0.408 | 0.032 | 0.532 | -0.378 | 0.048 | 0.644 | -0.178 | 0.364 | 0.831 | 0.060 | 0.760 | 0.965 | -0.056 | 0.775 | 0.994 | -0.150 | 0.445 | 0.967 |
| **Fpz** | 0.385 | 0.044 | 0.532 | -0.473 | 0.012 | 0.515 | 0.350 | 0.069 | 0.424 | -0.053 | 0.790 | 0.965 | 0.259 | 0.183 | 0.904 | -0.025 | 0.899 | 0.967 |
| **CPz** | 0.010 | 0.961 | 0.972 | -0.173 | 0.377 | 0.740 | 0.068 | 0.729 | 0.906 | -0.165 | 0.399 | 0.920 | -0.206 | 0.291 | 0.904 | -0.072 | 0.716 | 0.967 |
| **POz** | -0.109 | 0.580 | 0.972 | -0.191 | 0.329 | 0.740 | -0.130 | 0.509 | 0.831 | -0.018 | 0.928 | 0.996 | -0.029 | 0.884 | 0.994 | 0.005 | 0.981 | 0.981 |
| **FCz** | -0.285 | 0.142 | 0.822 | 0.128 | 0.514 | 0.838 | 0.148 | 0.451 | 0.831 | -0.155 | 0.428 | 0.920 | -0.191 | 0.329 | 0.904 | 0.023 | 0.908 | 0.967 |
|  | **C-TCT** | | | | | | | | | | | | | | | | | |
|  | **Gamma** | | | | | | | | | | | | | | | | | |
|  | **ΔHEP and Δpower** | | | | | | **ΔHEP and ΔITC** | | | | | | **Δpower and ΔITC** | | | | | |
|  | **Inhalation** | | | **Exhalation** | | | **Inhalation** | | | **Exhalation** | | | **Inhalation** | | | **Exhalation** | | |
| **Channel** | **rho** | **p** | **p_FDR_** | **rho** | **p** | **p_FDR_** | **rho** | **p** | **p_FDR_** | **rho** | **p** | **p_FDR_** | **rho** | **p** | **p_FDR_** | **rho** | **p** | **p_FDR_** |
| **Fp1** | 0.056 | 0.775 | 0.992 | -0.033 | 0.868 | 0.929 | 0.093 | 0.638 | 0.977 | -0.251 | 0.196 | 0.728 | -0.085 | 0.667 | 0.973 | -0.143 | 0.467 | 0.927 |
| **Fp2** | -0.089 | 0.653 | 0.992 | 0.115 | 0.557 | 0.908 | -0.045 | 0.820 | 0.977 | -0.099 | 0.615 | 0.933 | 0.086 | 0.661 | 0.973 | -0.129 | 0.513 | 0.927 |
| **F3** | 0.209 | 0.284 | 0.992 | 0.136 | 0.489 | 0.861 | 0.246 | 0.206 | 0.905 | -0.064 | 0.746 | 0.945 | 0.262 | 0.178 | 0.973 | -0.144 | 0.463 | 0.927 |
| **F4** | 0.073 | 0.712 | 0.992 | 0.300 | 0.120 | 0.709 | -0.045 | 0.820 | 0.977 | -0.140 | 0.475 | 0.891 | -0.174 | 0.374 | 0.973 | -0.039 | 0.844 | 0.956 |
| **C3** | -0.009 | 0.966 | 0.992 | -0.250 | 0.198 | 0.765 | -0.108 | 0.582 | 0.977 | -0.515 | 0.006 | 0.125 | 0.276 | 0.155 | 0.973 | -0.024 | 0.906 | 0.956 |
| **C4** | 0.211 | 0.279 | 0.992 | 0.211 | 0.280 | 0.765 | -0.406 | 0.033 | 0.905 | -0.033 | 0.866 | 0.953 | 0.053 | 0.788 | 0.985 | -0.094 | 0.634 | 0.931 |
| **P3** | -0.004 | 0.986 | 0.992 | -0.041 | 0.838 | 0.929 | 0.042 | 0.833 | 0.977 | 0.051 | 0.795 | 0.945 | -0.152 | 0.440 | 0.973 | 0.024 | 0.906 | 0.956 |
| **P4** | 0.158 | 0.421 | 0.992 | -0.303 | 0.117 | 0.709 | -0.081 | 0.681 | 0.977 | 0.274 | 0.158 | 0.697 | -0.079 | 0.689 | 0.973 | -0.304 | 0.116 | 0.730 |
| **Fz** | 0.108 | 0.582 | 0.992 | 0.272 | 0.161 | 0.765 | -0.001 | 0.997 | 0.997 | -0.330 | 0.087 | 0.545 | -0.039 | 0.842 | 0.985 | -0.370 | 0.053 | 0.470 |
| **Cz** | 0.248 | 0.203 | 0.992 | -0.060 | 0.760 | 0.929 | -0.208 | 0.287 | 0.971 | -0.013 | 0.948 | 0.957 | -0.216 | 0.269 | 0.973 | 0.063 | 0.748 | 0.956 |
| **Pz** | -0.137 | 0.484 | 0.992 | -0.228 | 0.242 | 0.765 | -0.047 | 0.814 | 0.977 | 0.081 | 0.681 | 0.945 | 0.179 | 0.361 | 0.973 | -0.028 | 0.888 | 0.956 |
| **AFz** | 0.356 | 0.063 | 0.930 | 0.005 | 0.979 | 0.979 | -0.117 | 0.551 | 0.977 | -0.479 | 0.011 | 0.157 | -0.449 | 0.017 | 0.470 | -0.304 | 0.116 | 0.730 |
| **FC1** | 0.210 | 0.282 | 0.992 | -0.099 | 0.617 | 0.929 | -0.053 | 0.788 | 0.977 | -0.158 | 0.420 | 0.879 | 0.051 | 0.795 | 0.985 | -0.177 | 0.365 | 0.927 |
| **FC2** | -0.065 | 0.741 | 0.992 | 0.026 | 0.895 | 0.929 | -0.295 | 0.127 | 0.905 | 0.013 | 0.950 | 0.957 | 0.032 | 0.873 | 0.985 | 0.160 | 0.413 | 0.927 |
| **CP1** | -0.049 | 0.805 | 0.992 | 0.141 | 0.474 | 0.861 | 0.160 | 0.415 | 0.977 | -0.163 | 0.405 | 0.879 | -0.076 | 0.702 | 0.973 | 0.068 | 0.731 | 0.956 |
| **CP2** | 0.140 | 0.475 | 0.992 | -0.123 | 0.533 | 0.901 | 0.276 | 0.154 | 0.905 | 0.404 | 0.034 | 0.373 | 0.128 | 0.514 | 0.973 | 0.028 | 0.886 | 0.956 |
| **FC5** | -0.037 | 0.851 | 0.992 | 0.458 | 0.015 | 0.334 | 0.283 | 0.144 | 0.905 | 0.250 | 0.198 | 0.728 | 0.055 | 0.782 | 0.985 | -0.204 | 0.297 | 0.927 |
| **FC6** | 0.002 | 0.992 | 0.992 | -0.037 | 0.851 | 0.929 | -0.130 | 0.507 | 0.977 | -0.051 | 0.795 | 0.945 | -0.281 | 0.148 | 0.973 | -0.492 | 0.009 | 0.189 |
| **CP5** | -0.084 | 0.671 | 0.992 | -0.181 | 0.355 | 0.765 | -0.009 | 0.963 | 0.997 | 0.011 | 0.957 | 0.957 | -0.199 | 0.309 | 0.973 | 0.496 | 0.008 | 0.189 |
| **CP6** | -0.076 | 0.700 | 0.992 | 0.109 | 0.578 | 0.908 | 0.251 | 0.197 | 0.905 | 0.181 | 0.355 | 0.879 | -0.189 | 0.334 | 0.973 | 0.101 | 0.609 | 0.927 |
| **F1** | 0.314 | 0.104 | 0.992 | 0.230 | 0.238 | 0.765 | 0.065 | 0.743 | 0.977 | -0.169 | 0.389 | 0.879 | -0.078 | 0.691 | 0.973 | -0.184 | 0.347 | 0.927 |
| **F2** | 0.004 | 0.986 | 0.992 | 0.294 | 0.129 | 0.709 | -0.262 | 0.178 | 0.905 | 0.058 | 0.769 | 0.945 | 0.085 | 0.667 | 0.973 | -0.080 | 0.683 | 0.956 |
| **C1** | 0.281 | 0.147 | 0.992 | -0.177 | 0.365 | 0.765 | 0.106 | 0.589 | 0.977 | -0.070 | 0.724 | 0.945 | -0.100 | 0.613 | 0.973 | -0.112 | 0.568 | 0.927 |
| **C2** | 0.230 | 0.237 | 0.992 | -0.033 | 0.868 | 0.929 | -0.050 | 0.801 | 0.977 | -0.100 | 0.611 | 0.933 | 0.250 | 0.199 | 0.973 | 0.100 | 0.611 | 0.927 |
| **P1** | 0.122 | 0.535 | 0.992 | -0.153 | 0.436 | 0.834 | 0.278 | 0.152 | 0.905 | -0.026 | 0.897 | 0.957 | 0.074 | 0.708 | 0.973 | 0.126 | 0.520 | 0.927 |
| **P2** | 0.160 | 0.413 | 0.992 | -0.215 | 0.270 | 0.765 | -0.131 | 0.505 | 0.977 | 0.294 | 0.128 | 0.627 | -0.176 | 0.370 | 0.973 | -0.202 | 0.301 | 0.927 |
| **AF3** | 0.380 | 0.047 | 0.930 | -0.407 | 0.032 | 0.475 | -0.010 | 0.959 | 0.997 | 0.038 | 0.847 | 0.953 | -0.091 | 0.644 | 0.973 | -0.143 | 0.465 | 0.927 |
| **AF4** | 0.017 | 0.932 | 0.992 | 0.027 | 0.893 | 0.929 | -0.006 | 0.977 | 0.997 | 0.151 | 0.443 | 0.886 | -0.436 | 0.021 | 0.470 | -0.015 | 0.941 | 0.963 |
| **FC3** | 0.125 | 0.525 | 0.992 | -0.185 | 0.344 | 0.765 | 0.131 | 0.504 | 0.977 | -0.091 | 0.644 | 0.945 | 0.041 | 0.836 | 0.985 | 0.004 | 0.986 | 0.986 |
| **FC4** | -0.050 | 0.801 | 0.992 | 0.084 | 0.671 | 0.929 | -0.142 | 0.468 | 0.977 | -0.036 | 0.857 | 0.953 | 0.008 | 0.970 | 0.993 | 0.180 | 0.358 | 0.927 |
| **CP3** | -0.018 | 0.928 | 0.992 | -0.163 | 0.407 | 0.814 | -0.090 | 0.647 | 0.977 | -0.206 | 0.291 | 0.864 | 0.010 | 0.959 | 0.993 | 0.054 | 0.786 | 0.956 |
| **CP4** | -0.141 | 0.472 | 0.992 | 0.071 | 0.718 | 0.929 | -0.067 | 0.733 | 0.977 | 0.355 | 0.064 | 0.471 | 0.086 | 0.661 | 0.973 | 0.114 | 0.561 | 0.927 |
| **PO3** | -0.043 | 0.829 | 0.992 | -0.537 | 0.004 | 0.160 | -0.177 | 0.367 | 0.977 | 0.137 | 0.486 | 0.891 | -0.122 | 0.536 | 0.973 | -0.023 | 0.908 | 0.956 |
| **PO4** | -0.011 | 0.957 | 0.992 | -0.181 | 0.356 | 0.765 | -0.155 | 0.428 | 0.977 | 0.181 | 0.355 | 0.879 | -0.268 | 0.167 | 0.973 | 0.109 | 0.578 | 0.927 |
| **F5** | 0.005 | 0.979 | 0.992 | -0.209 | 0.284 | 0.765 | 0.221 | 0.258 | 0.947 | -0.105 | 0.593 | 0.933 | -0.132 | 0.500 | 0.973 | 0.371 | 0.053 | 0.470 |
| **F6** | -0.058 | 0.769 | 0.992 | -0.025 | 0.901 | 0.929 | -0.307 | 0.112 | 0.905 | -0.099 | 0.615 | 0.933 | -0.192 | 0.326 | 0.973 | -0.044 | 0.823 | 0.956 |
| **C5** | -0.477 | 0.011 | 0.483 | 0.214 | 0.273 | 0.765 | 0.338 | 0.079 | 0.905 | -0.205 | 0.295 | 0.864 | 0.001 | 0.999 | 0.999 | 0.203 | 0.299 | 0.927 |
| **C6** | 0.018 | 0.928 | 0.992 | 0.304 | 0.115 | 0.709 | 0.129 | 0.511 | 0.977 | 0.302 | 0.118 | 0.627 | -0.033 | 0.866 | 0.985 | -0.148 | 0.450 | 0.927 |
| **P5** | -0.090 | 0.647 | 0.992 | -0.043 | 0.829 | 0.929 | -0.033 | 0.866 | 0.977 | 0.177 | 0.365 | 0.879 | 0.010 | 0.959 | 0.993 | 0.159 | 0.417 | 0.927 |
| **P6** | 0.042 | 0.833 | 0.992 | 0.023 | 0.908 | 0.929 | 0.035 | 0.860 | 0.977 | 0.369 | 0.054 | 0.471 | -0.022 | 0.910 | 0.993 | -0.135 | 0.493 | 0.927 |
| **Fpz** | 0.203 | 0.300 | 0.992 | -0.059 | 0.765 | 0.929 | 0.072 | 0.716 | 0.977 | -0.530 | 0.004 | 0.125 | -0.222 | 0.255 | 0.973 | -0.109 | 0.580 | 0.927 |
| **CPz** | 0.256 | 0.188 | 0.992 | -0.181 | 0.356 | 0.765 | -0.232 | 0.234 | 0.935 | -0.105 | 0.595 | 0.933 | 0.088 | 0.657 | 0.973 | 0.432 | 0.022 | 0.330 |
| **POz** | -0.200 | 0.307 | 0.992 | -0.299 | 0.122 | 0.709 | 0.146 | 0.456 | 0.977 | 0.206 | 0.291 | 0.864 | 0.127 | 0.518 | 0.973 | 0.147 | 0.453 | 0.927 |
| **FCz** | 0.111 | 0.572 | 0.992 | -0.042 | 0.833 | 0.929 | -0.020 | 0.919 | 0.997 | 0.066 | 0.739 | 0.945 | 0.228 | 0.242 | 0.973 | -0.022 | 0.912 | 0.956 |

**Supplementary Table 24. Correlations among heartbeat-evoked potential, heartbeat-related power, and heartbeat-related inter-trial coherence, computed separately for inhalation and exhalation phases, across frequency bands and experimental tasks.** Abbreviations: HCT = heartbeat counting task, C-TCT = cardiac-tone counting task, SE = standard error, FDR = false discovery rate.

|  |  | | | **HCT** | | |  | | |  | | |
| --- | --- | --- | --- | --- | --- | --- | --- | --- | --- | --- | --- | --- |
|  | **Theta** | | | **Alpha** | | | **Beta** | | | **Gamma** | | |
| **Msec** | **rho** | **p** | **p_FDR_** | **rho** | **p** | **p_FDR_** | **rho** | **p** | **p_FDR_** | **rho** | **p** | **p_FDR_** |
| **390** | -0.021 | 0.917 | 0.994 | 0.185 | 0.344 | 0.344 | 0.046 | 0.816 | 0.816 | -0.372 | 0.052 | 0.346 |
| **401** | -0.013 | 0.950 | 0.994 | 0.245 | 0.208 | 0.219 | 0.083 | 0.675 | 0.710 | -0.461 | 0.014 | 0.286 |
| **412** | 0.008 | 0.970 | 0.994 | 0.330 | 0.087 | 0.097 | 0.115 | 0.557 | 0.619 | -0.414 | 0.029 | 0.293 |
| **422** | 0.002 | 0.994 | 0.994 | 0.413 | 0.030 | **0.035** | 0.206 | 0.292 | 0.443 | -0.292 | 0.131 | 0.437 |
| **434** | 0.028 | 0.886 | 0.994 | 0.473 | 0.012 | **0.016** | 0.215 | 0.272 | 0.443 | -0.203 | 0.300 | 0.545 |
| **445** | 0.045 | 0.818 | 0.994 | 0.487 | 0.009 | **0.015** | 0.186 | 0.341 | 0.443 | -0.258 | 0.185 | 0.528 |
| **456** | 0.053 | 0.788 | 0.994 | 0.469 | 0.013 | **0.016** | 0.170 | 0.385 | 0.453 | -0.311 | 0.108 | 0.430 |
| **467** | 0.074 | 0.708 | 0.994 | 0.484 | 0.010 | **0.015** | 0.194 | 0.320 | 0.443 | -0.322 | 0.095 | 0.430 |
| **478** | 0.103 | 0.599 | 0.994 | 0.516 | 0.006 | **0.011** | 0.213 | 0.274 | 0.443 | -0.205 | 0.293 | 0.545 |
| **490** | 0.103 | 0.599 | 0.994 | 0.513 | 0.006 | **0.011** | 0.196 | 0.316 | 0.443 | -0.107 | 0.587 | 0.839 |
| **501** | 0.095 | 0.629 | 0.994 | 0.471 | 0.012 | **0.016** | 0.181 | 0.355 | 0.443 | -0.021 | 0.917 | 0.972 |
| **512** | 0.163 | 0.407 | 0.994 | 0.529 | 0.004 | **0.010** | 0.267 | 0.169 | 0.375 | 0.021 | 0.917 | 0.972 |
| **523** | 0.176 | 0.370 | 0.994 | 0.545 | 0.003 | **0.008** | 0.308 | 0.111 | 0.279 | 0.015 | 0.939 | 0.972 |
| **534** | 0.175 | 0.373 | 0.994 | 0.541 | 0.003 | **0.008** | 0.364 | 0.058 | 0.279 | 0.007 | 0.972 | 0.972 |
| **545** | 0.141 | 0.472 | 0.994 | 0.579 | 0.002 | **0.005** | 0.413 | 0.030 | 0.279 | -0.076 | 0.700 | 0.933 |
| **556** | 0.152 | 0.438 | 0.994 | 0.607 | 0.001 | **0.003** | 0.404 | 0.034 | 0.279 | -0.239 | 0.220 | 0.545 |
| **567** | 0.139 | 0.479 | 0.994 | 0.605 | 0.001 | **0.003** | 0.346 | 0.071 | 0.279 | -0.222 | 0.255 | 0.545 |
| **578** | 0.216 | 0.268 | 0.994 | 0.628 | 0.000 | **0.003** | 0.319 | 0.098 | 0.279 | -0.182 | 0.353 | 0.589 |
| **589** | 0.205 | 0.293 | 0.994 | 0.660 | 0.000 | **0.003** | 0.332 | 0.085 | 0.279 | -0.062 | 0.754 | 0.942 |
| **600** | 0.221 | 0.258 | 0.994 | 0.640 | 0.000 | **0.003** | 0.333 | 0.084 | 0.279 | 0.119 | 0.546 | 0.839 |

**Supplementary Table 25. Instantaneous correlations between respiratory phase-related differences (Δ = exhalation − inhalation) in heartbeat-evoked potential and heartbeat-related power across frequency bands during the heartbeat-detection task.** Abbreviations: HCT = heartbeat-counting task, Msec = milliseconds, FDR = false discovery rate.

|  |  | | | **C-TCT** | | |  | | |  | | |
| --- | --- | --- | --- | --- | --- | --- | --- | --- | --- | --- | --- | --- |
|  | **Theta** | | | **Alpha** | | | **Beta** | | | **Gamma** | | |
| **Msec** | **rho** | **p** | **p_FDR_** | **rho** | **p** | **p_FDR_** | **rho** | **p** | **p_FDR_** | **rho** | **p** | **p_FDR_** |
| **390** | 0.059 | 0.765 | 0.765 | -0.017 | 0.932 | 0.941 | 0.155 | 0.430 | 0.475 | -0.018 | 0.930 | 0.957 |
| **401** | 0.068 | 0.731 | 0.765 | 0.015 | 0.941 | 0.941 | 0.181 | 0.356 | 0.468 | 0.084 | 0.671 | 0.931 |
| **412** | 0.143 | 0.465 | 0.565 | 0.089 | 0.651 | 0.941 | 0.288 | 0.137 | 0.305 | -0.041 | 0.838 | 0.931 |
| **422** | 0.128 | 0.516 | 0.574 | 0.094 | 0.634 | 0.941 | 0.306 | 0.113 | 0.284 | -0.117 | 0.553 | 0.931 |
| **434** | 0.182 | 0.353 | 0.565 | 0.089 | 0.653 | 0.941 | 0.344 | 0.073 | 0.284 | -0.163 | 0.405 | 0.931 |
| **445** | 0.158 | 0.420 | 0.565 | 0.060 | 0.760 | 0.941 | 0.318 | 0.099 | 0.284 | -0.197 | 0.313 | 0.931 |
| **456** | 0.209 | 0.286 | 0.565 | 0.034 | 0.864 | 0.941 | 0.343 | 0.075 | 0.284 | -0.132 | 0.502 | 0.931 |
| **467** | 0.204 | 0.297 | 0.565 | 0.033 | 0.866 | 0.941 | 0.411 | 0.031 | 0.284 | -0.011 | 0.957 | 0.957 |
| **478** | 0.201 | 0.303 | 0.565 | 0.032 | 0.871 | 0.941 | 0.350 | 0.068 | 0.284 | 0.050 | 0.799 | 0.931 |
| **490** | 0.207 | 0.289 | 0.565 | 0.036 | 0.855 | 0.941 | 0.353 | 0.066 | 0.284 | -0.065 | 0.741 | 0.931 |
| **501** | 0.188 | 0.336 | 0.565 | -0.024 | 0.906 | 0.941 | 0.306 | 0.113 | 0.284 | -0.113 | 0.564 | 0.931 |
| **512** | 0.165 | 0.400 | 0.565 | -0.033 | 0.868 | 0.941 | 0.251 | 0.197 | 0.391 | -0.053 | 0.788 | 0.931 |
| **523** | 0.145 | 0.460 | 0.565 | -0.052 | 0.792 | 0.941 | 0.241 | 0.215 | 0.391 | 0.086 | 0.663 | 0.931 |
| **534** | 0.138 | 0.481 | 0.565 | -0.036 | 0.857 | 0.941 | 0.188 | 0.336 | 0.468 | 0.235 | 0.227 | 0.931 |
| **545** | 0.178 | 0.362 | 0.565 | -0.054 | 0.784 | 0.941 | 0.188 | 0.337 | 0.468 | 0.211 | 0.280 | 0.931 |
| **556** | 0.165 | 0.399 | 0.565 | -0.026 | 0.897 | 0.941 | 0.163 | 0.405 | 0.475 | 0.279 | 0.150 | 0.931 |
| **567** | 0.205 | 0.293 | 0.565 | -0.018 | 0.930 | 0.941 | 0.180 | 0.359 | 0.468 | 0.223 | 0.252 | 0.931 |
| **578** | 0.212 | 0.277 | 0.565 | 0.060 | 0.760 | 0.941 | 0.174 | 0.374 | 0.468 | 0.138 | 0.481 | 0.931 |
| **589** | 0.211 | 0.279 | 0.565 | 0.090 | 0.647 | 0.941 | 0.148 | 0.451 | 0.475 | 0.113 | 0.566 | 0.931 |
| **600** | 0.186 | 0.343 | 0.565 | 0.105 | 0.593 | 0.941 | 0.080 | 0.685 | 0.685 | 0.240 | 0.218 | 0.931 |

**Supplementary Table 26. Instantaneous correlations between respiratory phase-related differences (Δ = exhalation − inhalation) in heartbeat-evoked potential and heartbeat-related power across frequency bands during the cardiac-tone counting task.** Abbreviations: C-TCT = cardiac-tone counting task, Msec = milliseconds, FDR = false discovery rate.

|  | **Alpha** | | |
| --- | --- | --- | --- |
| **Msec** | **z** | **p** | **p_FDR_** |
| **390** | 0.7217 | 0.2352 | 0.2352 |
| **401** | 0.8327 | 0.2025 | 0.2132 |
| **412** | 0.8938 | 0.1857 | 0.2063 |
| **422** | 1.2221 | 0.1108 | 0.1304 |
| **434** | 1.5048 | 0.0662 | 0.0827 |
| **445** | 1.6663 | 0.0478 | 0.0638 |
| **456** | 1.6791 | 0.0466 | 0.0638 |
| **467** | 1.7512 | 0.0400 | 0.0615 |
| **478** | 1.9022 | 0.0286 | 0.0506 |
| **490** | 1.8755 | 0.0304 | 0.0506 |
| **501** | 1.8899 | 0.0294 | 0.0506 |
| **512** | 2.1964 | 0.0140 | **0.0312** |
| **523** | 2.3431 | 0.0096 | **0.0291** |
| **534** | 2.2685 | 0.0117 | **0.0291** |
| **545** | 2.5262 | 0.0058 | **0.0291** |
| **556** | 2.5806 | 0.0049 | **0.0291** |
| **567** | 2.5393 | 0.0056 | **0.0291** |
| **578** | 2.3953 | 0.0083 | **0.0291** |
| **589** | 2.4835 | 0.0065 | **0.0291** |
| **600** | 2.3067 | 0.0105 | **0.0291** |

**Supplementary Table 27. Instantaneous correlations between respiratory phase-related differences (Δ = exhalation − inhalation) in the heartbeat-evoked potential and alpha-band heartbeat-related power, compared across the heartbeat-counting task and the cardiac-tone counting tasks.** Abbreviations: Msec = milliseconds, FDR = false discovery rate.

| **N** | **R-notation formula** | **AIC** |
| --- | --- | --- |
| 1 | ΔHEP ∼ 1 + Δalpha power + (1 \| ID) | 880 |
| 2 | ΔHEP ∼ 1 + Δalpha ITC + (1 \| ID) | 879 |
| 3 | ΔHEP ∼ 1 + ΔECG + (1 \| ID) | 880 |
| 4 | ΔHEP ∼ 1 + ΔHR + (1 \| ID) | 880 |
| 5 | ΔHEP ∼ 1 + Δalpha power + Δalpha ITC + (1 \| ID) | 881 |
| **6** | **ΔHEP ∼ 1 + Δalpha power * Δalpha ITC + (1 \| ID)** | **878** |
| 7 | ΔHEP ∼ 1 + Δalpha power * Δalpha ITC + ΔHR (1 \| ID) | 880 |
| 8 | ΔHEP ∼ 1 + Δalpha power * Δalpha ITC + ΔHR * Δalpha power + (1 \| ID) | 882 |
| 9 | ΔHEP ∼ 1 + Δalpha power * Δalpha ITC + ΔHR * Δalpha ITC + (1 \| ID) | 882 |
| 10 | ΔHEP ∼ 1 + Δalpha power * Δalpha ITC * ΔHR + (1 \| ID) | 886 |
| 11 | ΔHEP ∼ 1 + Δalpha power * Δalpha ITC + ΔECG (1 \| ID) | 880 |
| 12 | ΔHEP ∼ 1 + Δalpha power * Δalpha ITC + ΔECG * Δalpha power + (1 \| ID) | 882 |
| 13 | ΔHEP ∼ 1 + Δalpha power * Δalpha ITC + ΔECG * Δalpha ITC + (1 \| ID) | 882 |
| 14 | ΔHEP ∼ 1 + Δalpha power * Δalpha ITC * ΔECG + (1 \| ID) | 883 |

**Supplementary Table 28. Comparison of linear mixed-effects models for predicting Δheartbeat-evoked potential during the heartbeat counting task.** Abbreviations: N = number, ID = participant, HEP = heartbeat-evoked potential, ITC = inter-trial coherence, HR = heart rate, ECG = electrocardiogram, AIC = Akaike information criterion. Note: the simplest model with the lowest AIC value is considered the best fitting (model 6, bold).

|  | **Estimate** | **SE** | **t** | **p** |
| --- | --- | --- | --- | --- |
| **Intercept** | 0.00311 | 0.0613 | 0.0508 | 0.9599 |
| **Δalpha power** | 0.02741 | 0.0572 | 0.4792 | 0.6322 |
| **Δalpha ITC** | 0.06196 | 0.0569 | 1.0891 | 0.2770 |
| **Δalpha power * Δalpha ITC** | 0.11828 | 0.0564 | 2.0958 | 0.0369 |

**Supplementary Table 29. Parameter estimates of the best-fitting linear mixed-effects model for predicting Δheartbeat-evoked potential during the heartbeat counting task.** Abbreviations: ITC = inter-trial coherence, SE = standard error.

| **Moderator** |  |  |  |  |  |
| --- | --- | --- | --- | --- | --- |
| **Δalpha power** | **Effect** | **Estimate** | **SE** | **t** | **p** |
| **Mean-1SD** | **Δalpha ITC** | -0.0563 | 0.0821 | -0.686 | 0.4931 |
| **Mean** | **Δalpha ITC** | 0.0620 | 0.0569 | 1.089 | 0.2770 |
| **Mean+1SD** | **Δalpha ITC** | 0.1802 | 0.0782 | 2.306 | **0.0218** |

**Supplementary Table 30. Parameter estimates for simple-effects of Δalpha inter-trial coherence during the heartbeat counting task.** Abbreviations: ITC = inter-trial coherence, SE = standard error, SD = standard deviation.

| **N** | **R-notation formula** | **AIC** |
| --- | --- | --- |
| 1 | Accuracy ∼ 1 + ΔHEP + (1 \| ID) | 590 |
| 2 | Accuracy ∼ 1 + Δalpha power + (1 \| ID) | 585 |
| 3 | Accuracy ∼ 1 + Δalpha ITC + (1 \| ID) | 591 |
| 4 | Accuracy ∼ 1 + ΔECG + (1 \| ID) | 591 |
| 5 | Accuracy ∼ 1 + ΔHR + (1 \| ID) | 591 |
| 6 | Accuracy ∼ 1 + ΔHEP + Δalpha power + (1 \| ID) | 586 |
| 7 | Accuracy ∼ 1 + ΔHEP*Δalpha power + (1 \| ID) | 584 |
| 8 | Accuracy ∼ 1 + confidence + (1 \| ID) | 542 |
| 9 | Accuracy ∼ 1 + confidence + (1 + confidence \| ID) | 527 |
| 10 | Accuracy ∼ 1 + ΔHEP*Δalpha power + confidence + (1 + confidence \| ID) | 524 |
| **11** | **Accuracy ∼ 1 + ΔHEP*Δalpha power*confidence + (1 + confidence \| ID)** | **523** |
| 12 | Accuracy ∼ 1 + ΔHEP*Δalpha power*confidence + ΔECG + (1 + confidence \| ID) | 525 |
| 13 | Accuracy ∼ 1 + ΔHEP*Δalpha power*confidence + ΔECG*ΔHEP + (1 + confidence \| ID) | 527 |
| 14 | Accuracy ∼ 1 + ΔHEP*Δalpha power*confidence + ΔECG*Δalpha power + (1 + confidence \| ID) | 527 |
| 15 | Accuracy ∼ 1 + ΔHEP*Δalpha power*confidence + ΔECG*confidence + (1 + confidence \| ID) | 523 |
| 16 | Accuracy ∼ 1 + ΔHEP*Δalpha power*confidence*ΔECG + (1 + confidence \| ID) | 532 |
| 17 | Accuracy ∼ 1 + ΔHEP*Δalpha power*confidence + ΔHR + (1 + confidence \| ID) | 525 |
| 18 | Accuracy ∼ 1 + ΔHEP*Δalpha power*confidence + ΔHR*ΔHEP + (1 + confidence \| ID) | 527 |
| 19 | Accuracy ∼ 1 + ΔHEP*Δalpha power*confidence + ΔHR*Δalpha power + (1 + confidence \| ID) | 527 |
| 20 | Accuracy ∼ 1 + ΔHEP*Δalpha power*confidence + ΔHR*confidence + (1 + confidence \| ID) | 523 |
| 21 | Accuracy ∼ 1 + ΔHEP*Δalpha power*confidence*ΔHR + (1 + confidence \| ID) | 533 |

**Supplementary Table 31. Comparison of linear mixed-effects models for predicting accuracy during the heartbeat counting task.** Abbreviations: N = number, ID = participant, HEP = heartbeat-evoked potential, ITC = inter-trial coherence, HR = heart rate, ECG = electrocardiogram, AIC = Akaike information criterion. Note: the simplest model with the lowest AIC value is considered the best fitting (model 11, bold).

|  | **Estimate** | **SE** | **t** | **p** |
| --- | --- | --- | --- | --- |
| **Intercept** | -0.01155 | 0.1521 | -0.0759 | 0.9401 |
| **Δalpha power** | -0.06146 | 0.0299 | -2.0579 | 0.0406 |
| **ΔHEP** | 0.03327 | 0.0281 | 1.1838 | 0.2376 |
| **Confidence** | 0.33154 | 0.0822 | 4.0314 | 0.0006 |
| **Δalpha power * ΔHEP** | 0.02733 | 0.0247 | 1.1070 | 0.2693 |
| **Δalpha power * confidence** | -0.02232 | 0.0325 | -0.6864 | 0.4931 |
| **ΔHEP * confidence** | -0.00121 | 0.0290 | -0.0418 | 0.9667 |
| **Δalpha power * ΔHEP * confidence** | 0.06790 | 0.0300 | 2.2636 | 0.0245 |

**Supplementary Table 32. Parameter estimates of the best-fitting linear mixed-effects model for predicting accuracy during the heartbeat counting task.** Abbreviations: ITC = inter-trial coherence, SE = standard error.

| **Moderator** | | **Effect** | **Estimate** | **SE** | **t** | **p** |
| --- | --- | --- | --- | --- | --- | --- |
| **Confidence** | **Δalpha power** |  |  |  |  |  |
| **Mean-1SD** | **Mean-1SD** | **ΔHEP** | 0.07505 | 0.0675 | 1.112 | 0.26743 |
|  | **Mean** | **ΔHEP** | 0.03448 | 0.0416 | 0.829 | 0.40805 |
|  | **Mean+1SD** | **ΔHEP** | -0.00608 | 0.0503 | -0.121 | 0.90378 |
| **Mean** | **Mean-1SD** | **ΔHEP** | 0.00594 | 0.0398 | 0.149 | 0.88157 |
|  | **Mean** | **ΔHEP** | 0.03327 | 0.0281 | 1.184 | 0.23761 |
|  | **Mean+1SD** | **ΔHEP** | 0.06060 | 0.0348 | 1.740 | 0.08303 |
| **Mean+1SD** | **Mean-1SD** | **ΔHEP** | -0.06317 | 0.0560 | -1.129 | 0.26018 |
|  | **Mean** | **ΔHEP** | 0.03206 | 0.0391 | 0.820 | 0.41312 |
|  | **Mean+1SD** | **ΔHEP** | 0.12729 | 0.0483 | 2.633 | **0.00900** |

**Supplementary Table 33. Parameter estimates for simple-effects of ΔHEP during the heartbeat counting task.** Abbreviations: HEP = heartbeat-evoked potential, ITC = inter-trial coherence, SE = standard error, SD = standard deviation.

| **N** | **R-notation formula** | **AIC** |
| --- | --- | --- |
| 1 | Accuracy ∼ 1 + ΔHEP + (1 \| ID) | 861 |
| 2 | Accuracy ∼ 1 + Δalpha power + (1 \| ID) | 865 |
| 3 | Accuracy ∼ 1 + Δalpha ITC + (1 \| ID) | 865 |
| 4 | Accuracy ∼ 1 + ΔECG + (1 \| ID) | 864 |
| 5 | Accuracy ∼ 1 + ΔHR + (1 \| ID) | 863 |
| 6 | Accuracy ∼ 1 + ΔHEP + Δalpha power + (1 \| ID) | 863 |
| 7 | Accuracy ∼ 1 + ΔHEP * Δalpha power + (1 \| ID) | 864 |
| 8 | Accuracy ∼ 1 + confidence + (1 \| ID) | 836 |
| **9** | **Accuracy ∼ 1 + confidence + (1 + confidence \| ID)** | **798** |
| 10 | Accuracy ∼ 1 + ΔHEP*Δalpha power + confidence + (1 + confidence \| ID) | 800 |
| 11 | Accuracy ∼ 1 + ΔHEP*Δalpha power*confidence + (1 + confidence \| ID) | 802 |
| 12 | Accuracy ∼ 1 + ΔHEP*Δalpha power*confidence + ΔECG + (1 + confidence \| ID) | 803 |
| 13 | Accuracy ∼ 1 + ΔHEP*Δalpha power*confidence + ΔECG*ΔHEP + (1 + confidence \| ID) | 804 |
| 14 | Accuracy ∼ 1 + ΔHEP*Δalpha power*confidence + ΔECG*Δalpha power + (1 + confidence \| ID) | 805 |
| 15 | Accuracy ∼ 1 + ΔHEP*Δalpha power*confidence + ΔECG*confidence + (1 + confidence \| ID) | 802 |
| 16 | Accuracy ∼ 1 + ΔHEP*Δalpha power*confidence*ΔECG + (1 + confidence \| ID) | 798 |
| 17 | Accuracy ∼ 1 + ΔHEP*Δalpha power*confidence + ΔHR + (1 + confidence \| ID) | 803 |
| 18 | Accuracy ∼ 1 + ΔHEP*Δalpha power*confidence + ΔHR*ΔHEP + (1 + confidence \| ID) | 803 |
| 19 | Accuracy ∼ 1 + ΔHEP*Δalpha power*confidence + ΔHR*Δalpha power + (1 + confidence \| ID) | 802 |
| 20 | Accuracy ∼ 1 + ΔHEP*Δalpha power*confidence + ΔHR*confidence + (1 + confidence \| ID) | 805 |
| 21 | Accuracy ∼ 1 + ΔHEP*Δalpha power*confidence*ΔHR + (1 + confidence \| ID) | 802 |

**Supplementary Table 34. Comparison of linear mixed-effects models for predicting accuracy during the cardiac-tone counting task.** Abbreviations: N = number, ID = participant, HEP = heartbeat-evoked potential, ITC = inter-trial coherence, HR = heart rate, ECG = electrocardiogram, AIC = Akaike information criterion. Note: the simplest model with the lowest AIC value is considered the best fitting (model 9, bold).

|  | **Estimate** | **SE** | **t** | **p** |
| --- | --- | --- | --- | --- |
| **Intercept** | -0.0117 | 0.0683 | -0.172 | 0.8646 |
| **Confidence** | 0.3779 | 0.1101 | 3.433 | 0.0018 |

**Supplementary Table 35. Parameter estimates of the best-fitting linear mixed-effects model for predicting accuracy during the cardiac-tone counting task.** Abbreviations: SE = standard error.
